# GSK-3 inhibitor elraglusib enhances tumor-infiltrating immune cell activation in tumor biopsies and synergizes with anti-PD-L1 in a murine model of colorectal cancer

**DOI:** 10.1101/2023.02.07.527499

**Authors:** Kelsey E. Huntington, Anna D. Louie, Praveen R. Srinivasan, Christoph Schorl, Shaolei Lu, David Silverberg, Daniel Newhouse, Zhijin Wu, Lanlan Zhou, Brittany A. Borden, Francis J. Giles, Mark Dooner, Benedito A. Carneiro, Wafik S. El-Deiry

**Author notes:** Correspondence; 70 Ship Street, Box G-E5, Providence, RI. Contacts for reagents and resource sharing: Requests for information and reagents should be directed to the corresponding author.

## Abstract

Inhibition of GSK-3 using small-molecule elraglusib has shown promising preclinical antitumor activity. Using in vitro systems, we found that elraglusib promotes immune cell-mediated tumor cell killing, enhances tumor cell pyroptosis, decreases tumor cell NF-κB-regulated survival protein expression, and increases immune cell effector molecule secretion. Using in vivo systems, we observed synergy between elraglusib and anti-PD-L1 in an immunocompetent murine model of colorectal cancer. Murine responders had more tumor-infiltrating T-cells, fewer tumor-infiltrating Tregs, lower tumorigenic circulating cytokine concentrations, and higher immunostimulatory circulating cytokine concentrations. To determine the clinical significance, we utilized human plasma samples from patients treated with elraglusib and correlated cytokine profiles with survival. Using paired tumor biopsies, we found that CD45+ tumor-infiltrating immune cells had lower expression of inhibitory immune checkpoints and higher expression of T-cell activation markers in post-elraglusib patient biopsies. These results introduce several immunomodulatory mechanisms of GSK-3 inhibition using elraglusib, providing a rationale for the clinical evaluation of elraglusib in combination with immunotherapy.

**Statement of significance:** Pharmacologic inhibition of GSK-3 using elraglusib sensitizes tumor cells, activates immune cells for increased anti-tumor immunity, and synergizes with anti-PD-L1 immune checkpoint blockade. These results introduce novel biomarkers for correlations with response to therapy which could provide significant clinical utility and suggest that elraglusib, and other GSK-3 inhibitors, should be evaluated in combination with immune checkpoint blockade.

## Introduction

Glycogen synthase kinase 3 (GSK-3) is a serine/threonine kinase with key roles in myriad biological processes such as tumor progression, and inhibition of GSK-3 using a novel small-molecule elraglusib has shown promising preclinical antitumor activity in multiple tumor types (1). There is a growing body of literature characterizing the immunomodulatory roles of GSK-3 in the context of anti-tumor immunity (2). GSK-3 is known to inhibit cytokine production and T cell activation (3,4). Aberrant overexpression of GSK-3 has been shown to promote tumor growth and epithelial-to-mesenchymal transition (EMT) through various mechanisms including modulation of pro-survival NF-κB signaling pathways (5). Thus, GSK-3 is a promising target in the treatment of human malignancies.

Globally, colorectal cancer (CRC) ranks third in terms of incidence and second in terms of mortality. Treatment options include surgery, chemotherapy, radiation therapy, targeted therapy, and immunotherapy. Immune checkpoint blockade (ICB) has now entered into clinical care for CRC with the recent U.S. Food and Drug Administration approvals of checkpoint inhibitors nivolumab and pembrolizumab for microsatellite instability-high (MSI-H) CRC cases after chemotherapy (6). Thus far, ICB clinical trials have demonstrated efficacy in MSI-H CRC, however, the impressive durability of tumor regression stands in stark contrast with the lack of response observed in microsatellite stable (MSS) CRC (6). Thus, there remains a substantial unmet need in the ∼85% of patients with MSS CRC in whom ICB is less effective (7). Moreover, the percentage of patients with MSS CRC dramatically increases to ∼96% in Stage IV disease (7).

We sought to evaluate elraglusib (9-ING-41), a small molecule that targets GSK-3 which has the potential to increase the efficacy of ICB. We chose to evaluate elraglusib, which inhibits both α and β isoforms, because it is a clinically relevant small molecule with superior pharmacokinetic properties and is significantly more potent than other GSK-3 inhibitors (8,9). Although there are ongoing efforts to further characterize the immunomodulatory effects of GSK-3 inhibitors, few utilize small-molecule elraglusib (10-12).

Here, we characterize the effects of elraglusib in vitro on tumor and immune cells, in vivo in combination with ICB in a syngeneic murine colon carcinoma BALB/c model using MSS cell line CT-26, and in human tumor biopsies and plasma samples from patients with refractory solid tumors of multiple tissue origins enrolled in a Phase 1 clinical trial investigating elraglusib (<u>NCT03678883</u>).

## Results

### Elraglusib sensitizes tumor cells to immune-mediated cytotoxicity

A co-culture of fluorescently-labeled SW480 MSS CRC cells and TALL-104 CD8+ T cells treated with elraglusib led to an increase in tumor cell death after 24 hours (Supplementary Figure 1A). Treatment doses chosen were significantly less than the 24-and 72-hour IC-50s calculated for all cell lines evaluated in the co-culture to ensure the majority of tumor cell death was immune cell-mediated (Supplementary Figure 2A-B). We observed limited tumor cell death in SW480 monocultures treated with drug only (Supplementary Figure 1B). In the co-culture with tumor and immune cells only, in the absence of the drug, we noted that the baseline percentage of dead cells out of total cells was approximately 40%, after normalization. Co-cultures of tumor cells and TALL-104 T cells treated with 5 µM elraglusib had an average of 60% dead cells, while co-cultures treated with 10 µM of elraglusib had an average of 65% dead cells (Supplementary Figure 1C).

Because TALL-104 cells are a human leukemic T cell line, we next wanted to determine the relevancy of these results using normal T cells. Donor-derived CD8+ T cells were isolated from a donor blood sample in accordance with an IRB-approved protocol. A co-culture of fluorescently-labeled SW480 tumor cells and CD8+ donor-derived CD8+ T cells was then treated with elraglusib and the percentage of dead cells out of total cells was quantified after 24 hours (Supplementary Figure 1D). We again observed limited tumor cell death in SW480 monocultures treated with drug only (Supplementary Figure 1E). The data was then normalized, as previously described, and we noted even more robust immune cell-mediated tumor cell death in the co-cultures treated with elraglusib (Supplementary Figure 1F). Co-cultures of tumor cells and donor-derived CD8+ T cells treated with 5 µM elraglusib had an average of 65% dead cells, while co-cultures treated with 10 µM of elraglusib had an average of 75% dead cells.

To determine if the increased amount of immune-cell mediated tumor cell-killing was due to the drug’s impact on the tumor cells or the immune cells, we next pre-treated tumor cells with elraglusib for 24 hours before the co-culture with immune cells began. We observed that pre-treatment with elraglusib sensitized SW480 tumor cells to TALL-104 cell-mediated tumor cell killing (Figure 1A). We again used the raw percentages of cell death to normalize the data and observed minimal amounts of drug cytotoxicity at the concentration and duration of treatment used (Figure 1B). Tumor cells pre-treated with 5 µM elraglusib for 24 hours and then co-cultured with TALL-104 T cells had an average of 65% dead cells (Figure 1C). Once again, we sought to confirm these co-culture results using donor-derived CD8+ T cells instead of TALL-104 cells (Figure 1D). We observed similar results with the CD8+ T cells where elraglusib pre-treatment of tumor cells led to a statistically significant increase in tumor cell death after 24 hours of co-culture (Figure 1E). Tumor cells pre-treated with 5 µM elraglusib for 24 hours and then co-cultured with donor-derived CD8+ T cells had an average of 65% dead cells, while co-cultures treated with 10 µM of elraglusib for 24 hours and then co-cultured with donor-derived CD8+ T cells had an average of 70% dead cells (Figure 1F).

**Figure 1.**
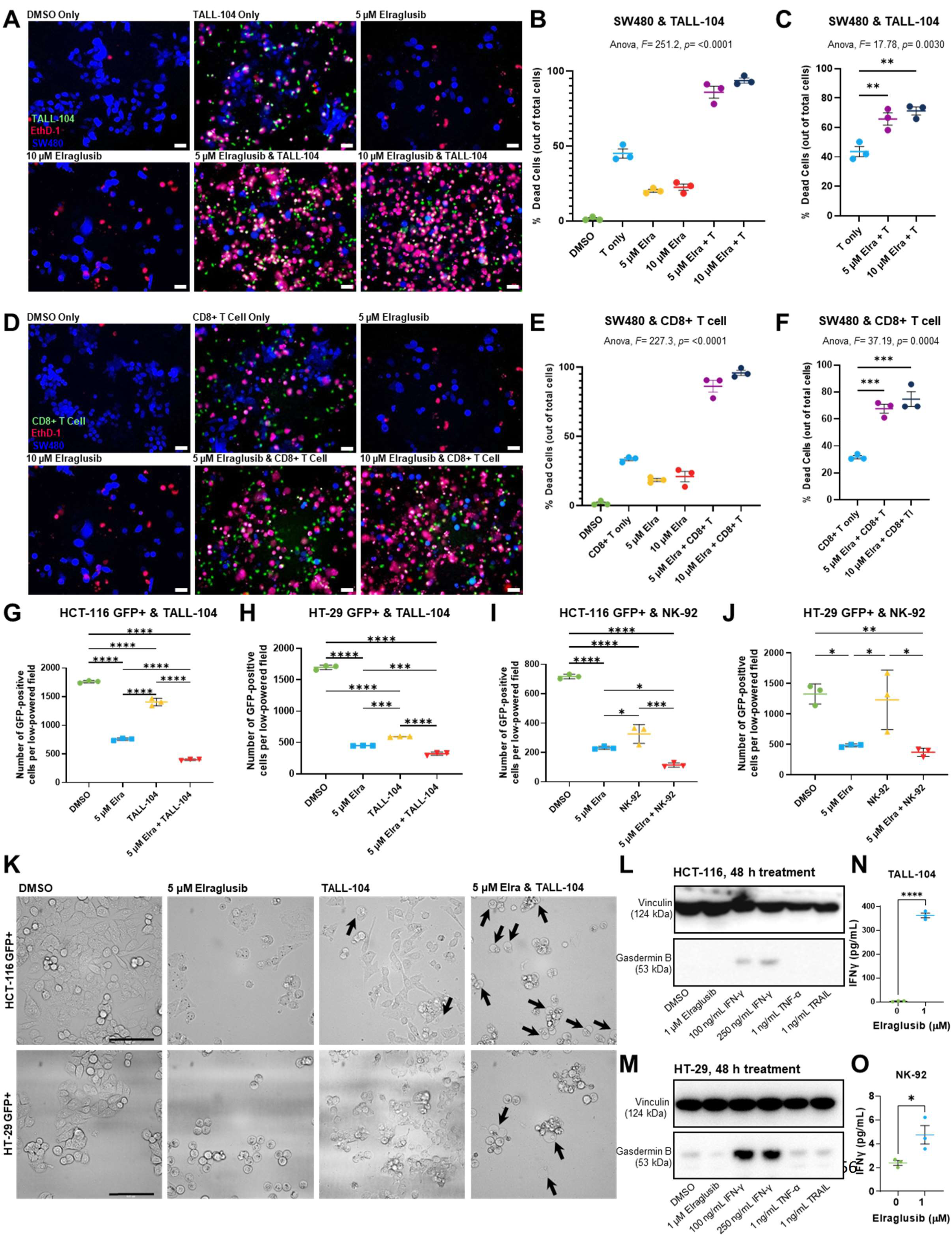
Elraglusib triggers pyroptosis and sensitizes tumor cells to increase immune-mediated cytotoxicity in a co-culture model. Co-cultures were treated with drug concentrations as indicated. A 1:1 effector to target (E:T) ratio was used with a 24-hour co-culture duration. EthD-1 was used to visualize dead cells, 10X magnification, scale bar indicates 100 µm. **(A)** SW480 and TALL-104 T cell co-culture assay images at the 24-hour timepoint. 24-hour tumor cell pre-treatment with 5 µM elraglusib, followed by 24-hour co-culture. **(B)** Quantification of co-culture experiment using the percentage of dead cells out of total cells (n=3). **(C)** Quantification normalized by cell death observed with drug treatment alone (n=3). **(D)** SW480 and donor-derived CD8+ T cell co-culture assay images at the 24-hour timepoint. 24-hour tumor cell pre-treatment with 5 µM elraglusib, followed by 24-hour co-culture. **(E)** Quantification of co-culture experiment using the percentage of dead cells out of total cells (n=3). **(F)** Quantification normalized by cell death observed with drug treatment alone (n=3). **(G)** The number of HCT 116 GFP+ cells were quantified after 48 hours of culture with DMSO, 5 μM elraglusib, and/or 5000 TALL-104 cells (n=3). **(H)** The number of HT-29 GFP+ cells were quantified after 48 hours of culture with DMSO, 5 μM elraglusib, and/or 5000 TALL-104 cells (n=3). **(I)** The number of HCT 116 GFP+ cells were quantified after 48 hours of culture with DMSO, 5 μM elraglusib, and/or 5000 NK-92 cells (n=3). **(J)** The number of HT-29 GFP+ cells were quantified after 48 hours of culture with DMSO, 5 μM elraglusib, and/or 5000 NK-92 cells (n=3). **(K)** 40X images were collected with a Molecular Devices ImageXpress® Confocal HT.ai High-Content Imaging System. White arrows indicate pyroptotic events. Western blot analysis of **(L)** HCT-116 and **(M)** HT-29 CRC cells for expression of indicated proteins after treatment with indicated cytokines or drugs. Quantification of IFN-γ secretion (pg/mL) post-DMSO or elraglusib treatment for 24 hours in **(N)** TALL-104 cells and **(O)** NK-92 cells (n=3). Error bars represent the mean +/-standard deviation. Statistical test: one-way ANOVA with Tukey’s test for multiple comparisons. P-value legend: * p < 0.05, ** p < 0.01, *** p < 0.001, **** p < 0.0001.

To confirm these results, we repeated these experiments using a GFP+ co-culture system with additional CRC cell lines HCT-116 and HT-29 (Supplementary Figure 1G). We chose to evaluate both HCT-116 and HT-29 CRC cells in this co-culture model to determine if the elraglusib-mediated increase in immune cell-mediated SW480 cell killing could be reproduced in additional CRC cell lines. These cell lines were selected based on their varied mutational profiles, with both MSI-H and MSS statuses reflected (Supplementary Figure 2C). When HCT-116 GFP+ cells were co-cultured with TALL-104 cells in the presence or absence of 5 µM elraglusib we noted a significant decrease in GFP+ cells per low-powered field in the 5 µM elraglusib only, TALL-104 only, and combination therapy groups as compared to the DMSO only control group (Figure 1G). We noted a significant decrease in the number of GFP+ cells per field in the combination therapy group of TALL-104 and 5 µM elraglusib co-culture condition as compared to TALL-104 which recapitulated the results observed in the first co-culture system. We observed a similar trend in the HT-29 cell line where the combination therapy group showed increased tumor cell death as compared to the drug-only or T cell-only groups (Figure 1H). To determine if these results applied to other cytotoxic immune cell lines we repeated the co-culture experiments with a natural killer cell line, NK-92. We observed similar trends in the co-culture of NK-92 cells with HCT-116 cells where the combination of 5 µM elraglusib and NK-92 cells showed increased tumor cell death as compared to the drug-only treatment or immune cell-only treatment (Figure 1I). In the HT-29 cells, we noted increased tumor cell death in the combination therapy group as compared to immune cell only and as compared to DMSO only (Figure 1J).

### Elraglusib enhances tumor cell pyroptosis in a co-culture of colorectal cancer cells and immune cells

To determine if pyroptosis-mediated immune cell activity played a role in the co-culture results observed, we examined higher-power co-culture images for evidence of pyroptosis. Indeed, we observed some pyroptotic events in the co-cultures involving tumor cells and TALL-104 cells only (Figure 1K). We did not observe any pyroptotic events in the DMSO or drug-only conditions suggesting that tumor cell pyroptosis was mediated by an immune cell-secreted molecule as it was only observed in the co-culture wells with immune cells (Figure 1K, Supplementary Figure 1H). Interestingly, in the co-culture of CRC (HCT-116, HT-29) and TALL-104 cells in the presence of 5 μM elraglusib treatment, we noted a significant increase in pyroptotic events (Figure 1K, Supplementary Figure 1I-J). To determine what immune cell-secreted molecules were most likely contributing to tumor cell pyroptosis, we probed for downstream mediator of pyroptotic death gasdermin B expression in tumor cells treated with a vehicle-only control (DMSO), 1 μM elraglusib, 100 mg/mL IFN-γ, 250 ng/mL IFN-γ, 1 ng/mL TNF-α, and 1 ng/mL TRAIL (Figure 1L). We observed an increase in gasdermin B expression with both concentrations of IFN-γ used in both the HCT-116 cells and the HT-29 cells suggesting that IFN-γ secreted by immune cells was a major contributor to the pyroptosis observed (Figure 1M). To test whether immune cells secrete more IFN-γ post-treatment with elraglusib, we treated immune cell lines (TALL-104, NK-92) with elraglusib for 24 hours and indeed noted a significant increase in IFN-γ post-treatment in cell culture supernatants (Figure 1N-O).

### Elraglusib upregulates tumor cell PD-L1 and proapoptotic pathway expression as well as downregulates immunosuppressive/angiogenic protein expression and pro-survival pathways

To help elucidate the mechanism behind the CRC cell sensitization to immune cell killing that we observed in the co-culture assays, we performed western blot analyses on CRC cells (HCT-116, HT-29) treated with elraglusib over a 72-hour timecourse. Using the same low dose of elraglusib utilized in the co-culture assays, we observed little to no cleaved PARP (cPARP) in both cell lines analyzed until the 48-hour timepoint confirming that the tumor cell death observed in the co-culture assays was not a product of drug cytotoxicity (Figure 2A). Because GSK-3 is a known regulator of NF-κB signaling pathways we also probed for NF-κB p65 and noted decreased expression as the timecourse progressed. However, we observed increases in PD-L1 expression as the treatment duration increased. To further elucidate elraglusib-mediated effects on tumor cell survival we probed for survival factors Bcl-2 and Survivin and noted decreases in protein expression in both cell lines analyzed, especially at the later timepoints (48, 72 hr). In HCT-116 cells, we also probed for survival factor Mcl-1 and again noted marked decreases in protein expression by the 24-hour timepoint (Figure 2B).

**Figure 2.**
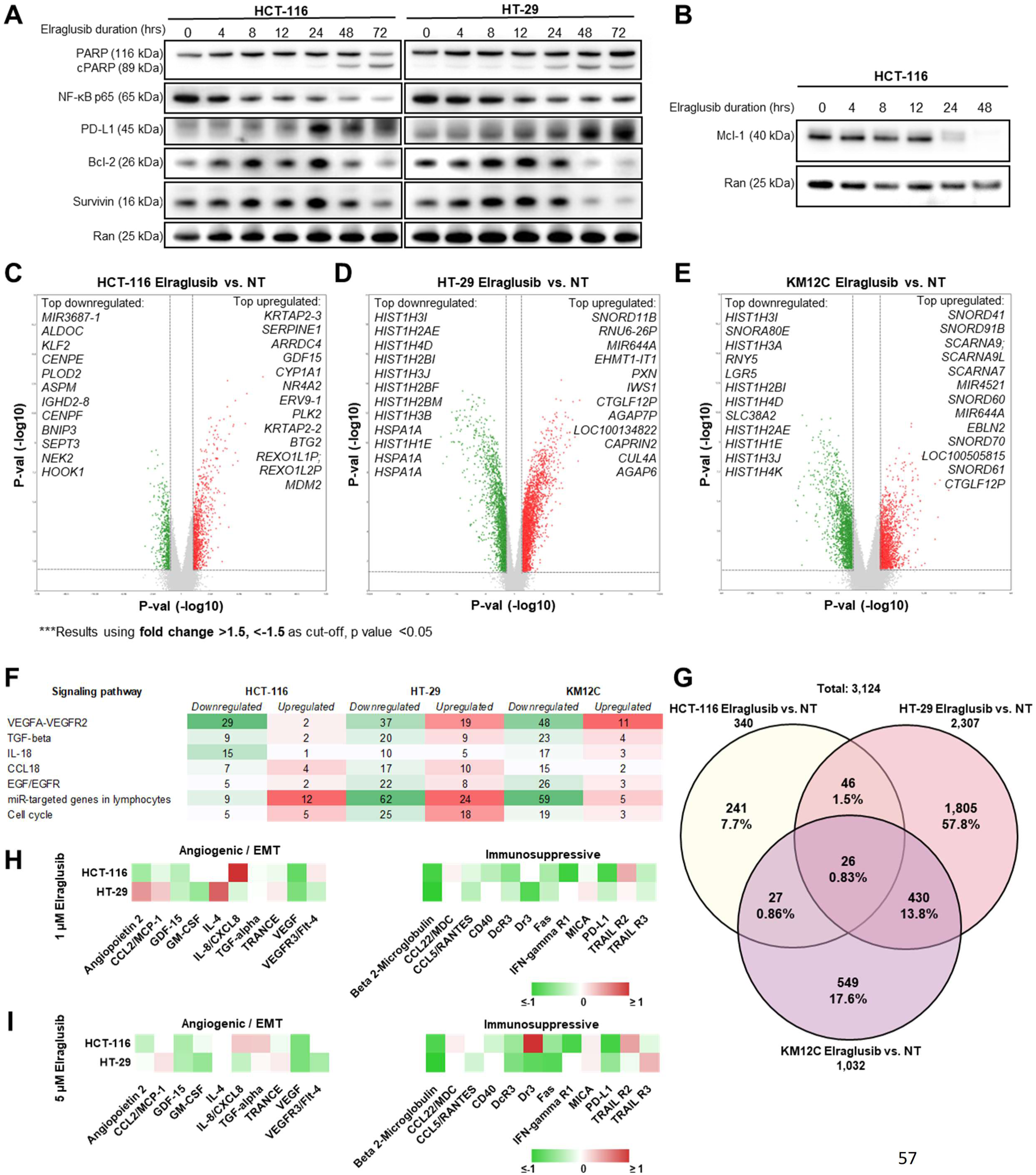
Elraglusib treatment induces apoptosis and suppresses survival pathways in tumor cells. Western blot analysis of **(A)** HCT-116 and HT-29 CRC cells for expression of indicated proteins after increasing durations of elraglusib treatment (0-72 hours). **(B)** Western blot analysis of Mcl-1 expression in HCT-116 CRC cells after increasing durations of elraglusib treatment. CRC cells were treated with 1 μM elraglusib for 24 hours and treated versus untreated samples were compared in triplicate. Microarray analysis results were visualized using volcano plots for **(C)** HCT-116, **(D)** HT-29, and **(E)** KM12C CRC cell lines. Top down-and up-regulated genes post-elraglusib treatment as compared to controls are shown. Results were calculated using a FC cutoff of >1.5, <-1.5, and a p value of <0.05. **(F)** The number of genes up-or down-regulated in each of the three cell lines within several signaling pathways of interest. Green indicates downregulation and red indicates upregulation of gene expression. **(G)** A Venn Diagram was used to compare the 3,124 genes that were differentially expressed post-treatment with elraglusib in the three colon cancer cell lines (HCT-116, HT-29, KM12C). **(H)** Tumor cells (HCT-116, HT-29) were treated with 1 μM elraglusib for 48 hours and cell culture supernatant was analyzed with the Luminex 200. Red indicates a positive FC and green indicates a negative FC (n=3). **(I)** Tumor cells (HCT-116, HT-29) were treated with 5 μM elraglusib for 48 hours and cell culture supernatant was analyzed with the Luminex 200. Red indicates a positive FC and green indicates a negative FC (n=3).

We then utilized microarray analysis to gain insights into gene expression changes in CRC cell lines post-GSK-3 inhibition with elraglusib. Several CRC cell lines (HCT-116, HT-29, KM12C) were treated with elraglusib at IC-50 concentrations or DMSO as vehicle control for 24 hours, and treated versus untreated samples were compared in triplicate using microarray analysis (Supplementary Figure 3A-F). Results were calculated using a fold change (FC) cutoff of >1.5, <-1.5, and a minimum p-value of <0.05. HCT-116 cells had 340 differentially expressed genes post-treatment (Figure 2C). Top differentially expressed genes of interest that were upregulated in HCT-116 cells included many anti-proliferative (BTG2, TP53INP1, LYZ, GADD45A, CDKN1A, ATF3, SESN1, SUSD6) and proapoptotic (DRAM1, FAS, BLOC1S2, TNFRSF10B, KLLN, PLK3, MXD1, GADD45B, TRIM31, TP53I3, TNFRSF10A, BAK1) genes (Supplementary Table 1). Of note, several of the upregulated genes are known p53 targets (BTG2, MDM2, TP53INP1, DRAM1, GADD45A, CDKN1A, PMAIP1, ATF3, FAS, SESN1, TNFRSF10D, TNFRSF10B, AEN, PLK3, TP53I3, SUSD6, GDF15) (13). Meanwhile, many of the downregulated genes included those that promote cell cycle progression (CDC25C, PRC1, ANLN, BARD1, PDK1, DHX32, CCNF, PRR11, TTK, FANCD2, AURKB, UHRF1), EMT (ENO2, MST1R) or cellular proliferation (FASN, ARHGEF39, FOXC1, CDCA3, MKI67). Another upregulated (1.78-FC) gene of interest was PPP1R1C and increased expression may increase tumor cell susceptibility to TNF-induced apoptosis (14). Interestingly, CMTM4 expression was downregulated (−1.84-FC) post-treatment and is known to protect PD-L1 from being polyubiquitinated and targeted for degradation (15). Furthermore, NEK2 was downregulated (−2.21-FC) post-treatment and NEK2 protein inhibition is known to sensitize PD-L1 blockade (16).

In HT-29 cells, we observed 2,307 differentially expressed genes post-treatment (Figure 2D). We also observed that many of the upregulated genes post-treatment were proapoptotic (AEN, TNFRSF12A, CCAR1, SFN) or anti-proliferative (SOCS7, CDKN1A, SMAD3, BCCIP, CRLF3) and many of the downregulated genes were involved with the promotion of cellular proliferation (TNIK, BRAF, EAPP, JAK1, PDS5B, CDCA3), cell cycle progression (MCIDAS, DYNC1H1, CDC45, UHRF1, CDK2, CDC25C, CCNE1, CDK1, BARD1, CCNE2), EMT (MTA3, AGGF1, E2F8, E2F7), or have antiapoptotic functions (PIM1, SGK1, BCL6, E2F7, TRIB1) (Supplementary Table 2). Interestingly, NCR3LG1 (B7-H6) was upregulated (1.78-FC) post-treatment and is known to trigger NCR3-dependent NK cell activation and cytotoxicity (17).

Finally, KM12C cells had 1,032 differentially expressed genes post-treatment (Figure 2E). We observed upregulation of proapoptotic genes (TNFRSF12A, BIK) while we observed downregulation of genes involved with the promotion of EMT (CXCL1, AGGF1, IRF2BP2, MET, NRP1, GDF15, E2F8), the promotion of cell cycle progression (CDCA2, IGFBP2, CDC25C, CCNE1, CCND2, CDK1, CCNE2, BARD1), cellular proliferation (MKI67, BRAF), and the regulation of TGFβ signaling (TGFBR2, LTBP1, TGFBR3, CD109) in KM12C cells post-treatment as compared to control (Supplementary Table 3). Of note, we noticed an upregulation (1.53-FC) of GZMA (granzyme A) expression post-treatment which is known to cleave gasdermin B to induce pyroptosis (18).

Several relevant signaling pathways had differentially expressed genes post-elraglusib in all three cell lines examined including the VEGFA-VEGFR2, TGFβ, IL-18, CCL18, EGF/EGFR, miR-targeted genes in lymphocytes, Apoptosis, and cell cycle signaling pathways (Figure 2F). The most significant commonly downregulated signaling pathway was VEGFA-VEGFR2 which had 29 downregulated genes in HCT-116 cells, 37 downregulated genes in HT-29 cells, and 48 downregulated genes in KM12C cells. A Venn Diagram was used to compare the 3,124 genes that were differentially expressed post-treatment as compared to control with elraglusib in the three colon cancer cell lines (HCT-116, HT-29, KM12C) (Figure 2G). HCT-116 cells had 241 (7.7%), HT-29 cells had 1,805 (57.8%) and KM12C cells had 549 (17.6%) differentially expressed genes post-treatment as compared to control. HCT-116 and HT-29 cells shared 46 differentially expressed genes (1.5%), HCT-116 and KM12C shared 27 (0.86%), KM12C and HT-29 shared 430 (13.8%), and all three cell lines shared 26 (0.83%). All three cell lines showed post-treatment differential expression of NF-κB regulators with increased expression of many negative regulators of NF-κB (NFKBIA, TNFAIP3, TRAIP, IL32) and decreased expression of several positive regulators of NF-κB (IRAK1BP1, FADD, IL17RA, MYD88, ERBB2IP, IL17RB, TNFSF15, NFKBIZ, NFKBIA, MAP3K1, TRAF5, TRAF6, TAB3, TNFRSF11A, MTDH, TLR3) (Supplementary Tables 1-3).

We previously found that elraglusib treatment of human CRC cell lines (HCT-116, HT-29, KM12C) with varied mutational profiles modified cytokine, chemokine, and growth factor secretion into cell culture media (19). Here, we treated tumor cells (HCT-116, HT-29) with 1 μM or 5 μM elraglusib for 48 hours and subsequently analyzed the cell culture supernatant using Luminex 200 technology (Figure 2H-I). Several cytokines, chemokines, and growth factors associated with angiogenesis and/or EMT were downregulated in both cell lines (HCT-116, HT-29) at both concentrations of elraglusib. Notably, GDF-15, GM-CSF, and VEGF all had decreased secretion post-treatment in both cell lines and at both concentrations of elraglusib. Likewise, several cytokines, chemokines, and growth factors associated with immunosuppression were also downregulated post-treatment including CCL5/RANTES, DcR3, Fas, and soluble PD-L1 (sPD-L1).

### Elraglusib enhances immune cell effector function

We next analyzed immune cell lines (TALL-104, NK-92) using western blot analysis. Interestingly, when we probed for the same proteins in the cytotoxic immune cell lysates, we observed many opposing trends to those observed in the tumor cells. In TALL-104 cells, we did not notice significant changes in NF-κB or survival protein Bcl-2 as treatment duration increased (Figure 3A). Because of the differential impact of elraglusib on tumor and immune cells that we observed via western blot, we next sought to compare the levels of another survival protein Mcl-1 in NK-92 natural killer cells and we did not observe a significant decrease in Mcl-1 protein expression through the 72-hour endpoint (Figure 3B). Surprisingly, we noted increases in survival protein Survivin and NF-κB-inducing kinase (NIK), a protein commonly associated with activation of the non-canonical NF-κB signaling pathway which led us to create a working model of NIK-mediated increased immune cell recruitment (Figure 3C). Although GSK-3 plays a role in the regulation of β-catenin, we did not focus on elraglusib-mediated effects on β-catenin because colon cancers often harbor mutations in β-catenin or adenomatous polyposis coli (APC), thus nullifying any impact GSK-3 inhibition would have on β-catenin expression. HCT116 cells are heterozygous for β-catenin, harboring one wild-type allele and one mutant allele with inactivation of one of the residues (SER45) phosphorylated by GSK3β that is frequently mutated in tumors (20). Moreover, HT-29, KM12C, and SW480 cells harbor APC mutations (21) (Supplementary Figure 2C).

**Figure 3.**
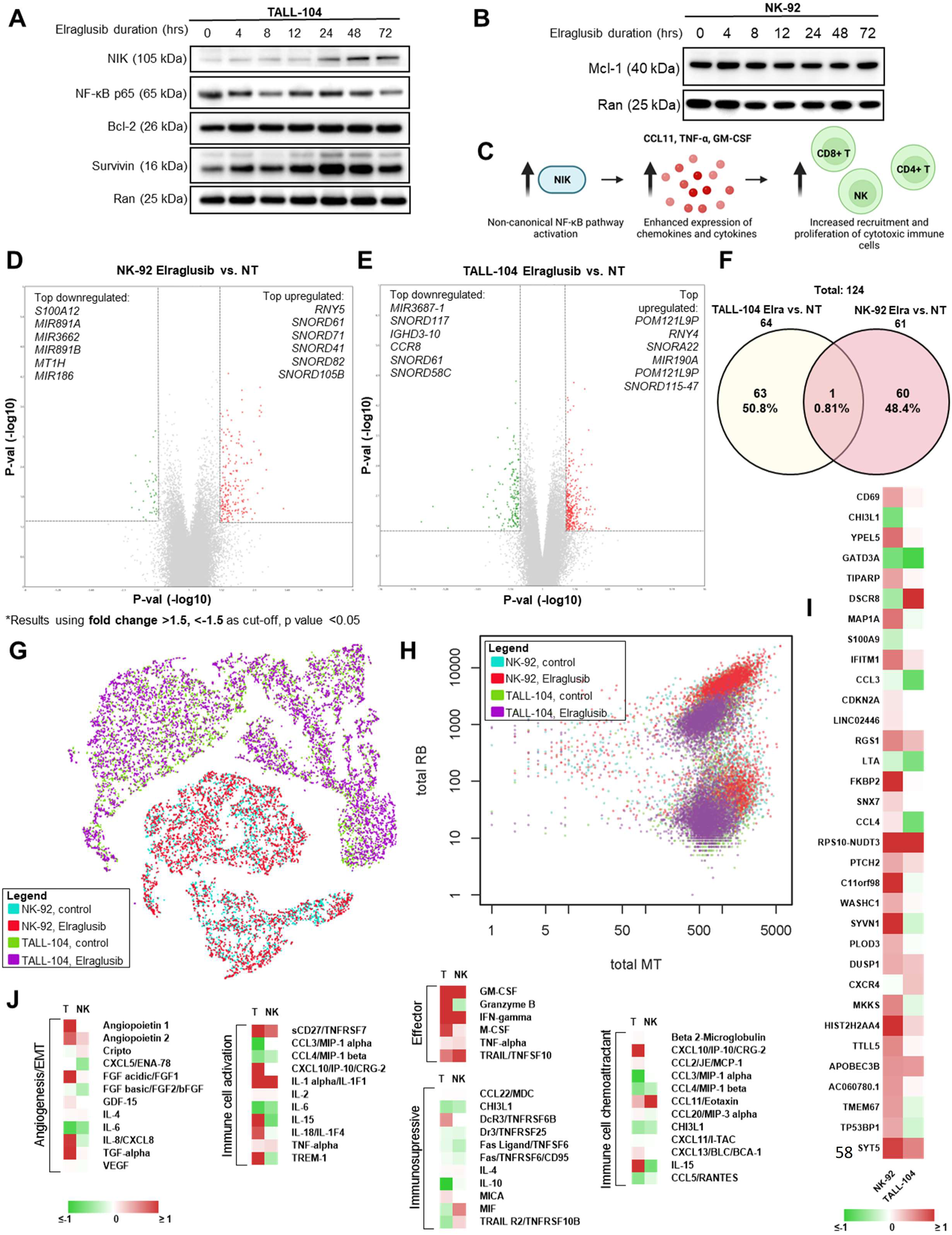
Elraglusib treatment increases effector molecule secretion and induces an energetic shift in cytotoxic immune cells. Western blot analysis of **(A)** TALL-104 and **(B)** HT-29 cytotoxic immune cells for expression of indicated proteins after increasing durations of elraglusib treatment (0-72 hours). **(C)** Proposed model for non-canonical NF-κB pathway activation: increased NIK expression indicates non-canonical NF-κB pathway activation which enhances the expression of chemokines and cytokines (CCL11, TNF-α, GM-CSF) and subsequently leads to increased recruitment and proliferation of cytotoxic immune cells (CD8+ T, CD4+ T, NK cells). Immune cells were treated with 1 μM elraglusib for 24 hours and treated versus untreated samples were compared in triplicate. Microarray analysis results were visualized using volcano plots for **(D)** NK-92 and **(E)** TALL-104 immune cell lines. **(F)** A Venn Diagram was used to compare the 124 genes that were differentially expressed post-treatment with elraglusib in the two immune cell lines (TALL-104, NK-92). **(G)** 10X single-cell sequencing analysis of immune cells treated with elraglusib. TALL-104 and NK-92 cells were treated with 1 μM elraglusib for 24 hours and aggregate data was visualized using a t-SNE plot. **(H)** Immune cells show differential expression of mitochondria-encoded genes (MT) and ribosomal genes (RB) post-treatment with elraglusib. **(I)** Heatmap comparing gene expression post-elraglusib treatment as compared to control. Red indicates a positive FC and green indicates a negative FC. **(J)** Immune cells (TALL-104, NK-92) were treated with 1 μM elraglusib for 48 hours and cell culture supernatant was analyzed with the Luminex 200. Red indicates a positive FC and green indicates a negative FC (n=3).

Next, microarray analysis was used to gain insights into gene expression changes in immune cell lines post-GSK-3 inhibition with elraglusib. Immune cell lines (TALL-104, NK-92) were treated with elraglusib at IC-50 concentrations or DMSO as vehicle control for 24 hours, and treated versus untreated samples were compared in triplicate using microarray analysis (Supplementary Figure 4A-F). Results were calculated using a FC cutoff of >1.5, <-1.5, and a minimum p value of <0.05. NK-92 cells had 61 differentially expressed genes post-treatment (Figure 3D). We observed an increase in genes that promote immune cell proliferation (TNFSF14, RAB38) and control immune cell adhesion and migration (WNK1) post-elraglusib treatment (Supplementary Table 4). We also noted decreases in proapoptotic genes (MIR186, S100A12) and genes that are involved in the activation of latent TGFβ to suppress immune cell function (ITGB8). TALL-104 cells had 64 differentially expressed genes post-treatment (Figure 3E). We observed increased expression of genes involved in the modulation of NF-κB activity (RNY4, RNY5), cytotoxic granule exocytosis (STX19, VAMP8), and anti-apoptotic gene BCL2A1 (BCL2-related protein A1). We also saw an upregulation (1.56-FC) of KIF7 (kinesin family member 7) which is required for T cell development and MHC expression as well as an increased expression (1.52-FC) of CCL3 (chemokine [C-C motif] ligand 3) which is known to recruit and enhance the proliferation of CD8+ T cells (22). In contrast, we observed decreased expression of genes involved in TGFβ signaling pathways (ACVR1B, PTPN14) and proapoptotic genes (HSPA1A, UBE3A). We also saw a decreased expression (−1.56-FC) of inhibitory immune checkpoint PTPN3 (protein tyrosine phosphatase, non-receptor type 3). In total there were 124 differentially expressed genes post-treatment and only 1, an unnamed gene (probe set ID TC22000564.hg.1, coding), was shared between both cell lines (Figure 3F).

To determine if there was any heterogeneity in response to drug treatment, we employed 10X single-cell sequencing analysis on both immune cell lines (TALL-104, NK-92) treated with low-dose 1 μM elraglusib or vehicle control (DMSO) for 24 hours. As expected, samples clustered by cell type when aggregate data was visualized using a t-SNE plot (Figure 3G). Interestingly, immune cells showed differential expression of mitochondrial-encoded genes (MT) and ribosomal genes (RB) post-treatment with elraglusib suggesting a metabolic shift in line with the extensive metabolic reprogramming undergone in immune cells post-activation (23) to support immune cell activities such as cytokine production (Figure 3H). Several genes showed the same trends post-treatment in both cell lines (Figure 3I). In both cell lines, we observed an increase in immune cell activation marker CD69 and a decrease in the immunosuppressive marker CHI3L1. Finally, we noted an increase in immune cell attractant CCL4 in the NK-92 cells and an increase in immune cell chemoattractant CXCR4 in the TALL-104 cells.

Because the previously observed non-canonical NF-κB pathway activation is known to enhance the expression of immune cell chemotactic chemokines and cytokines, we sought to determine how elraglusib treatment impacts the immune cell secretome. TALL-104 and NK-92 cells were treated with 1 µM elraglusib for 48 hours before cell culture supernatant was collected for cytokine profile analysis. TALL-104 cells treated with elraglusib showed increases in effector molecules IFN-γ, granzyme B, and TRAIL concentrations, as measured in picogram per milliliter (Figure 3J). In contrast, NK-92 cells treated with elraglusib showed increases in IFN-γ and TRAIL but showed decreases in the concentration of secreted granzyme B.

### Elraglusib significantly prolongs survival in combination with anti-PD-L1 therapy in a syngeneic MSS CRC murine model

Because elraglusib activated immune cells and increased tumor cell PD-L1 expression, we sought to evaluate the potential for elraglusib to increase the efficacy of ICB and utilized a syngeneic murine colon carcinoma BALB/c murine model using a MSS cell line CT-26 (Figure 4A). In this MSS CRC model, we observed a significantly improved survival curve in the elraglusib and anti-PD-L1 combination therapy group (Figure 4B). We also observed statistically significant improved survival in the elraglusib, anti-PD-1, and anti-PD-L1 alone groups as compared to the control (Supplementary Figure 5A-E). However, we saw the most sustained response in the elraglusib and anti-PD-L1 combination therapy group (Supplementary Figure 5F). Murine body weights did not differ significantly regardless of the treatment group (Supplementary Figure 5G-L). Also, the mice did not show evidence of significant treatment-related toxicity on complete blood count or serum chemistry analysis (Supplementary Table 5). Both the elraglusib individual treatment and dual treatment groups maintained normal renal function as evidenced by normal blood urea nitrogen (BUN) and creatinine and were free of significant electrolyte perturbations. Liver function tests did not reveal any evidence of liver toxicity and the dual-treatment mice did not have any elevations of AST, ALT, or bilirubin. As can be expected in mice with significant tumor burdens, mice across treatment groups had decreased albumin levels and evidence of mild marrow hypoplasia resulting in mild anemia, and lower white blood cell and platelet counts. This effect was independent of the treatment group and likely related to tumor burden at the time of sacrifice.

**Figure 4.**
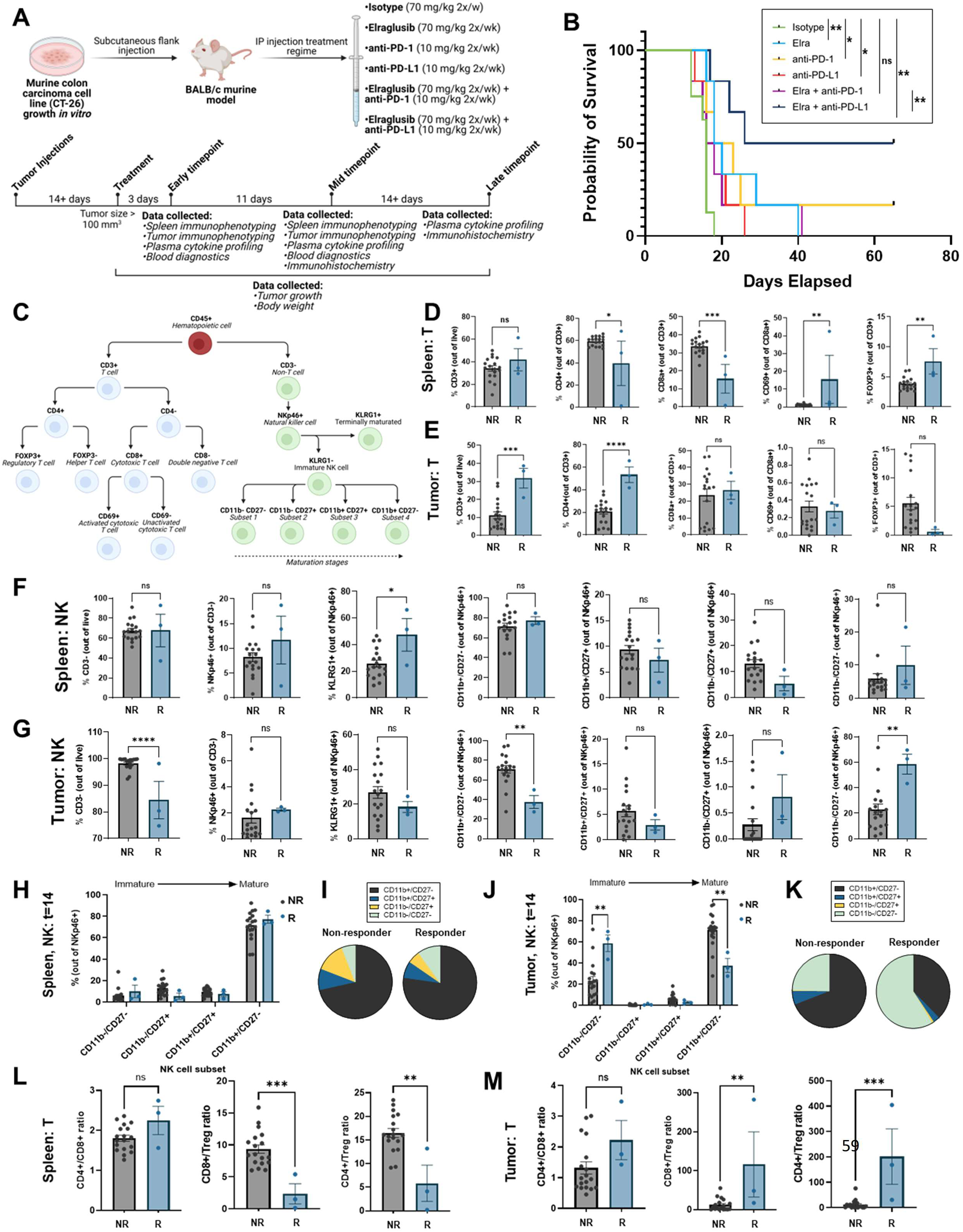
Elraglusib enhances immune cell tumor-infiltration to prolong survival in combination with anti-PD-L1 therapy in a syngeneic murine model of MSS colon carcinoma. **(A)** Experimental model overview of the syngeneic murine colon carcinoma BALB/c murine model using MSS cell line CT-26. **(B)** Kaplan–Meier estimator curves for all treatment groups as indicated. Statistical significance was determined using a Log-rank (Mantel-Cox) test. **(C)** Overview of cell lineage markers used for flow cytometric immunophenotyping analysis. 14-days post-treatment initiation immune cell subpopulations were analyzed in the spleen and tumor. Splenic T cells, **(E)** Tumor-infiltrating T cells, **(F)** Splenic NK cells, and **(G)** Tumor-infiltrating NK cells were compared between responders (R, n=3) and non-responders (NR, n=18). NK cell subsets based on the expression of CD11b and CD27 were compared in the spleen and visualized via **(H)** bar graph and **(I)** pie chart. NK cell subsets based on the expression of CD11b and CD27 were also compared in the tumor and visualized via **(J)** bar graph and **(K)** pie chart. T cell ratios were compared in the **(L)** Spleen and **(M)** Tumor. Statistical significance was determined using two-tailed unpaired T tests. P-value legend: * p < 0.05, ** p < 0.01, *** p < 0.001, **** p < 0.0001.

### Murine responders have more T cell tumor-infiltration and higher tumoral CD8+/Treg and CD4+/Treg ratios

To begin to evaluate our hypothesis that elraglusib increases immune cell activation and recruitment, we utilized multi-color flow cytometry to characterize the natural killer (NK) and T cell populations 14-days post-treatment initiation, and immune cell subpopulations were analyzed in both the spleen and the tumor (Figure 4C). 14 days post-treatment initiation, mice were grouped as responders **(R)** or non-responders (NR) based on a tumor volume less than or greater than 100 mm^3^, respectively. Compared to non-responders, regardless of treatment group, responders 14-days post-treatment had statistically significantly lower levels of splenic CD4+ and CD8+ T cells and had increased percentages of CD69+ activated T cells and Foxp3+ regulatory T cells (Tregs) (Figure 4D). Meanwhile, responders had increased percentages of tumor-infiltrating CD3+ and CD4+ T cells (Figure 4E). We also observed that responders had increased percentages of splenic KLRG1+ mature NK cells and tumor-infiltrating CD11b-/CD27-immature NK cells, and decreased percentages of tumor-infiltrating CD11b+/CD27-activated NK cells 14-days post-treatment initiation (Figure 4F-G). We did not observe striking differences between non-responders and responders in the splenic immature natural killer cell subsets (CD11b-/CD27-, CD11b-/CD27+, CD11b+/CD27+, CD11b+CD27-) (Figure 4H-I). In contrast, we did observe significant differences between non-responders and responders in the tumor-infiltrating immature natural killer cell subsets (Figure 4J-K). We observed that responders had a greater proportion of immature (CD11b-/CD27-) NK cells and a lower proportion of mature (CD11b+CD27-) NK cells 14 days post-treatment initiation. When comparing the T cell ratios, compared to non-responders, responders had a lower splenic CD8+/Treg and CD4+/Treg ratio (Figure 4L). The CD8+/Treg ratio is commonly used as an index of TIL’s effector function (24). Additionally, responders had a higher intra-tumoral CD8+/Treg and CD4+/Treg ratio (Figure 4M). Overall, the observed changes in immune cell subsets in responders are consistent with increased infiltration of cytotoxic immune cells into the tumor.

### Murine responders show an immunostimulatory tumor microenvironment by IHC

To further interrogate the tumor microenvironment (TME), we utilized immunohistochemistry (IHC) analysis on tumor sections from the 14-day post-treatment initiation timepoint or from the end-of-study (EOS) timepoint. We compared non-responders (NR) and responders **(R)** and stained for T cell marker CD3 and observed that responders had significantly more CD3+ T cells as compared to non-responders at both timepoints analyzed. (Figure 5A-B). To determine if there were any differences in immune cell activation, we stained for Granzyme B and again observed that responders had significantly more Granzyme B+ staining at both timepoints analyzed as compared to non-responders (Figure 5C-D). We stained for Ki67 as a marker of tumor cell proliferation and observed that responders had less tumor cell proliferation as compared to responders at both of the timepoints analyzed (Figure 5E-F). Because we found that elraglusib upregulated tumor cell PD-L1 expression and because we observed such a striking difference in survival when elraglusib was combined with anti-PD-L1 therapy as compared to anti-PD-1 therapy, we next looked at PD-L1 staining in the tumor sections (Figure 5G-H). We observed that responders had more PD-L1+ tumor cells as compared to non-responders at both timepoints analyzed. To examine tumor cell apoptosis, we then stained for cleaved-caspase 3 (CC3) and noted that there was no difference in CC3 expression at the mid-timepoint (14 days post-treatment initiation), however, responders did have significantly more CC3 expression than non-responders at the EOS timepoint (Figure 5I-J). We also analyzed the expression of CD4, CD8, Foxp3+, NKp46, TRAIL, PD-1, VEGF, and TGFβ2 to gain additional insights into the tumor immune microenvironment at both the 14 days post-treatment initiation timepoint and the EOS timepoint, respectively (Supplementary Figure 6). To examine helper T cell presence, we stained for CD4 and observed that responders had more CD4+ T cells than non-responders at both of the timepoints examined (Supplementary Figure 6A). Interestingly, we saw the same trends when we examined CD8 expression where responders had more CD8+ T cells as compared to non-responders which differed from the flow cytometry results but could be explained by the large variability in CD8a+ T cells we observed by flow cytometry in the non-responder group (Supplementary Figure 6B). We did not observe statistically significant differences in Foxp3+ Treg expression between responders and non-responders at either timepoint (Supplementary Figure 6C). When we examined NK cell tumor-infiltration by IHC we noted more NKp46+ NK cells in responders at the 14-day post-treatment initiation timepoint, but this difference was not significant at the EOS timepoint (Supplementary Figure 6D). We chose to examine another cytotoxic mediator TRAIL, and observed no difference between responders and non-responders at the mid-timepoint but, interestingly, observed less TRAIL expression in the responders as compared to the non-responders at the EOS timepoint (Supplementary Figure 6E). We also examined PD-1 expression and did not note any significant differences between responders and non-responders at either of the timepoints examined (Supplementary Figure 6F). Again, we noted a similar lack of significance when we examined immunosuppressive and angiogenic VEGF expression (Supplementary Figure 6G). Finally, we examined immunosuppressive TGFβ2 expression and noted no differences between responders and non-responders at the mid-timepoint but noted that responders had significantly lower expression at the EOS timepoint (Supplementary Figure 6H). Signal was quantified by converting randomly sampled 20X images into 16-bit images and then utilizing Fiji to employ MaxEntropy thresholding (Supplementary Figure 7).

**Figure 5.**
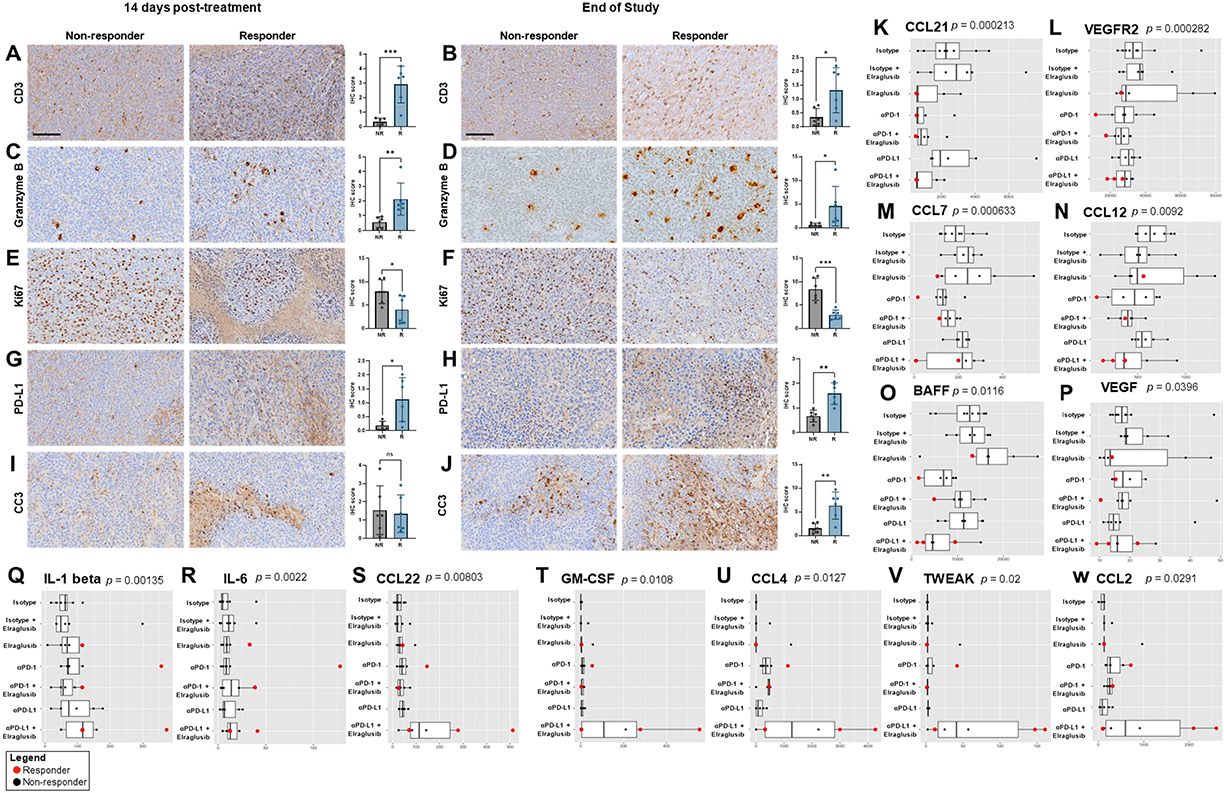
Responders have a more immunostimulatory tumor microenvironment as compared to non-responders. IHC analysis of tumors 14 days post-treatment initiation or tumors from long-term mice. Non-responders (NR) and responders **(R)** were compared. 20X images, scale bar represents 100 μm. (A-B) CD3, (C-D) Granzyme B, (E-F) Ki67, (G-H) PD-L1, and (I-J) cleaved-caspase 3 (CC3) were compared at the 14 days post-treatment initiation timepoint, and the long-term timepoint, respectively. Statistical significance was determined using two-tailed unpaired T tests (n=6). Serum from long-term mice sacrificed was analyzed via cytokine profiling for **(K)** CCL21, **(L)** VEGFR2, **(M)** CCL7, **(N)** CCL12, **(O)** BAFF, **(P)** VEGF, **(Q)** IL-1 β, **(R)** IL-6, **(S)** CCL22, **(T)** GM-CSF, **(U)** CCL4, **(V)** TWEAK, and **(W)** CCL2. Responders (red) and non-responders (black) were compared. A Kruskal-Wallis test was used to calculate statistical significance followed by a Benjamini-Hochberg correction for multiple comparisons. p values are shown for analytes that were significantly different between responders and non-responders and are ordered by significance. P-value legend: * p < 0.05, ** p < 0.01, *** p < 0.001, **** p < 0.0001.

### Murine responders have lower tumorigenic and higher immunomodulatory cytokine concentrations

We next analyzed murine serum samples from EOS mice for cytokine profiles and noted interesting trends between responders and non-responders. Responders were more likely to have lower serum concentrations of CCL21 (p=0.000213), VEGFR2 (p=0.000282), CCL7 (p=0.000633), CCL12 (p=0.0092), BAFF (p=0.0116), and VEGF (p=0.0396) compared to non-responders (Figure 5K-P). In contrast, responders had higher serum concentrations of IL-1 β (p=0.00135), IL-6 (p=0.0022), CCL22 (p=0.00803), GM-CSF (p=0.0108), CCL4 (p=0.0127), TWEAK (p=0.02), and CCL2 (p=0.0291) compared to non-responders (Figure 5Q-W).

Analytes that were statistically significant between responders and non-responders at both timepoints (14 days post-treatment initiation, EOS) included CCL7/MCP-3/MARC (p=2.19E-05), CCL12/MCP-5 (p=0.000606), TWEAK/TNFSF12 (p=0.00112), BAFF/TNFSF13B (p=0.00469), IL-1 β/IL-1F2 (p=0.00507), CCL21/6Ckine (p=0.00539), VEGF (p=0.00646), IFN-γ (p=0.00817), CCL4/MIP-1 β (p=0.0133), IL-6 (p=0.229), and GM-CSF (p=0.0257). When comparing responders and non-responders, a Kruskal-Wallis test was used to calculate statistical significance followed by a Benjamini-Hochberg correction for multiple comparisons. The entire panel of cytokines, chemokines, and growth factors analyzed by multiplex immunoassay in murine serum from the EOS timepoint included BAFF, MCP-1, MIP-1 α, MIP-1 β, RANTES, MCP-3, Eotaxin, MCP-5, VEGFR2, MIP-3 α, CCL21, MDC, IP-10, CXCL12, GM-CSF, Granzyme B, IFN-γ, IL-1 α, IL-18, IL-2, IL-3, IL-4, IL-6, IL-7, IL-10, IL-12 p70, IL-13, IL-16, VEGF, M-CSF, Prolactin, and TWEAK (Supplementary Figure 8A-S).

### Patient plasma concentrations of cytokines from a Phase 1 clinical trial investigating elraglusib correlate with progression-free survival, overall survival, and in vivo response to therapy results

To determine the clinical relevance of the biomarkers of response identified in our murine model, we next employed Luminex 200 technology to analyze plasma samples from patients with refractory solid tumors of multiple tissue origins enrolled in a Phase 1 clinical trial investigating elraglusib (<u>NCT03678883</u>). We found that baseline concentrations of several analytes (IL-12, CXCL11, Fas Ligand, IL-8, VEGF, IL-1 β, M-CSF, IL-2) correlated with progression-free survival (PFS) (Figure 6A). Heatmaps were used to visualize linear regression, R squared, and p values. Likewise, concentrations of several analytes (IL-12, IL-1 β, IL-21, IL-8, IFN-α, IFN-γ, M-CSF, CCL4, Fas Ligand, IL-2, IL-10, CCL11, IL-15, IL-4, Granzyme B, CXCL11) 24-hours post-dose also correlated with PFS (Figure 6B). We next analyzed overall survival (OS) data and noted that baseline concentrations of several analytes (IL-8, CXCL11, CCL11, IFN-α, TNF-α, Fas Ligand, TRAIL R2, IL-1 β) correlated with OS (Figure 6C). Next, 24-hour post-dose concentrations of several analytes (IFN-α, Fas Ligand, TRAIL R2, CCL11) also correlated with OS (Figure 6D). Patients included in this analysis represented multiple tumor types including appendix (n=3, 15.8%), adult T-cell leukemia/lymphoma (ATLL) (n=1, 5.3%), cholangiocarcinoma (n=1, 5.3%), colorectal (n=7, 36.8%), desmoid (n=1, 5.3%), hepatocellular carcinoma (HCC) (n=1, 5.3%), leiomyosarcoma (n=1, 5.3%), non-small cell lung cancer (NSCLC) (n=2, 10.5%), and pancreas (n=2, 10.5%) cancer (Figure 6E). The median PFS was 75.9 days, and the median OS was 101 days (Supplementary Table 6).

**Figure 6.**
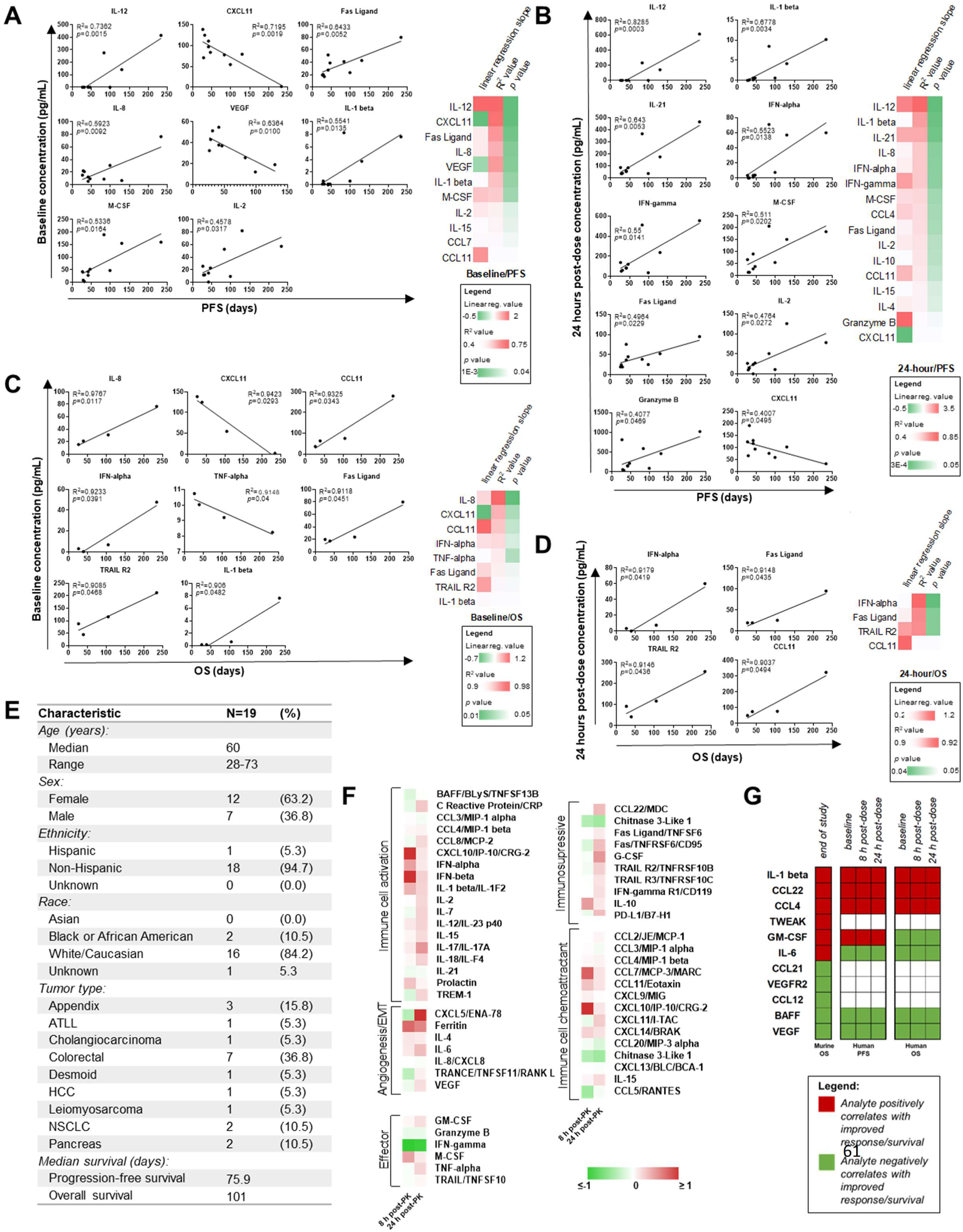
Patient plasma concentrations of cytokines correlate with progression-free survival (PFS), overall survival (OS), and in vivo response to therapy results. Plasma samples from human patients with refractory solid tumors of multiple tissue origins enrolled in a Phase 1 clinical trial investigating a novel GSK-3 inhibitor elraglusib (NCT03678883) were analyzed using a Luminex 200 (n=19). **(A)** Baseline analyte concentrations in pg/mL were plotted against PFS. **(B)** 24-hour post-dose analyte concentrations in pg/mL were plotted against PFS. **(C)** Baseline analyte concentrations in pg/mL were plotted against OS. **(D)** 24-hour post-dose analyte concentrations in pg/mL were plotted against OS. Simple linear regressions were used to calculate significance. R squared and p values were reported. p values less than 0.05 were reported as statistically significant. Graphs are ordered from most to least significant starting at the upper left. Heatmaps show linear regression slope values, R squared values, and p values ordered by significance starting from the top. **(E)** Table summarizing demographics. **(F)** Cytokines grouped by function. FC is shown where green indicates a negative (<0) FC compared to the baseline (pre-dose) value and red indicates a positive (>0) FC. **(G)** Table comparing murine and human circulating biomarker trends. Red boxes indicate that an analyte concentration positively correlated with response to therapy/PFS/OS while green boxes indicate that an analyte concentration negatively correlated with response to therapy/PFS/OS.

Many of the analytes were upregulated at 8-and 24-hours post-dose as compared to baseline (Figure 6F). When the cytokines, chemokines, and growth factors were grouped by timepoint and raw values were visualized with a heatmap we noticed several interesting trends (Supplementary Figure 9A). When grouped by primary tumor location (appendix, adult T cell leukemia/lymphoma [ATLL], cholangiocarcinoma, colorectal, desmoid, hepatocellular carcinoma [HCC], leiomyosarcoma, non-small cell lung cancer [NSCLC], pancreatic), we noted that the patient with a desmoid tumor had elevated expression of many of the analytes included in the panel. When cytokines were grouped by elraglusib dose (1, 2, 3.3, 5, 7, 9.3, 12.37) in milligrams per kilogram, we noted that patients receiving a 7 mg/kg dose had increased expression of many of the analytes included in the panel at both the 8-and 24-hour post-dose timepoints (Supplementary Figure 9B). Finally, when cytokines were grouped by cytokine, chemokine, or growth factor family we noted that TNF family molecules (BAFF, Fas Ligand, Fas, TNF-α, TRAIL R2, TRAIL R3, TRAIL, TRANCE) has a decreased expression at the 8-hour post-dose timepoint as compared to baseline and had increased over baseline levels by the 24-hour timepoint (Supplementary Figure 9C).

To compare both murine and human circulating biomarker trends, we created a table to visualize major trends (Figure 6G). EOS analyte concentrations that positively correlated with OS in the mouse model included IL-1 β, CCL22, CCL4, TWEAK, GM-CSF, and IL-6. Those that negatively correlated with OS in the mouse model included CCL21, VEGFR2, CCL12, BAFF, and VEGF. Interestingly, we observed that many of these trends held true when analyzing the human data. IL-1 β, CCL22, and CCL4 all were positively correlated with PFS and OS in the human cohort, likewise, BAFF and VEGF were negatively correlated with OS and PFS. GM-CSF and IL-6 had opposing correlations in the human cohort as compared to the murine cohort.

### PanCK+ expression of immunosuppressive CD39 negatively correlated with time-on-treatment (Tx time) while CD45+ expression of monocyte/macrophage marker CD163 positively correlated with Tx time

To gain insights into the human TME post-elraglusib, we utilized GeoMx Digital Spatial Profiling (DSP) technology to profile the expression of 59 proteins in tumor biopsies (n=12) from patients treated with elraglusib (n=7). 42% (n=5) of the tumor biopsies analyzed were collected near or before treatment start (pre-treatment) and 58% (n=7) of the biopsies analyzed were collected from post-treatment (average time-on-treatment [Tx time] at post-treatment biopsy: 270 days) (Figure 7A). Primary tumor types included CRC (n=4, 33%) and pancreatic cancer (n=8, 67%), while metastatic biopsy tissue sites included lung (n=2, 17%), liver (n=7, 58%), rectum (n=2, 17%), and pleura (n=1, 8%). We analyzed five paired tumor biopsies (n=10 slides total, 83%), and 2 unpaired biopsies (n=2 slides total, 17%). Half (n=6, 50%) of the tumor sections were needle biopsies (Supplementary Figure 10) and half (n=6, 50%) were tissue biopsies (Supplementary Figure 11). All patients included in this analysis were considered responders based on the definition used in the Phase 1 trial that treatment response is equal to disease control greater than 16 weeks. Our region of interest (ROI) selection strategy focused on mixed tumor and immune cell segments within FFPE tissue. ROIs were segmented based on panCK+ and CD45+ morphology stains to compare tumor versus immune cells protein expression (Figure 7B, Supplementary Figure 12). We utilized a PCA plot to visualize dimensionality reduction and as expected, samples tended to cluster by tissue type (liver, lung, pleura, rectum) and further separated by segment (CD45, panCK) on PC2 (Supplementary Figure 13A-B). We utilized a Sankey diagram to visualize the study design where the width of a cord in the figure represents how many segments are in common between the two annotations they connect (Figure 7C). Also as expected, samples tended to cluster together based on patient ID, primary tumor location, biopsy timepoint, metastatic biopsy tissue site, immune cell location, or segment (CD45, panCK) type when visualized on an aggregate heatmap (Figure 7D). As we were interested in the ability to predict a patient’s time-on-treatment (Tx time), we sought to correlate pre-treatment protein expression levels among the responders with Tx time data and found that PanCK+ segment expression of immunosuppressive CD39 negatively correlated with Tx time (Figure 7E) while CD45+ segment monocyte/macrophage marker CD163 expression positively correlated with Tx time (Figure 7F) (25).

**Figure 7.**
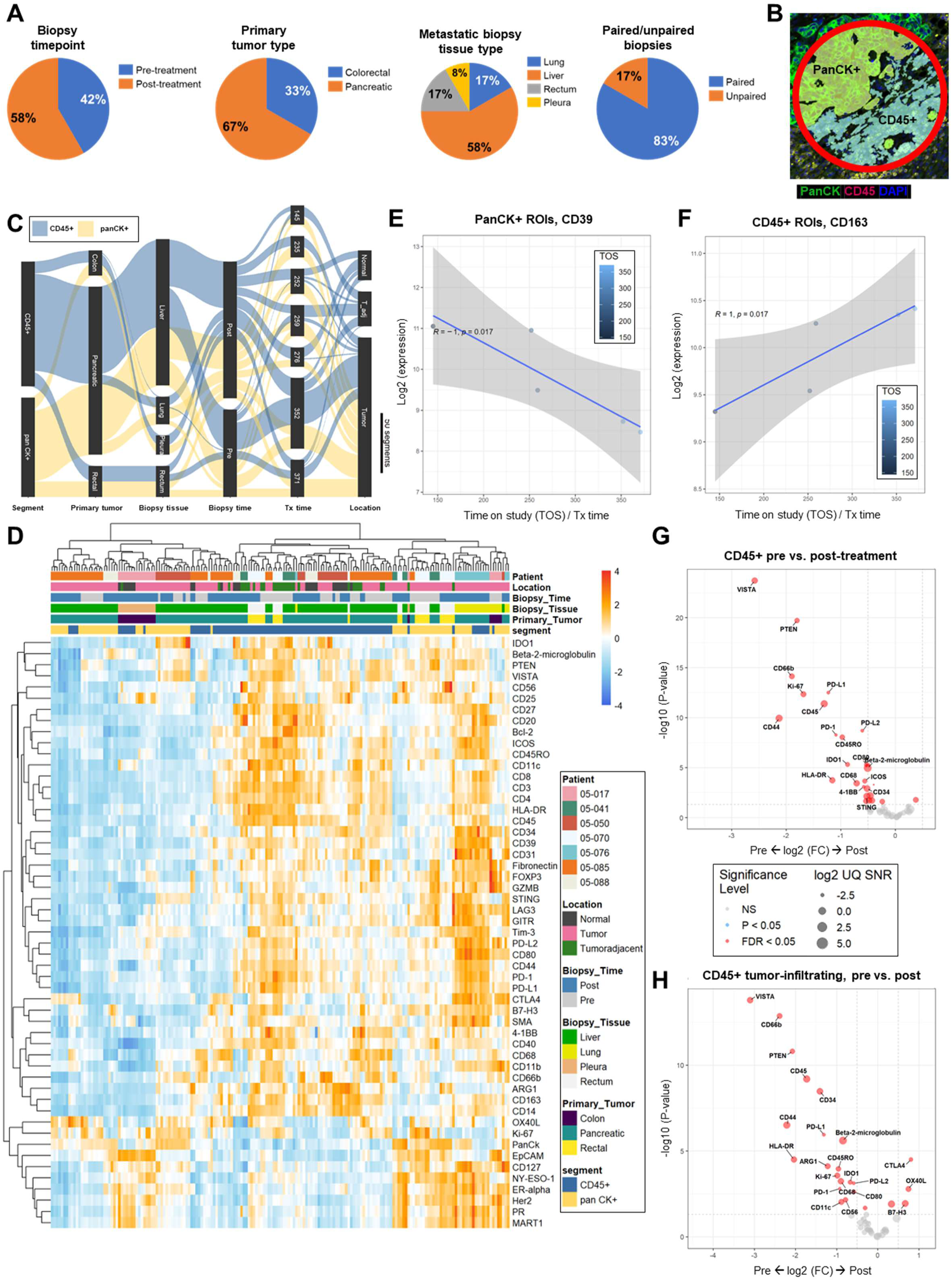
Spatial profiling of patient tumor biopsies reveals a more immunostimulatory tumor microenvironment post-treatment with elraglusib. Patient samples were analyzed using NanoString GeoMx Digital Spatial Profiling (DSP) technology. **(A)** Pie charts showing biopsy timepoint, primary tumor type, metastatic biopsy tissue type, and paired/unpaired biopsy sample information breakdowns. **(B)** A representative region of interest (ROI) showing PanCK+ and CD45+ masking. Green indicates CK, red indicates CD45, and blue indicates DAPI staining. **(C)** A Sankey diagram was used to visualize the study design where the width of a cord in the figure represents how many segments are in common between the two annotations they connect. The scale bar represents 50 segments. Blue cords represent CD45+ segments and yellow cords represent panCK+ segments. **(D)** Heatmap of all areas of interest (AOIs). Patient IDs, immune cell locations, biopsy timepoint, biopsy tissue, primary tumor location, and segment identity information are color coded as indicated in the legend. **(E)** PanCK+ ROI CD39 expression plotted against time-on-study (TOS). Points are color-coded by time on study (TOS) / time on treatment with darker blue points indicating a shorter TOS or time on treatment. **(F)** CD45+ ROI CD163 expression plotted against TOS. **(G)** Volcano plot showing a comparison of CD45+ region protein expression in post-treatment biopsies and pre-treatment biopsies regardless of timepoint. Grey points are non-significant (NS), blue points have p values < 0.05, and red points have false discovery rate (FDR) values less than 0.05. The size of the point represents the log2 UQ Signal-to-noise ratio (SNR). **(H)** Volcano plot showing a comparison of tumor-infiltrating CD45+ immune cell segment protein expression in pre-versus post-treatment biopsies. Grey points are non-significant (NS), blue points have p values < 0.05, and red points have false discovery rate (FDR) values less than 0.05. The size of the point represents the log2 UQ Signal-to-noise ratio (SNR).

### Tumor-infiltrating immune cells have reduced inhibitory checkpoint expression and increased expression of T cell activation markers post-elraglusib

When comparing all samples, CD45+ regions of post-treatment biopsies had increased protein expression of T cell activation marker OX40L (p = 0.016) and decreased protein expression of checkpoint molecules VISTA (p = 2.0E-24), PD-L1 (p = 3.2E-13), PD-L2 (p = 2.0E-9), LAG3 (p = 5.1E-4), and PD-1 (p = 5.6E-9). CD45+ regions of post-treatment biopsies also had decreased protein expression of myeloid/neutrophil marker CD66b (p = 7.5E-15), myeloid markers IDO1 (p = 4.8E-6), CD80 (p = 5.4E-6), and CD11b (p = 6.7E-3), TAM/M2 macrophage marker CD68 (p = 3.8E-4), myeloid/T cell activation marker OX40L (p = 0.016), myeloid marker CD40 (p = 0.020), and DC/myeloid marker CD11c (p = 0.022) as compared to pre-treatment samples (Figure 7G). Because we were interested in the differential expression of proteins based on immune cell location in relation to the tumor, we annotated CD45+ ROI locations as tumor-infiltrating, tumor-adjacent, or normal tissue (Supplementary Figure 14A-C). When next compared tumor-infiltrating CD45+ immune cell segments in pre-versus post-treatment biopsies and found that post-treatment tumor-infiltrating CD45+ immune cell segments had lower protein expression of immune checkpoints VISTA (p = 1.6E-14), PD-L1 (p = 1.1E-6), PD-L2 (p = 7.6E-4), and PD-1 (p = 1.6E-3) and higher protein expression of T cell activation markers CTLA4 (p = 3.1E-5) and OX40L (p = 1.6E-3) (Figure 7H). We also noted that post-treatment tumor-infiltrating CD45+ immune cell segments had lower protein expression of myeloid marker CD66b (p = 1.4E-13), antigen PTEN (p = 1.5E-11), hematopoietic marker CD34 (p = 3.3E-9), T cell activation marker CD44 (p = 3.1E-7), antigen presentation B2M (p = 2.5E-6), immune cell activation marker HLA-DR (p = 3.1E-5), TAM/M2 macrophage marker ARG1 (p = 7.6E-5), memory T cell marker CD45RO (p = 1.1E-4), proliferation marker Ki-67 (p = 2.7E-4), TAM/M2 macrophage marker CD68 (p = 5.8E-4), myeloid marker IDO1 (p = 6.6E-4), myeloid marker CD80 (p = 2.4E-3), NK cell marker CD56 (p = 6.9E-3), DC/myeloid marker CD11c (p = 9.5E-3), and T cell activation marker GITR (p = 0.021) and had higher protein expression of immune checkpoint molecule B7-H3 (p = 0.012) and Treg marker CD127 (p = 0.012) as compared to pre-treatment tumor-infiltrating CD45+ immune cell segments.

### Patients with a long time-on-treatment have decreased B cell and myeloid marker expression in immune cell regions and have decreased immune checkpoint expression in tumor cell regions

Next, we sought to compare pre-treatment biopsy protein expression in CD45+ segments between patients who were on treatment for a longer duration of time called “Long Tx patients” and patients who were on the study for a shorter duration of time called “Short Tx patients” and observed that Long Tx patients had lower protein expression of B cell marker CD20 (p = 0.012) and myeloid activation marker CD80 (p = 0.047) (Supplementary Figure 13C). Long Tx was defined as a Tx time greater than 275 days (∼39 weeks). Then we compared protein expression in CD45+ segments in post-treatment biopsies between Long Tx patients and Short Tx patients. We observed that Long Tx patients had lower protein expression of antigen NY-ESO-1 (p = 0.021) and progesterone receptor (PR) (p = 0.022) (Supplementary Figure 13D). We then compared pre-treatment biopsy protein expression in PanCK+ segments between Long Tx patients and Short Tx patients and observed that Long Tx patients had lower protein expression of cytotoxic T cell marker CD8 (p = 3.5E-3), antigen Her2 (p = 0.033), Treg marker Foxp3 (p = 0.033), T cell marker CD3 (p = 0.035), and B cell marker CD20 (p = 0.046). Long Tx patients also had lower immune checkpoint protein expression of LAG3 (p = 0.023), PD-L2 (p = 0.028), and PD-1 (p = 0.046) (Supplementary Figure 13E). We carried out the same analysis with a focus on panCK+ segments in post-treatment biopsies. Long Tx patients had lower protein expression of mature B cell/DC marker CD35 (p = 8.5E-3), antigen NY-ESO-1 (p = 8.7E-3), antigen Her2 (p = 0.022), antigen MART1 (p = 0.029), cytotoxic T cell marker CD8 (p = 0.030), Treg marker Foxp3 (p = 0.030), antigen PTEN (p = 0.032), DC/myeloid marker CD11c (p = 0.034), memory T cell marker CD45RO (p = 0.036), checkpoint PD-L1 (p = 0.047), and PR (p = 0.049) as compared to Short Tx patients (Supplementary Figure 13F). Several additional comparisons were made between pre-and post-treatment biopsies, immune cell location in proximity to the tumor, and paired and unpaired biopsies (Supplementary Figure 15A-N).

## Discussion

ICB is a promising treatment strategy for many cancer patients, including MSI-H CRC patients. However, the response rate to ICB in MSS CRC patients is very limited, especially as the tumor stage advances, thus there is a clear need for improved treatment strategies for this patient population. Evaluating the combination of ICB with small molecules in oncology represents one of the ways we might improve the efficacy of ICB in MSS CRC patients. Here, we focus on small-molecule inhibitor of GSK-3 elraglusib and characterize several immunomodulatory mechanisms that provide a clinical rationale for the combination of GSK-3 inhibitors such as elraglusib in combination with ICB.

We demonstrate that small-molecule inhibition of GSK-3 using elraglusib leads to increased natural killer and T cell-mediated CRC cell killing in a co-culture model. Moreover, elraglusib acts on tumor cells to sensitize them to immune cell-mediated killing. This tumor cell sensitization could be resultant of drug-induced modifications in the tumor cell secretome such as decreased VEGF expression, decreased soluble PD-L1, and increased CXCL14, as we previously described (19). VEGF has been shown to inhibit T cell activation (26) while CXCL14 is a known NK cell chemoattractant (27). It has been shown that the soluble or shed version of PD-L1 can retain the ability to bind PD-1 and function as a decoy receptor to negatively regulate T cell function, despite being a truncated version lacking the membrane domain of the protein (28). Therefore, the increase in efficacy in combination with ICB that we observed in the co-culture model could be due to a concomitant downregulation of sPD-L1 and an upregulation of cell surface-expressed PD-L1. This in vitro work was presented, in part, at the 2021 American Association of Cancer Research (AACR) Annual Conference.

Elraglusib-mediated immunostimulation may also function, in part, by inducing pyroptosis in cancer cells. Pyroptosis is a lytic and pro-inflammatory type of programmed cell death that results in cell swelling and membrane perforation. Although the role of pyroptosis in cancer is controversial, it has been suggested that pyroptosis may contribute to anti-tumor immunity (29). Since we observed gasdermin B expression post-IFN-γ treatment in CRC cells and because we found that elraglusib treatment upregulated immune cell IFN-γ secretion, we hypothesize that the IFN-γ released from CD8+ T cells and NK cells is responsible for triggering pyroptosis which may, in part, contribute to elraglusib-mediated immunostimulation.

Another mechanism behind elraglusib-mediated immunomodulation is the suppression of inflammatory NF-κB signaling and survival pathways in the tumor cells. We demonstrated that elraglusib treatment of CRC cells decreased Survivin, NF-κB p65, Bcl-2, and Mcl-1expression while increasing PD-L1 expression. This is in accordance with previous studies that have shown that GSK-3 is a positive regulator of NF-κB (30). Microarray data showed increased expression of antiproliferative, proapoptotic, and NF-κB regulator genes and decreased expression of genes involved in cell cycle progression, antiapoptotic, and EMT genes in CRC cell lines. Multiplex immunoassay data showed decreased tumor cell secretion of proteins involved in angiogenesis, EMT, and immunosuppression.

Meanwhile, we observed the opposite effect on NF-κB signaling in immune cells, where we observed that drug treatment increased NF-κB-inducing kinase (NIK) expression. NIK is the upstream kinase that regulates activation of the non-canonical NF-κB signaling pathway, and may implicate a role for non-canonical NF-κB signaling in immune cells post-treatment with elraglusib, which future studies could further evaluate. It is known that increased expression of NIK leads to enhanced expression of chemokines and cytokines such as CCL3, TNF-α, and MCP-1, which thus leads to increased recruitment and proliferation of cytotoxic immune cells (31). Moreover, elraglusib treatment of immune cells increased effector molecule secretion in both T and NK cells as well as led to increased expression of genes involved in cytotoxic granule exocytosis, cellular proliferation, and modulators of NF-κB activity. Moreover, elraglusib treatment resulted in decreased gene expression of proapoptotic molecules and regulators of TGFβ signaling which may also contribute to the tumor suppressive and anti-angiogenic effects of elraglusib that have been previously described (32).

In a syngeneic murine colon carcinoma BALB/c murine model using MSS cell line CT-26, we observed significantly improved survival of mice treated with elraglusib and anti-PD-L1 therapy. We also demonstrated increased survival of mice treated with elraglusib alone as compared to the control group. We also observed statistically significant improved survival in the anti-PD-1 and anti-PD-L1 alone groups as compared to the control. Responders had lower percentages of splenic CD4+ T cells and splenic CD8+ T cells and had increased percentages of CD69+ activated T cells and Foxp3+ Tregs. The increased splenic percentages of both activated and end-stage T cells in the responder groups could be indicative of an anti-tumor immune response that was mounted earlier in the treatment course. Future studies could analyze the changes in these immune cell populations during the course of therapy in greater detail, especially considering we could have missed important changes in immune cell subtypes due to limited timepoints. Compared to non-responders, responders also had more CD3+ and CD4+ tumor-infiltrating lymphocytes. Further studies could evaluate the contribution of CD4+ versus CD8+ tumor-infiltrating T cells to the observed response to elraglusib and anti-PD-L1 therapy, especially considering the recent interest in the contribution of CD4+ helper T cells to anti-tumor immunity (33). We did not observe many significant differences in splenic NK cell subpopulations in either the tumor or the spleen, although perhaps the timepoint we chose to analyze was not representative of NK cell subpopulation changes that may have occurred earlier or later in the course of treatment. One limitation of this model is that it is a heterotopic flank tumor model as opposed to an orthotopic colon tumor model which may be more representative of the CRC TME. Follow-up experiments could examine the contribution of CD4+ T cells, CD8+ T cells, and NK cells to response to therapy in the murine MSS CRC model by blocking the function of each cell population in individual experiments.

We observed that murine responders had lower serum concentrations of BAFF, CCL7, CCL12, VEGF, VEGFR2, and CCL21. BAFF is a cytokine that belongs to the TNF ligand superfamily, that may promote tumorigenesis indirectly by induction of inflammation in the TME and directly by induction of EMT (34). Meanwhile, CCL7 has been shown to enhance both cancer progression and metastasis via EMT, including in CRC cells (35). Similarly, others have demonstrated that CXCR4 plays a critical role in the promotion of the progression of inflammatory CRC (36). It is commonly known that expression of VEGF-1 in CRC is associated with disease localization, stage, and long-term survival (37). We had previously observed suppression of VEGF in a panel of CRC cell lines post-elraglusib treatment and saw a similar suppression of VEGF in the murine responders. Moreover, we noted a decrease in VEGFR2 in murine responders, a protein that is highly expressed in CRC and promotes angiogenesis (38). Finally, CCL21 has been shown to play a role in colon cancer metastasis (39). Since many of the downregulated analytes in responders play a role in EMT, future studies of elraglusib could include metastatic CRC models.

We observed that responders had higher serum concentrations of CCL4, TWEAK, GM-CSF, CCL22, and IL-12p70 as compared to non-responders. Others have demonstrated that CCL4 is an important chemokine in the TME in determining response to ICB and that a lack of CCL4 can lead to the absence of CD103+ dendritic cells (DCs) (40). DCs are an important cell population influencing the response to ICB, and although we did not monitor their levels in this study, it is conceivable that they played a role in influencing response to therapy. For this reason, further studies could monitor DC populations during the course of therapy. TWEAK is commonly expressed by peripheral blood monocytes and upregulates its expression after exposure to IFN-γ (41). TWEAK has also been shown to promote the nuclear translocation of both classical and alternative NF-κB pathway subunits (42). GM-CSF is a well-known immunomodulatory factor that has immunostimulatory functions but it is also predictive of poor prognosis in CRC (43). Finally, we observed increased levels of IL-12p70 in murine responders. IL-12 is a potent, pro-inflammatory cytokine that has been shown to increase activation and cytotoxicity of both T and NK cells as well as inhibit immunosuppressive cells, such as TAMs (44) and myeloid-derived suppressor cells (MDSCs) (45). We demonstrated that GSK-3 inhibitors such as elraglusib represent a possible combination strategy to increase the efficacy of ICB in patients with MSS CRC. The elraglusib-mediated increase in tumor surface cell-expressed PD-L1 presumably makes this an ideal small molecule to combine with anti-PD-L1 therapies. As this study was concerned solely with CRC, future studies could evaluate the combination of GSK-3 inhibitors with ICB in other malignancies of interest such as pancreatic cancer. This in vivo work was presented, in part, at the 2022 American Association of Cancer Research (AACR) Annual Conference.

Cytokine analysis of plasma samples from patients with refractory solid tumors of multiple tissue origins enrolled in a Phase 1 clinical trial investigating elraglusib (<u>NCT03678883</u>) revealed that elevated baseline plasma levels of proteins such as IL-1 β and reduced levels of proteins such as VEGF correlated with improved PFS and OS. PFS was also found to be positively correlated with elevated plasma levels of immunostimulatory analytes such as Granzyme B, IFN-γ, and IL-2 at 24 hours post-treatment with elraglusib. Several of these secreted proteins correlated with results from the in vivo study where expression of proteins such as IL-1 β, CCL22, CCL4, and TWEAK was positively correlated with improved response to therapy while expression of proteins such as BAFF and VEGF negatively correlated with response to therapy. These results introduce novel circulating biomarkers for correlations with response to therapy which could provide significant clinical utility.

DSP analysis of paired FFPE tumor biopsies from patients with CRC or pancreatic cancer before and after treatment revealed that CD39 expression in PanCK+ segments was negatively correlated with duration of treatment while CD163 expression in CD45+ segments was positively correlated with duration of treatment and potential therapeutic benefit. It is known that CD39 can inhibit costimulatory signaling, increase immunosuppression during T cell priming, and its expression is associated with TAMs, Tregs, and inhibited cytotoxic immune cell function (46). CD39 has been shown to suppress pyroptosis, impair immunogenic cell death, and CD39 expression on endothelial cells regulates the migration of immune cells and promotes angiogenesis (46). Moreover, CD163 is a marker of cells from the monocyte/macrophage lineage therefore future studies could evaluate the impact of monocyte/macrophages on response to elraglusib. We also noted that immune cell segments showed differential protein expression based on the proximity to the tumor where tumor-infiltrating immune cells had decreased expression of immune checkpoints (PD-L1, Tim-3, PD-1) and Treg markers (CD25, CD127) as compared to tumor-adjacent immune cells regardless of timepoint. While the downregulation of immune checkpoint proteins PD-1, TIGIT, and LAG-3 by elraglusib has been previously described in melanoma models (47), our findings regarding VISTA and PD-L2 have not yet been reported. These novel observations regarding emerging immune checkpoint inhibitors should be included in future correlative studies regarding GSK-3 inhibition.

When we analyzed differential protein expression between Long Tx patients and Short Tx patients, we found that Long Tx patients had lower post-treatment expression of mature B cell/DC marker CD35, antigen NY-ESO-1, antigen Her2, antigen MART1, cytotoxic T cell marker CD8, Treg marker Foxp3, antigen PTEN, DC/myeloid marker CD11c, memory T cell marker CD45RO, checkpoint PD-L1, and PR in PanCK+ segments as compared to Short Tx patients which introduces several novel potential biomarkers of response to GSK-3 therapy which should be validated in further studies. Moreover, when we compared post-treatment protein expression in tumor-infiltrating CD45+ immune cell segments in Long Tx patients and Short Tx patients and found that Long Tx patients had decreased expression of antigens NY-ESO-1, PTEN, and PR as compared to Short Tx patients. Interestingly, these three antigens (NY-ESO-1, PTEN, and PR) had decreased expression in Long Tx patients post-treatment regardless of tumor or immune cell region.

There are several potential limitations of this study. One such limitation is that we tested the combination of elraglusib and ICB therapy in a mouse model using only one MSS CRC cell line. Future studies could determine how other MSS CRC cell lines, and perhaps MSI-H cell lines, will respond to this combination treatment. We also had sample size limitations for the number of mice that were included in each treatment group at each flow cytometry timepoint throughout the course of the study, due to the feasibility of the mouse work. Future studies could include larger numbers of mice per flow cytometry timepoint as well as include the comparison of both male and female mice to determine if there are any sex-specific effects. Furthermore, given access to an expanded cohort of tumor biopsies from patients treated with elraglusib, it would be interesting to analyze pre-treatment biopsies between responders and non-responders using DSP technology to aid in identifying predictive biomarkers.

In conclusion, this work demonstrates that small-molecule inhibition of GSK-3 using elraglusib may be a potential means to increase the efficacy of ICB and improve response in patients with MSS CRC, and possibly other tumor types. These findings support further studies and clinical development of elraglusib in combination with ICB, anti-PD-L1 therapy in particular. Moreover, this study, to our knowledge, represents the first digital spatial analysis of tumor biopsies from patients treated with elraglusib and very few oncology drugs have been evaluated using GeoMx technology to date. The novel circulating biomarkers of response to GSK-3 inhibition identified using the cytokine profiling data could provide significant clinical utility and the spatial proteomics data gives us novel insights into the immunomodulatory mechanisms of GSK-3 inhibition.

## Methods

### Cell culture maintenance

Human CRC cells SW480 (RRID: CVCL_0546), HCT-116 (RRID: CVCL_0291), HT-29 (RRID: CVCL_0320), and KM12C (RRID: CVCL_9547) were used in this study. SW480 cells were cultured in Dulbecco’s Modified Eagle Medium (DMEM) supplemented with 10% FBS and 1% Penicillin-Streptomycin HCT-116 and HT-29 were cultured in McCoy’s 5A (modified) Medium supplemented with 10% FBS and 1% Penicillin-Streptomycin. KM12C cells were cultured in Eagle’s Minimal Essential Medium supplemented with 10% FBS and 1% Penicillin-Streptomycin. Human immune cells NK-92 (RRID: CVCL_2142), TALL-104 (RRID: CVCL_2771), and patient-derived CD8+ T cells were also used in this study. NK-92 cells were cultured in Alpha Minimum Essential medium supplemented with 2 mM L-glutamine, 1.5 g/L sodium bicarbonate, 0.2 mM inositol, 0.1 mM 2-mercaptoethanol, 0.02 mM folic acid, 12.5% horse serum, and 12.5% FBS. TALL-104 cells (CD2 +; CD3 +; CD7 +; CD8 +; CD56 +; CD4 −; CD16 −) and patient-derived T cells (CD3 +; CD8 +) were cultured in RPMI-1640 containing 20% FBS, 100 U/ml penicillin, and 100 μg/ml streptomycin. Recombinant human IL-2 (Miltenyi cat# 130-097744) with a final concentration of 100 units/mL was added to all immune cell culture media. All cell lines were incubated at 37°C in a humidified atmosphere containing 5% CO2. Cell lines were authenticated and tested to ensure the cultures were free of mycoplasma infection.

### Measurement of cell viability

Cells were seeded at a density of 3 × 10^3^ cells per well in a 96-well plate (Greiner Bio-One, Monroe, NC, USA). Cell viability was assessed using the CellTiter Glo assay (Promega, Madison, WI, USA). Cells were mixed with 25 μL of CellTiter-Glo reagents in 100 μL of culture volume, and bioluminescence imaging was measured using the Xenogen IVIS imager (Caliper Life Sciences, Waltham, MA). The percent of cell viability was determined by normalizing the luminescence signal to control wells. Dose–response curves were generated and the half maximal inhibitory concentration (IC-50) was calculated using Graph-Pad Prism (RRID: SCR_002798) version 9.2.0. For IC50 generation, concentrations were log-transformed and data were then normalized to control and a log (inhibitor) versus response (three parameters) test was used.

### Pyroptosis assay

Recombinant Human TNF-α (Cat #300-01A, PeproTech, Rocky Hill, NJ, USA) and Recombinant Human IFN-γ (Cat # 300-02, Peprotech, Rocky Hill, NJ, USA) were purchased for use in western blot analysis while rhTRAIL was generated in-house (48).

| Reagent or Resource | Source | Identifier | Dilution |
| --- | --- | --- | --- |
| Vinculin (E1E9V) XP® Rabbit mAb | Cell Signaling | Cat# 13901 | 1:1000 |
| Anti-GSDMB antibody | Sigma-Aldrich | Cat# HPA052407 | 1:1000 |

### Isolation of donor-derived CD8+ T cells

An Easy Step Human CD8+ T Cell Isolation Kit was used to isolate CD8+ T cells from a donor PBMC sample via negative selection (Cat #, 17913, Stem Cell Technologies, Vancouver, Canada).

### Collection of cell culture supernatants used in cytokine measurements

Cells were plated at 3.5 × 10^4^ cells in a 48-well plate (Thermo Fisher Scientific, Waltham, MA, USA) in complete medium and incubated at 37°C with 5% CO2. At 24 hours after plating, almost all the tumor cells were adherent to the bottom of the flask and the complete medium was replaced with the drug-containing medium. Subsequently, the culture supernatants were collected after 48 hours of incubation and were frozen at −80°C until the measurement of cytokines was performed. On the day of analysis, samples were thawed and centrifuged to remove cellular debris.

### Human cytokine profiling

Human cell line culture supernatants were analyzed using an R&D systems Human Premixed Multi-Analyte Kit (R&D Systems, Inc., Minneapolis, MN, USA) and a Luminex 200 (RRID: SCR_018025) Instrument (LX200-XPON-RUO, Luminex Corporation, Austin, TX, USA) according to the manufacturer’s instructions. Sample levels of TNF-α, 4-1BB/TNFRSF9/CD137, IL-8/CXCL8, Ferritin, IFN-β, IL-10, CCL2/JE/MCP-1, VEGF, CXCL13/BLC/BCA-1, IFN-γ, CCL20/MIP-3 α, CCL3/MIP-1 α, CCL22/MDC, CCL4/MIP-1 β, Fas Ligand/TNFSF6, IL-17/IL-17A, IL-2, BAFF/BLyS/TNFSF13B, GM-CSF, CXCL5/ENA-78, TRANCE/TNFSF11/RANK L, CXCL9/MIG, G-CSF, IFN-γ R1/CD119, VEGFR3/Flt-4, C-Reactive Protein/CRP, CXCL11/I-TAC, IL-21, CXCL14/BRAK, IL-6, Fas/TNFRSF6/CD95, TRAIL R3/TNFRSF10C, IL-4, CCL5/RANTES, PD-L1/B7-H1, CCL7/MCP-3/MARC, Chitinase 3-like 1, CXCL10/IP-10/CRG-2, IL-1 β/IL-1F2, IL-7, Prolactin, CCL8/MCP-2, TRAIL R2/TNFRSF10B, M-CSF, IL-15, Granzyme B, IFN-α, TREM-1, IL-12/IL-23 p40, TRAIL/TNFSF10, CCL11/Eotaxin, and IL-18/IL-1F4. Quantitative analysis with 6 standards and a minimum of 50 counts per bead region was used with the Luminex to generate analyte values reported as picograms/ milliliter (pg/mL). Sample concentrations less than the lower limit of detection for each particular analyte were recoded as the lower limit value divided by ten. Sample concentrations above the upper limit of detection for a particular analyte were recoded as the upper limit of detection.

### Murine cytokine profiling

Whole blood from mice was collected, allowed to clot, and serum was isolated using a serum separator tube (SST) according to manufacturer instructions. Murine serum samples were analyzed using an R&D systems Murine Premixed Multi-Analyte Kit (R&D Systems, Inc., Minneapolis, MN, USA) and a Luminex 200 (RRID: SCR_018025) Instrument (LX200-XPON-RUO, Luminex Corporation, Austin, TX, USA) according to the manufacturer’s instructions. Sample levels of GM-CSF, IL-7, IL-12 p70, CCL2/JE/MCP-1, IL-1 β/IL-1F2, VEGF, IL-2, IL-4, VEGFR2/KDR/Flk-1, IL-6, IL-10, IL-13, IFN-γ, IL-3, IL-16, CXCL10/IP-10/CRG-2, CCL5/RANTES, CCL7/MCP-3/MARC, CCL12/MCP-5, Prolactin, M-CSF, CCL3/MIP-1 α, IL-1 α/IL-1F1, CCL20/MIP-3 α, CCL4/MIP-1 β, TWEAK/TNFSF12, CXCL12/SDF-1 α, BAFF/BLyS/TNFSF13B, Granzyme B, CCL21/6Ckine, CCL11/Eotaxin, and CCL22/MDC. Sample values are reported in picograms per milliliter (pg/mL). Quantitative analysis with 6 standards and a minimum of 50 counts per bead region was used with the Luminex to generate analyte values reported as picograms/ milliliter (pg/mL). Sample concentrations less than the lower limit of detection for each particular analyte were recoded as the lower limit value divided by ten. Sample concentrations above the upper limit of detection for a particular analyte were recoded as the upper limit of detection. Data analysis and visualization were generated using R (RRID: SCR_001905) software (R Development Core Team, 2020).

### GFP+ cell line generation

50,000 HT-29 or HCT 116 cells were seeded in a 12-well tissue culture plate and allowed to adhere overnight. They were then transduced with lentivirus containing the plasmid pLenti_CMV_GFP_Hygro [pLenti CMV GFP Hygro (656-4) was a gift from Eric Campeau & Paul Kaufman (Addgene viral prep # 17446-LV; RRID: Addgene_17446)] at a multiplicity of infection of 10 with 8 μg/mL polybrene (hexadimethrine bromide [Cat # 107689, Sigma Aldrich, St. Louis, MO, USA) for 24 hours before washing with PBS and replacing with fresh medium (49). The cells were then sorted for GFP-positivity using a BD FACSAria™ III Cell Sorter (RRID: SCR_016695).

### Multicolor immune cell co-culture experiments

10,000 HCT-116, SW480, or HT-29 cells were plated per well in a clear-bottom, black-walled 48-well tissue culture plate and were allowed to adhere overnight. Cells were subsequently treated with DMSO, 5 μM or 10 μM elraglusib, and/or 10,000 TALL-104 or NK-92 cells (for an effector-to-tumor ratio of 1:1) for 24 hours. CRC cells were labeled using CellTracker^TM^ Green CMFDA (5-chloromethylfluorescein diacetate), immune cells (NK-92, TALL-104) were labeled using CellTracker™ Blue CMAC Dye (7-amino-4-chloromethylcoumarin), and ethidium homodimer-1 (EthD-1) was used as a marker of cell death (Invitrogen, Waltham, MA). 10X images were captured using a Nikon Ti-U Inverted Fluorescence Microscope and NIS-Elements F Package imaging software 3.22.00 Build 710 (Nikon Instruments Inc, USA). The number of red/green color cells in random fields was determined using thresholding and particle analysis in the Fiji modification (RRID: SCR_002285) of ImageJ and expressed as a dead/live cell ratio. Normalization was carried out by subtracting the percentage of cell death due to drug or vehicle control (DMSO) only from the percentage of dead cells observed in the co-culture of tumor and immune cells treated with the drug. At least 100 cells were evaluated per sample, with 3 independent replicates. Statistical analysis was done using GraphPad Prism 9 (RRID: SCR_002798).

### Single-color immune cell co-culture experiments

5000 HT-29 GFP+ or HCT 116 GFP+ cells were plated per well in a clear-bottom, black-walled 96-well tissue culture plate and were allowed to adhere overnight. Cells were subsequently treated with DMSO, 5 μM elraglusib, and/or 5000 TALL-104 or NK-92 cells (for an effector-to-tumor ratio of 1:1) for 48 hours. Nine images were taken per well at 10X magnification using a Molecular Devices ImageXpress® Confocal HT.ai High-Content Imaging System and quantified for the number of GFP+ objects using the MetaXpress (RRID: SCR_016654) software (Molecular Devices, San Jose, CA, USA). 40X Images were also taken at 24 hours for representative images of cellular morphology changes. Statistical analysis was done using GraphPad Prism 9 (RRID: SCR_002798).

### Generation of single-cell suspensions

Spleens were strained, filtered, and washed while tumors were collected, washed, and digested before lymphocytes were collected using a Percoll gradient (Cat # P1644-100ML, Sigma Aldrich, St. Louis, MO).

### Flow cytometry

Flow cytometry viability staining was conducted by suspending murine spleen and tumor single cell suspensions in Zombie Violet fixable viability kit (Cat # 423114, BioLegend, San Diego, CA, USA) according to manufacturer instructions for 30 minutes at room temperature. Staining for membrane surface proteins was conducted using conjugated primary antibodies for 1 hour on ice, according to manufacturer instructions. Cells were fixed and permeabilized using the eBioscience™ Foxp3/Transcription Factor Staining Buffer Set according to manufacturer instructions (Cat# 00-5523-00, Invitrogen, Waltham, MA). Cells were resuspended in Flow Cytometry Staining Buffer (R&D Systems, Minneapolis, MN, USA) and were analyzed using a BD Biosciences LSR II (RRID: SCR_002159) and FlowJo (RRID: SCR_008520) version 10.1 (FlowJo, Ashland, OR, USA).

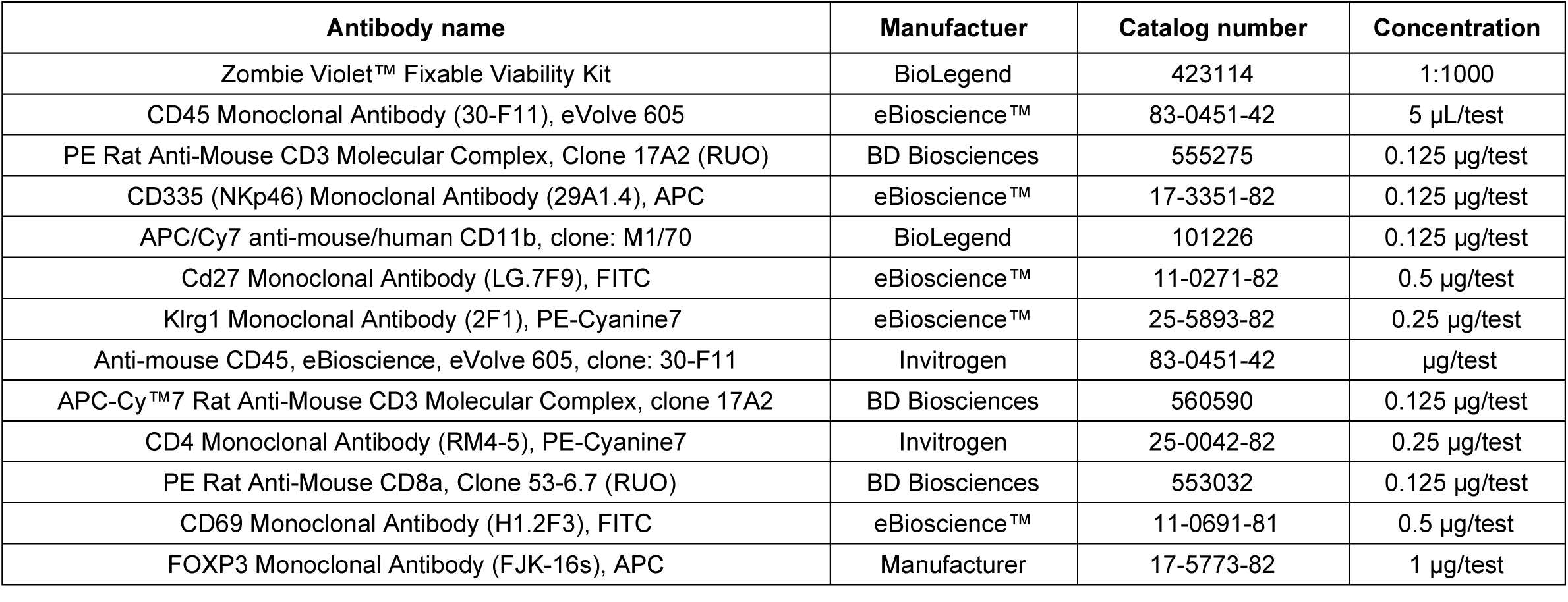

### Natural killer cell immunophenotyping

The NK cell flow cytometry panel included the following directly-conjugated primary antibodies: Anti-mouse CD45, eBioscience eVolve 605 clone: 30-F11 (Ref # 83-0451-42, Invitrogen), PE anti-mouse CD3 molecular complex (17A2) (mat. #: 555275, BD biosciences), Anti-mouse NKp46 APC (Ref # 17-3351-82), APC/Cy7 anti-mouse/human CD11b clone: M1/70 (cat# 101226, BioLegend), anti-Cd27 Monoclonal Antibody (LG.7F9) FITC (eBioscience™, Thermo Scientific, cat # 11-0271-82), and (Klrg1 Monoclonal Antibody (2F1) PE-Cyanine7 (eBioscience, Thermo Scientific, cat # 25-5893-82). Gating strategies are as follows:

NK cell: live/CD45/CD3-/NK1.1+
Mature NK cell: live/CD45/CD3-/NK1.1+/ KRLG1+
Activated NK cell: live/CD45/CD3-/NK1.1+/CD11b+
NK cell subset 1: live/CD45/CD3-/NK1.1+/ CD11b-CD27-
NK cell subset 2: live/CD45/CD3-/NK1.1+/ CD11b-CD27+
NK cell subset 3: live/CD45/CD3-/NK1.1+/ CD11b+CD27+
NK cell subset 4: live/CD45/CD3-/NK1.1+/ CD11b+CD27-

### T cell immunophenotyping

The T cell flow cytometry panel included the following directly-conjugated primary antibodies: Anti-mouse CD45 superbright 600 clone: 30-511 (ref# 63-0451-82, eBioscience), anti-CD3 APC-Cy7 clone 17A2(BD Biosciences, cat # 560590), eBioscience anti-mouse CD4 PE-Cy7 clone: RM4-5 (Ref # 25-0042-82, Invitrogen), PE anti-mouse CD8a (Ly-2)(53-6.7) (cat # 553032, BD), Anti-mouse CD69 FITC clone: H1.2F3 (Ref# 11-0691-81, eBioscience), and Foxp3 (FJK-16s) APC (eBioscience). Gating strategies are as follows:

CD4+ T cell: live/CD45+/CD3+/CD4+/Foxp3-
CD8+ T cell: live/CD45+/CD3+/CD8+
Treg: live/CD45+/CD3+/CD4+/Foxp3+
Activated CD8+ T cell: live/CD45+/CD3+/CD8+/CD69+

### Western blot analysis

Cells were plated in a 6-well plate and incubated overnight before the spent media was replaced with drugged media. Drug treatment lasted for indicated durations. Protein was extracted using radioimmunoprecipitation (RIPA) assay buffer (Cat # R0278, Sigma-Aldrich, St. Louis, MO) containing cOmplete™, Mini, EDTA-free Protease Inhibitor Cocktail (Cat # 4693159001, Roche, Basel, Switzerland) from sub-confluent cells. Denaturing sample buffer was added, samples were boiled at 95 degrees for 10 minutes, and an equal amount of protein lysate was electrophoresed through NuPAGE™ 4 to 12%, Bis-Tris, 1.5 mm, Mini Protein Gels (Invitrogen, Waltham, MA) then transferred to PVDF membranes. The PVDF membrane was blocked with 5% non-fat milk (Sigma-Aldrich, St. Louis, MO) in 1x TTBS. Primary antibodies were incubated with the transferred PVDF membrane in blocking buffer at 4°C overnight. Secondary antibodies used included Goat anti-Rabbit IgG (H + L) Secondary Antibody, HRP (Cat # 31460, Invitrogen, Waltham, MA), and Goat anti-Mouse IgG (H + L) Secondary Antibody, HRP (Cat # 31430, Invitrogen, Waltham, MA). Signal was detected using Pierce™ ECL Western Blotting Substrate (Cat # 32106, Thermo Scientific, Waltham, MA, USA) and a Syngene Imaging System (RRID: SCR_015770).

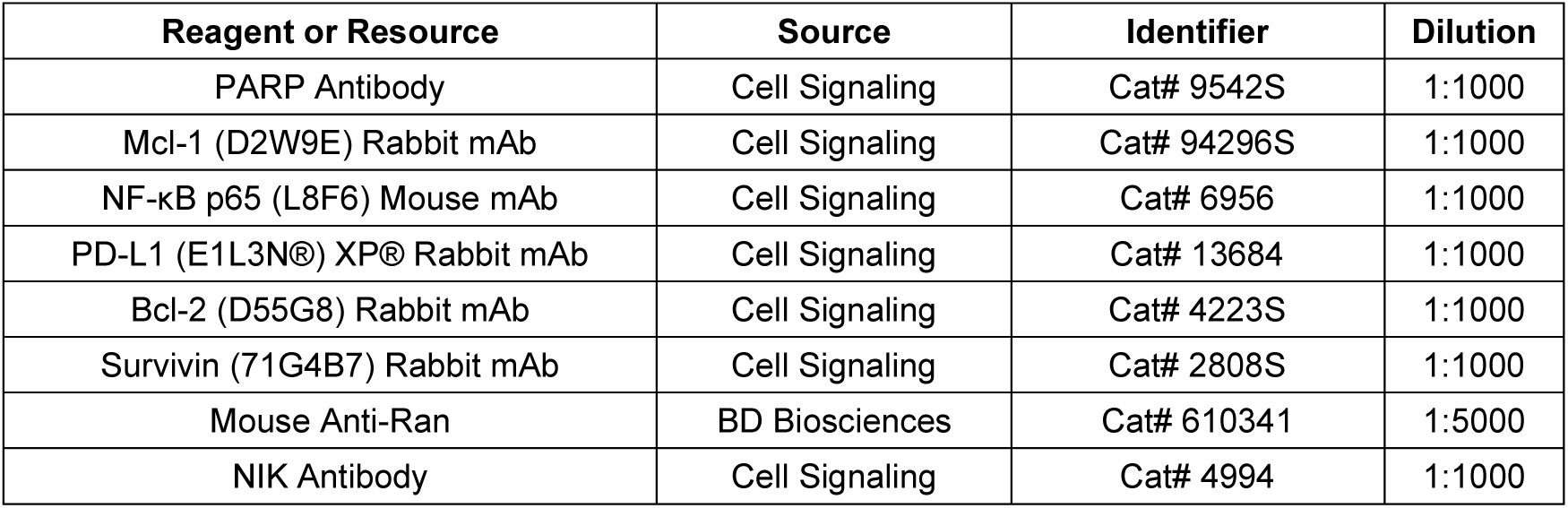

| Reagent or Resource | Source | Identifier | Dilution |
| --- | --- | --- | --- |
| PARP Antibody | Cell Signaling | Cat# 9542S | 1:1000 |
| Mcl-1 (D2W9E) Rabbit mAb | Cell Signaling | Cat# 94296S | 1:1000 |
| NF-κB p65 (L8F6) Mouse mAb | Cell Signaling | Cat# 6956 | 1:1000 |
| PD-L1 (E1L3N®) XP® Rabbit mAb | Cell Signaling | Cat# 13684 | 1:1000 |
| Bcl-2 (D55G8) Rabbit mAb | Cell Signaling | Cat# 4223S | 1:1000 |
| Survivin (71G4B7) Rabbit mAb | Cell Signaling | Cat# 2808S | 1:1000 |
| Mouse Anti-Ran | BD Biosciences | Cat# 610341 | 1:5000 |
| NIK Antibody | Cell Signaling | Cat# 4994 | 1:1000 |

### In vivo studies

The experimental in vivo protocol (Protocol # 19-01-003) was approved by the Institutional Animal Care and Use Committee of Brown University (Providence, RI, USA). Six to 7 weeks-old female BALB/c mice (RRID: IMSR_JAX:000651) were purchased from Taconic. 50,000 cells were suspended in 50 µL ice-cold PBS and 50 µL Matrigel (Catalog # 354234, Corning, New York, USA), and 100 uL was injected subcutaneously into the rear flanks. Once tumor volume reached at least 100 mm^3^, mice were randomly assigned to one of seven groups (3 mice/group): Control (isotype), elraglusib, elraglusib + Isotype, anti-PD-1, anti-PD-L1, elraglusib + anti-PD-1, and elraglusib + anti-PD-L1. All treatments were delivered by IP injection on the following dosing schedule: Isotype (70 mg/kg, twice a week), elraglusib (70 mg/kg, twice a week), anti-PD-1 (10 mg/kg, twice a week), anti-PD-L1 (10 mg/kg, twice a week). The treatment continued until mice developed signs of discomfort from excessive tumor growth. Mice were weighed once a week to monitor signs of drug toxicity. The length (L) and width (W) of the masses were measured three times per week with a digital caliper, and the tumor volume was calculated by applying the formula: 0.5LW2. Collection of whole blood and serum was performed by cardiac puncture and sent to Antech GLP for blood cell count and chemistry tests, or in-house cytokine profiling. Tumors and organs were dissected and harvested for analysis by IHC and flow cytometry.

### Immunohistochemistry

Excised tissues are fixed with 10% neutral buffered formalin and paraffin-embedded. 5-micrometer tissue sections are cut with a microtome and mounted on glass microscope slides for staining. Hematoxylin and eosin staining was completed for all tumor specimens. Paraffin embedding and sectioning of slides were performed by the Brown University Molecular Pathology Core Facility. Slides were dewaxed in xylene and subsequently hydrated in ethanol at decreasing concentrations. Antigen retrieval was carried out by boiling the slides in 2.1 g citric acid (pH 6) for 10 minutes. Endogenous peroxidases were quenched by incubating the slides in 3% hydrogen peroxide for 5 minutes. After nuclear membrane permeabilization with Tris-buffered saline plus 0.1% Tween 20, slides were blocked with horse serum (Cat# MP-7401-15, Vector Laboratories, Burlingame, CA, USA), and incubated with primary antibodies overnight (Supplementary Table S1) in a humidified chamber at 4C. After washing with PBS, a secondary antibody (Cat# MP-7401-15 or MP-7402, Vector Laboratories, Burlingame, CA, USA) was added for 30 minutes, followed by diaminobenzidine application (Cat# NC9276270, Thermo Fisher Scientific, Waltham, MA, USA) according to the manufacturer’s protocol. Samples were counterstained with hematoxylin, rinsed with distilled water, dehydrated in an increasing gradient of ethanol, cleared with xylene, and mounted with Cytoseal mounting medium (Thermo Fisher Scientific, catalog no. 8312-4). Images were recorded on a Zeiss Axioskop microscope (RRID: SCR_014587), using QCapture (RRID: SCR_014432). QuPath software (RRID: SCR_018257) was used to automatically count positive cells. For each IHC marker, five 20X images per group were analyzed, and results were represented as the absolute number of positive cells per 20X field.

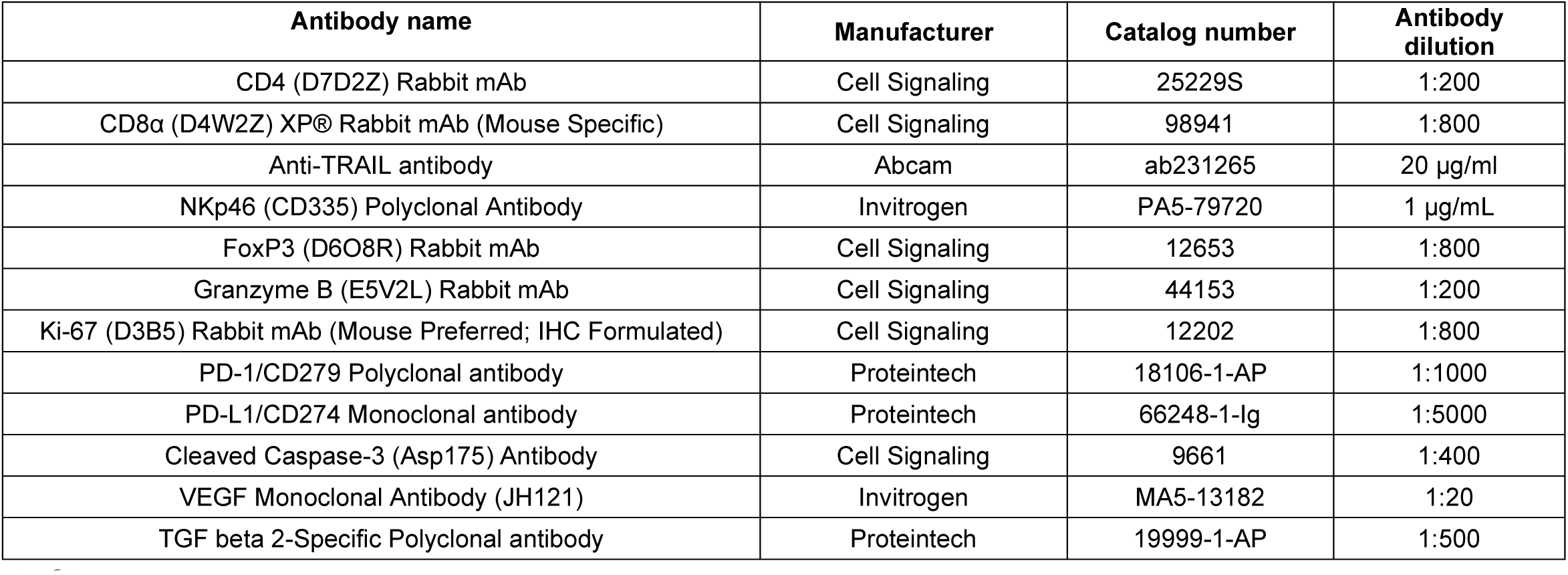

| Antibody name | Manufacturer | Catalog number | Antibody dilution |
| --- | --- | --- | --- |
| CD4 (D7D2Z) Rabbit mAb | Cell Signaling | 25229S | 1:200 |
| CD8α (D4W2Z) XP® Rabbit mAb (Mouse Specific) | Cell Signaling | 98941 | 1:800 |
| Anti-TRAIL antibody | Abcam | ab231265 | 20 µg/ml |
| NKp46 (CD335) Polyclonal Antibody | Invitrogen | PA5-79720 | 1 µg/mL |
| FoxP3 (D6O8R) Rabbit mAb | Cell Signaling | 12653 | 1:800 |
| Granzyme B (E5V2L) Rabbit mAb | Cell Signaling | 44153 | 1:200 |
| Ki-67 (D3B5) Rabbit mAb (Mouse Preferred; IHC Formulated) | Cell Signaling | 12202 | 1:800 |
| PD-1/CD279 Polyclonal antibody | Proteintech | 18106-1-AP | 1:1000 |
| PD-L1/CD274 Monoclonal antibody | Proteintech | 66248-1-Ig | 1:5000 |
| Cleaved Caspase-3 (Asp175) Antibody | Cell Signaling | 9661 | 1:400 |
| VEGF Monoclonal Antibody (JH121) | Invitrogen | MA5-13182 | 1:20 |
| TGF beta 2-Specific Polyclonal antibody | Proteintech | 19999-1-AP | 1:500 |

### Microarrays

A total of 0.5×10^6^ tumor cells (HCT-116, HT-29, KM12C) were plated in a 6-well plate and allowed to adhere overnight before 24-hour treatment as indicated. 1×10^6^ immune cells (NK92, TALL-104) were plated and treated with elraglusib as indicated for 24 hours. RNA was isolated from cell pellets in batches of 6 using an RNeasy Plus Mini Kit (Cat # 74134, Qiagen, Hilden, Germany). Acceptable RNA concentration and quality were verified with Nanodrop and Bioanalyzer measurements. GeneChip™ Human Transcriptome Array 2.0 assays were conducted according to manufacturer instructions in two batches using randomized samples to limit batch effects (Cat# 902162, Applied Biosystems, Waltham, MA, USA). Applied Biosystems Transcriptomic Analysis Console (TAC) software (RRID: SCR_016519) was used to calculate fold changes in gene expression relative to the untreated control cells. Values were considered statistically significant for p values <0.05.

### Single-cell RNA sequencing

Single cells were captured and 3’ single-cell gene expression libraries were conducted (Next GEM v3.1) using the 10x Genomics Chromium system by SingulOmics (SingulOmics, New York, NY, USA). Gene expression libraries were sequenced with ∼200 million PE150 reads per sample on Illumina (RRID: SCR_016387) NovaSeq (Illumina, Inc., San Diego, CA, USA). After sequencing clean reads were then analyzed with human reference genome GRCh38 using Cell Ranger v6.1.2 ([RRID: SCR_017344],10X Genomics, Pleasanton, CA, USA). Data were analyzed and visualized using Loupe Browser ([RRID: SCR_018555], 10X Genomics, Pleasanton, CA, USA).

### Digital Spatial Profiling

An Agilent technologies hybridization oven was used for baking tissue onto slides (Agilent, Santa Clara, CA, USA). A NanoString GeoMx® Digital Spatial Profiler (DSP) instrument (NanoString, Seattle, WA, USA) was used to scan slides, identify regions of interest (ROIs), and collect photocleavable barcodes according to manufacturer instructions. A custom panel was designed to include the following proteins: Ms IgG1, Ms IgG2a, Rb IgG, GAPDH, Histone H3, S6, Beta-2-microglobulin, CD31, CD45, Ki-67, ARG1, CD11b, CD11c, CD14, CD163, CD39, CD40, CD68, HLA-DR, GZMB, CD20, CD3, CD34, CD4, CD56, CD66b, CD8, Foxp3, Fibronectin, 4-1BB, B7-H3, CTLA4, GITR, IDO1, LAG3, OX40L, STING, Tim-3, VISTA, Bcl-2, ER-α, EpCAM, Her2, MART1, NY-ESO-1, PR, PTEN, PanCk, SMA, CD127, CD25, CD27, CD44, CD45RO, CD80, ICOS, PD-1, PD-L1, and PD-L2. An Eppendorf MasterCycler Gradient Thermal Cycler was used to generate the Illumina sequencing libraries from the photocleaved tags. (Eppendorf, Hamburg, Germany). An Agilent Fragment Analyzer (RRID: SCR_019417) was used for library size distribution analysis with a high-sensitivity NGS Fragment Kit (Cat# DNF-474-0500, Agilent, Santa Clara, CA, USA). qPCR for quantification was run using an Illumina-compatible KAPA Library Quantification Kits (ROX Low) (cat# KK4873) on an Applied Biosystems ViiA 7 Real-Time qPCR / PCR Thermal Cycler System (Applied Biosystems, San Francisco, CA, USA) and was analyzed using QuantStudio software (RRID: SCR_018712). Sequencing was performed using a NextSeq 500/550 High Output Kit v2.5 (75 Cycles) kit (cat# 20024906) on an Illumina Sequencing NextSeq 550 System ([RRID: SCR_016381], Illumina, San Diego, CA, USA). The initial annotated dataset went through quality control (QC) to check if housekeeper genes and background (isotype) control molecules were themselves correlated with the predictors of interest. Every ROI was tested for raw sequencing reads (segments with <1000 raw reads were removed), % sequencing saturation (defined as [1-deduplicated reads/aligned reads]%, segments below ∼50% were not analyzed), and nuclei count per segment (>100 nuclei per segment is generally recommended). Both immunoglobulins (IgGs) and housekeeper genes were highly correlated with one another. Signal to noise (SNR) ratio was calculated using background probes and all probes were detected above the background in at least one ROI. Finally, data were normalized based on background IgG expression and all normalization factors were well distributed. Data analysis and visualization were generated using R ([RRID: SCR_001905], R Development Core Team, 2020).

### Clinical specimens

Archival tumor specimens and peripheral blood samples were collected from patients enrolled in the Phase I study of Elraglusib (9-ING-41), a small molecule selective glycogen synthase kinase-3 beta (GSK-3b) inhibitor, as monotherapy or combined with cytotoxic regimens in patients with relapsed or refractory hematologic malignancies or solid tumors (Clinicaltrials.gov NCT03678883) who received treatment at the Lifespan Cancer Institute (Providence, RI, USA). The study was conducted in accordance with the Declaration of Helsinki and the International Conference on Harmonization Good Clinical Practice guidelines. The study protocol was approved by the Institutional Review Board (IRB) of Rhode Island Hospital under protocol number 1324888-120. The patients also participated in a Lifespan Cancer Institute research protocol designed to investigate molecular and genetic features of tumors and mechanisms of resistance (Rhode Island Hospital IRB protocol number 449060-38). All patients provided written informed consent.

### Statistical analysis

GraphPad Prism (RRID: SCR_002798) version 9.5.0 was used for statistical analyses and graphical representation (GraphPad, San Diego, CA, USA). Data are presented as means ± standard deviation (SD) or standard error of the mean (SEM). The relations between groups were compared using two-tailed, paired student’s T tests or one-way ANOVA tests. Survival was analyzed with the Kaplan-Meier method and was compared with the log-rank test. For multiple testing, Tukey’s or Benjamini-Hochberg’s methods were employed. Statistical significance is reported as follows: P ≤ 0.05: *, P ≤ 0.01: **, and P ≤ 0.001: ***.

## Data Availability Statement

The microarray data generated in this study are publicly available in Gene Expression Omnibus (GSE222849) at GSE. Other data generated in this study are available within the article and its supplementary data files. Further information and requests for resources and reagents should be directed to and will be fulfilled by the lead contact.

## Institutional review board statement

This study was approved by the Institutional Review Board (IRB) of Rhode Island Hospital under protocol numbers 449060-38 and 1324888-120.

## Acknowledgments

W.S.E-D. is an American Cancer Society Research Professor and is supported by the Mencoff Family University Professorship at Brown University. Research reported in this manuscript was supported by the National Cancer Institute of the National Institutes of Health under award number 1F31CA271636-01 to K.E.H. The contents of this manuscript are solely the responsibility of the authors and do not necessarily represent the official views of the National Cancer Institute, the National Institutes of Health, or the American Cancer Society. This work was supported by the Teymour Alireza P’98, P’00 Family Cancer Research Fund established by the Alireza Family. We thank Dongfang Yang at the Lifespan Molecular Pathology Core Laboratory for assistance with sectioning patient tumor biopsies. We thank the Campbell lab for the NK-92 cell line and we thank Eric Campeau & Paul Kaufman for the pLenti CMV GFP Hygro (656-4) plasmid.

---

**Supplementary Figure 1.**
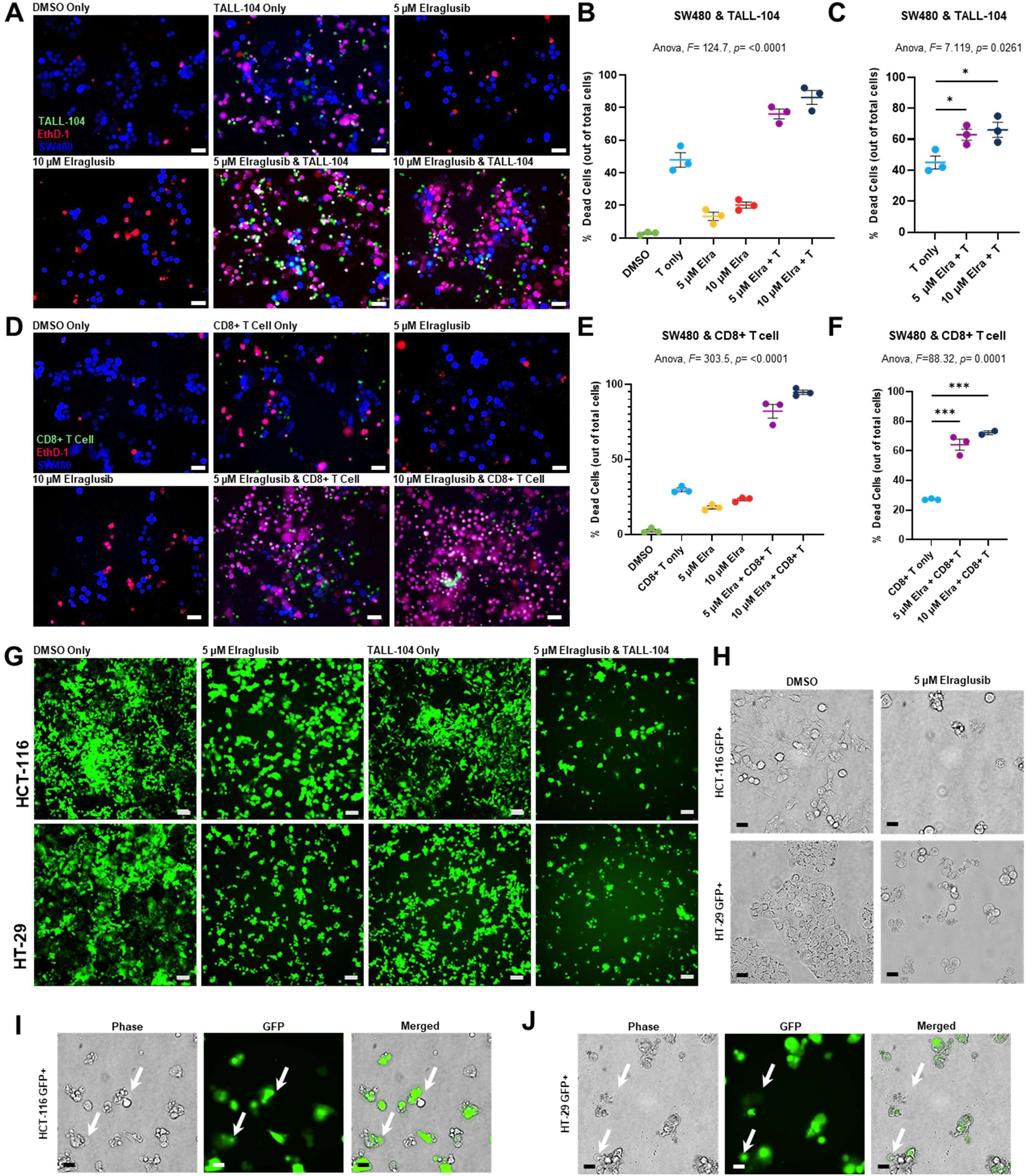
Elraglusib increases immune-mediated cytotoxicity in a co-culture model with CRC cells. **(A)** Co-culture SW480 and TALL-104 T cell co-culture assay images at the 24-hour timepoint. 24-hour tumor cell pre-treatment with 5 µM elraglusib, followed by 24-hour co-culture. EthD-1 was used to visualize dead cells, 10X magnification, scale bar indicates 100 µm. **(B)** Quantification of co-culture experiment using the percentage of dead cells out of total cells (n=3). **(C)** Quantification normalized by cell death observed with drug treatment alone (n=3). **(D)** SW480 and donor-derived CD8+ T cell co-culture assay images at the 24-hour timepoint. **(E)** Quantification of co-culture experiment using the percentage of dead cells out of total cells (n=3). **(F)** Quantification normalized by cell death observed with drug treatment alone (n=3). A one-way ANOVA followed by a post-hoc Dunnett’s multiple comparisons test was used to calculate statistical significance. Statistical significance is reported as follows: p ≤ 0.05: *, p ≤ 0.01: **, and p ≤ 0.001. **(G)** HCT-116 GFP+ or HT-29 GFP+ cells were co-cultured with TALL-104 cells at a 1:1 E:T ratio and were treated with DMSO or 5 μM elraglusib (n=3). **(H)** 40X images were collected after 24 hours of DMSO or 5 μM elraglusib treatment. **(I)** 40X images of a co-culture of HCT-116 GFP+ cells and NK-92 cells at a 1:1 E:T ratio were collected after 36 hours of 5 μM elraglusib treatment. White arrows indicate pyroptotic events. **(J)** 40X images of a co-culture of HT-29 GFP+ CRC cells and NK-92 cells at a 1:1 E:T ratio were after 36 hours of 5 μM elraglusib treatment. White arrows indicate pyroptotic events. P-value legend: * p < 0.05, ** p < 0.01, *** p < 0.001, **** p < 0.0001.

**Supplementary Figure 2.**
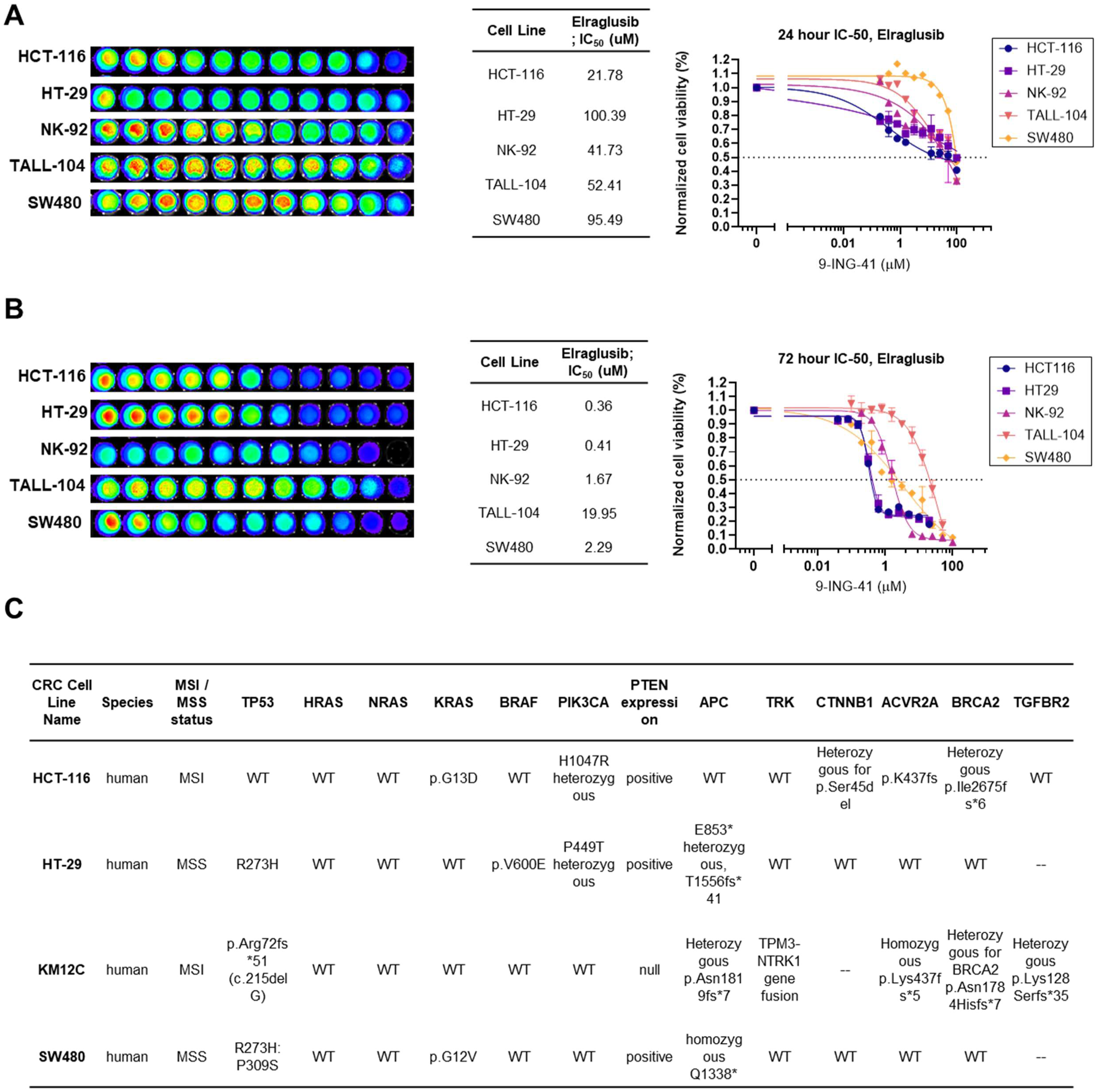
CRC cell lines selected represent diverse mutational backgrounds and exhibit varying elraglusib IC-50 values. Tumor cell lines (HCT-116, HT-29, SW480) and immune cell lines (NK-92, TALL-104) were treated as indicated for **(A)** 24-hour or **(B)** 72-hour cell viability was assessed to determine IC-50 values. **(C)** Table of CRC cell lines included in the study and their diverse mutational profiles.

**Supplementary Figure 3.**
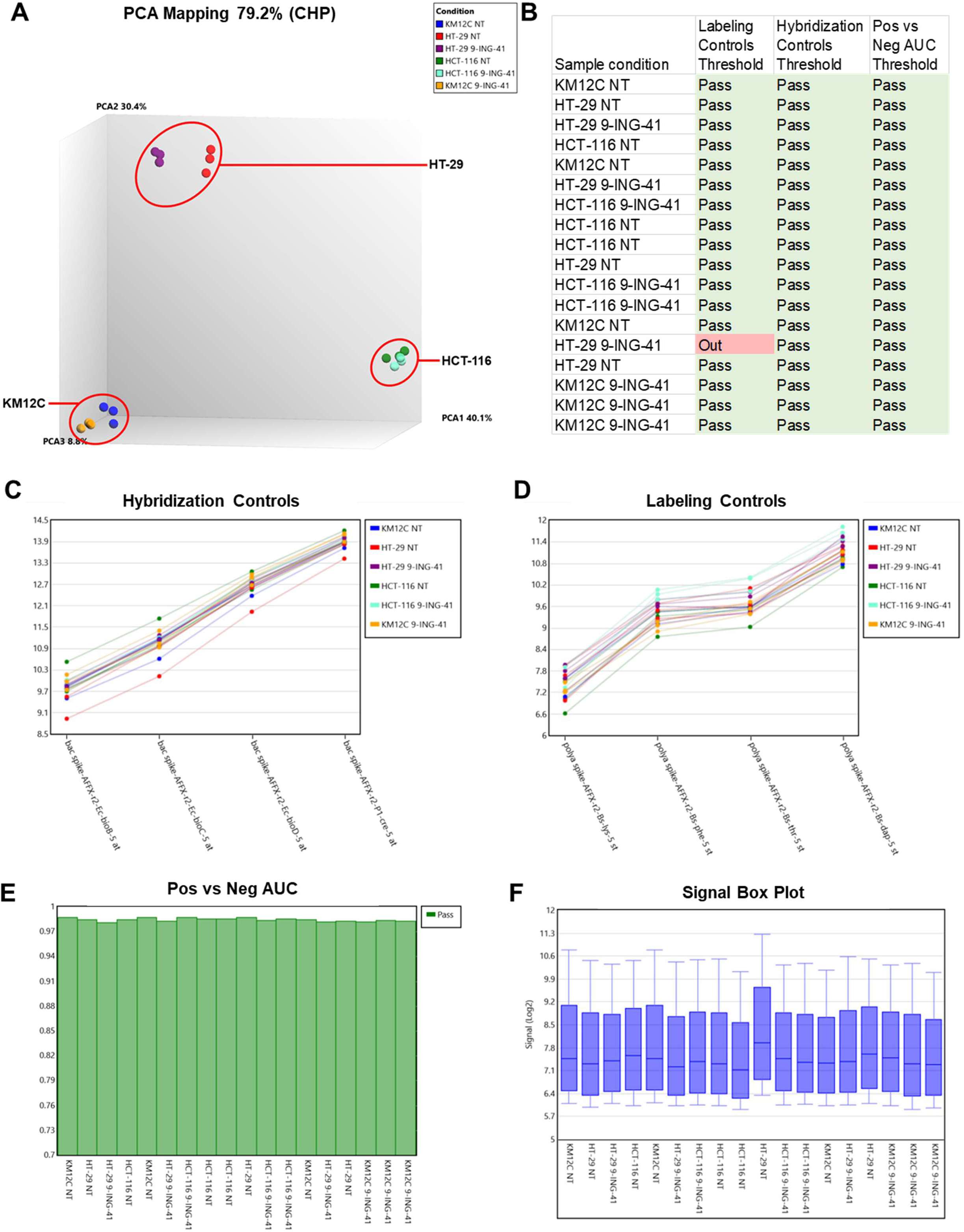
Internal controls for tumor cell microarray analysis. **(A)** PCA mapping demonstrated clear mapping of HCT-116, HT-29, and KM12C cells. **(B)** Quality control was determined satisfactory for further analysis. **(C)** Hybridization controls. **(D)** Labeling controls. **(E)** Pos vs Neg Area Under the Curve (AUC). **(F)** Signal box plot.

**Supplementary Figure 4.**
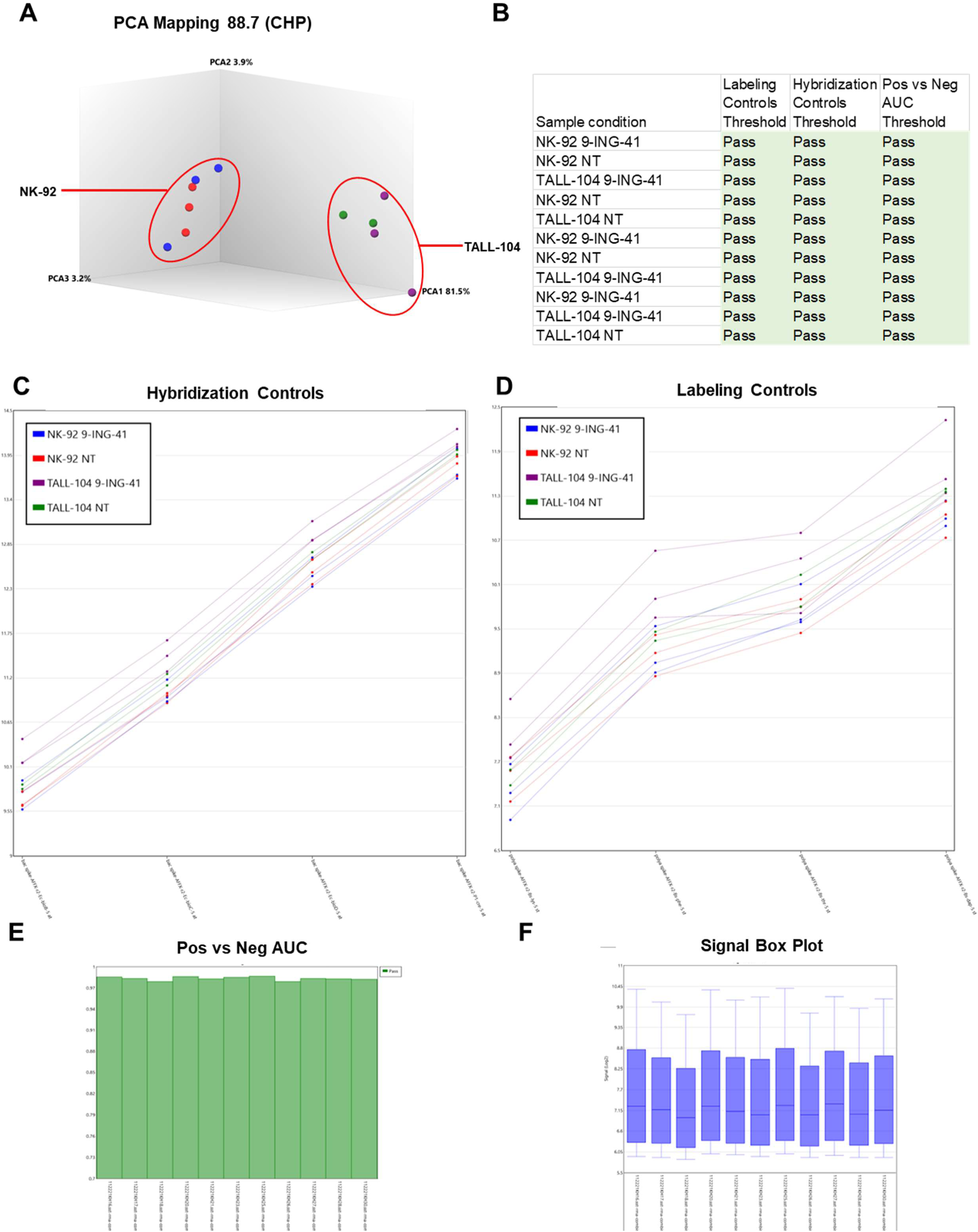
Internal controls for immune cell microarray analysis. **(A)** PCA mapping demonstrated clear mapping of NK-91 and TALL-104 cells. **(B)** Quality control was determined satisfactory for further analysis. **(C)** Hybridization controls. **(D)** Labeling controls. **(E)** Pos vs Neg Area Under the Curve (AUC). **(F)** Signal box plot.

**Supplementary Figure 5.**
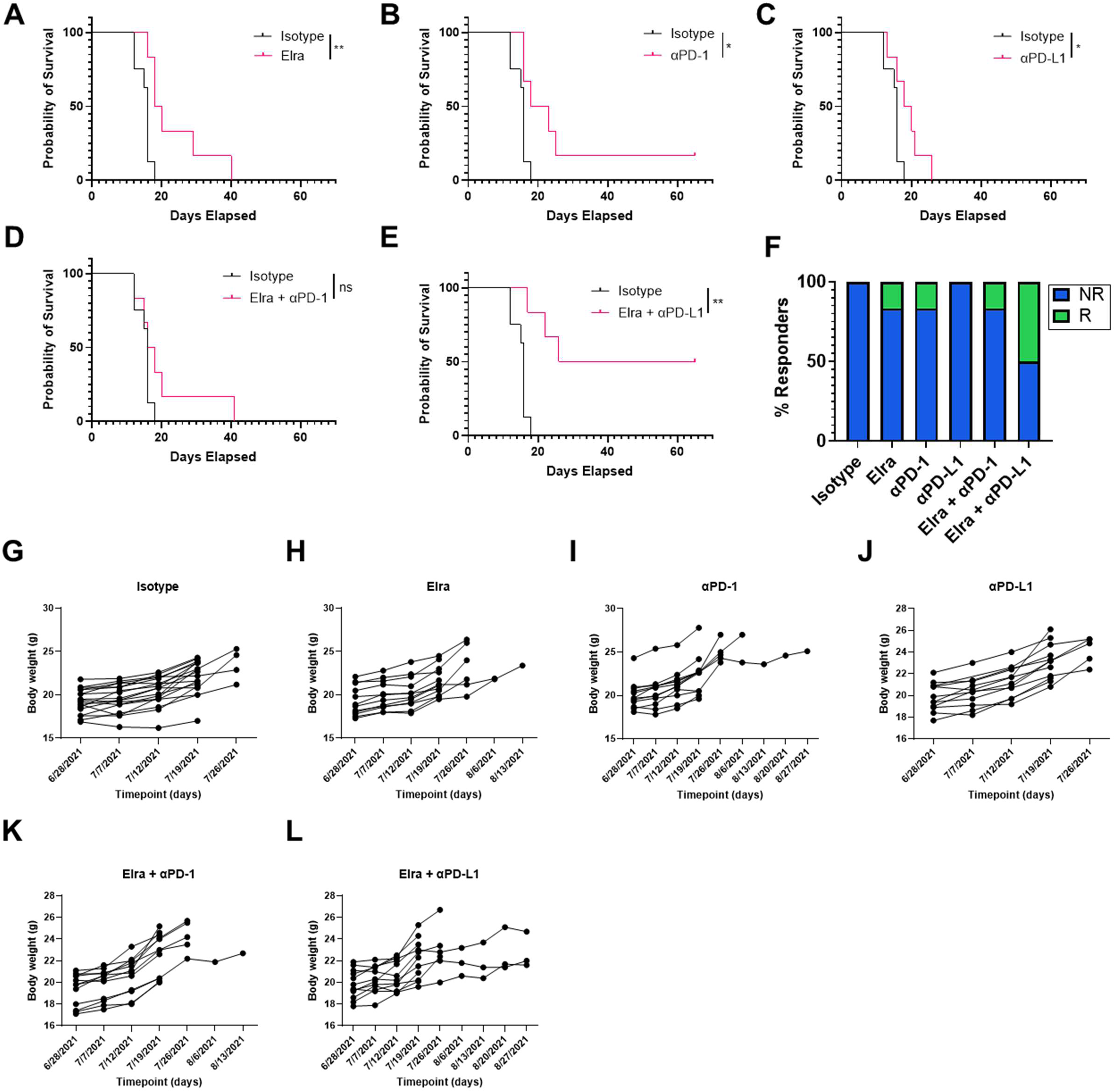
Syngeneic murine colon carcinoma BALB/c murine model with MSS cell line CT-26 Kaplan Meier curves and mouse body weights grouped by treatment. Individual Kaplan Meier curves for isotype control compared to **(A)** elraglusib, **(B)** anti-PD-1, **(C)** anti-PD-L1, **(D)** elraglusib + anti-PD-1, and **(E)** elraglusib + anti-PD-L1. **(F)** Bar graph indicating the percentage of responders **(R)** and non-responders (NR) per treatment group. Individual body weight plots for **(G)** Isotype control, **(H)** elraglusib, **(I)** anti-PD-1, **(J)** anti-PD-L1, **(K)** elraglusib + anti-PD-1, and **(L)** elraglusib + anti-PD-L1. P-value legend: * p < 0.05, ** p < 0.01, *** p < 0.001, **** p < 0.0001.

**Supplementary Figure 6.**
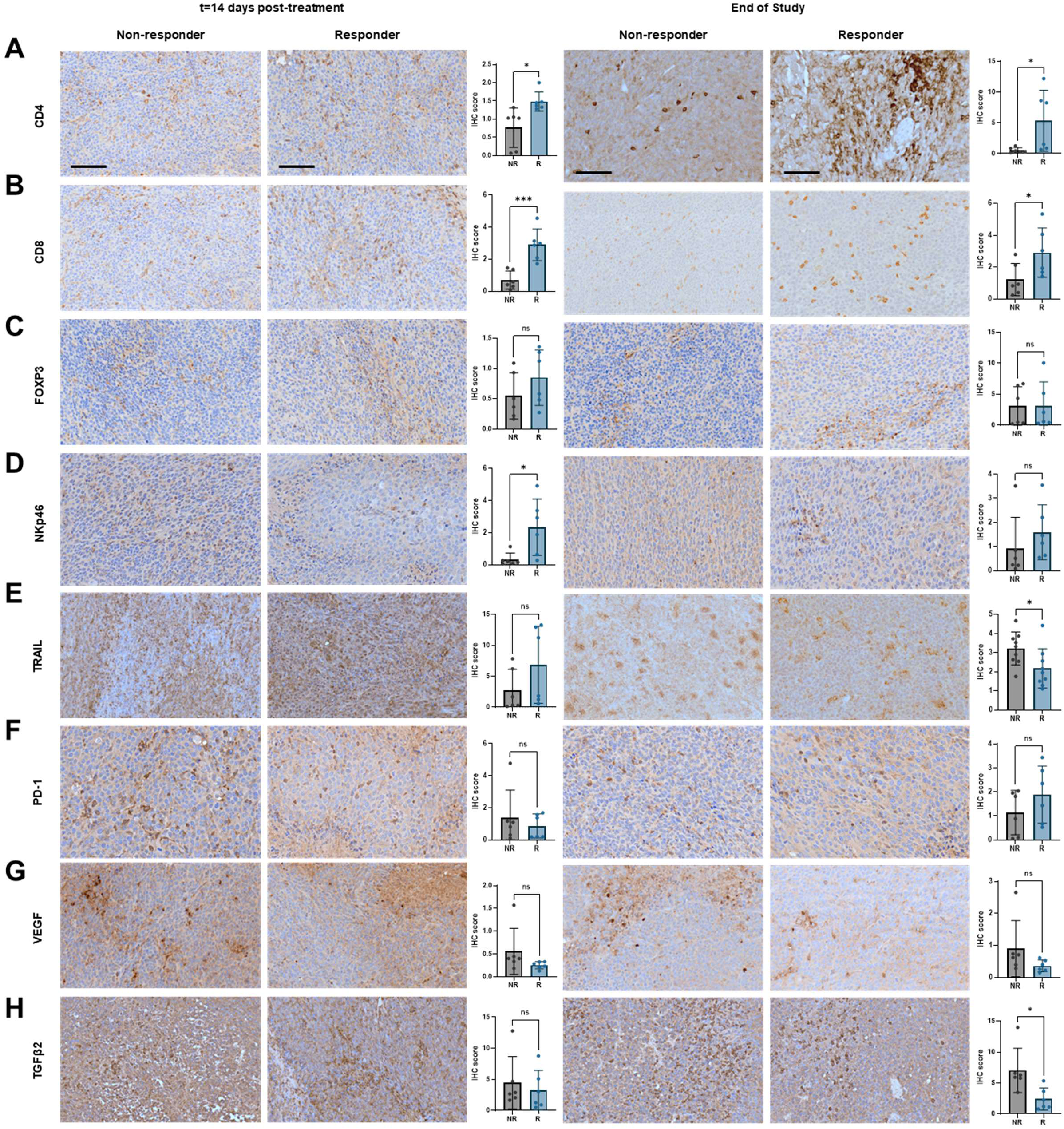
Immunohistochemistry analysis of tumor sections in responders as compared to non-responders. **(A)** CD4, **(B)** CD8, **(C)** Foxp3, **(D)** NKp46, **(E)** TNF-related apoptosis-inducing ligand (TRAIL), **(F)** PD-1, **(G)** Vascular Endothelial Growth Factor (VEGF), and **(H)** Transforming Growth Factor Beta 2 (TGFβ2) were compared at the 14 days post-treatment initiation timepoint and the end-of-study (EOS) timepoint, respectively. Non-responders (NR) and responders **(R)** were compared. Statistical significance was determined using two-tailed unpaired T tests (n=6). 20X images, scale bar represents 100 μm.

**Supplementary Figure 7.**
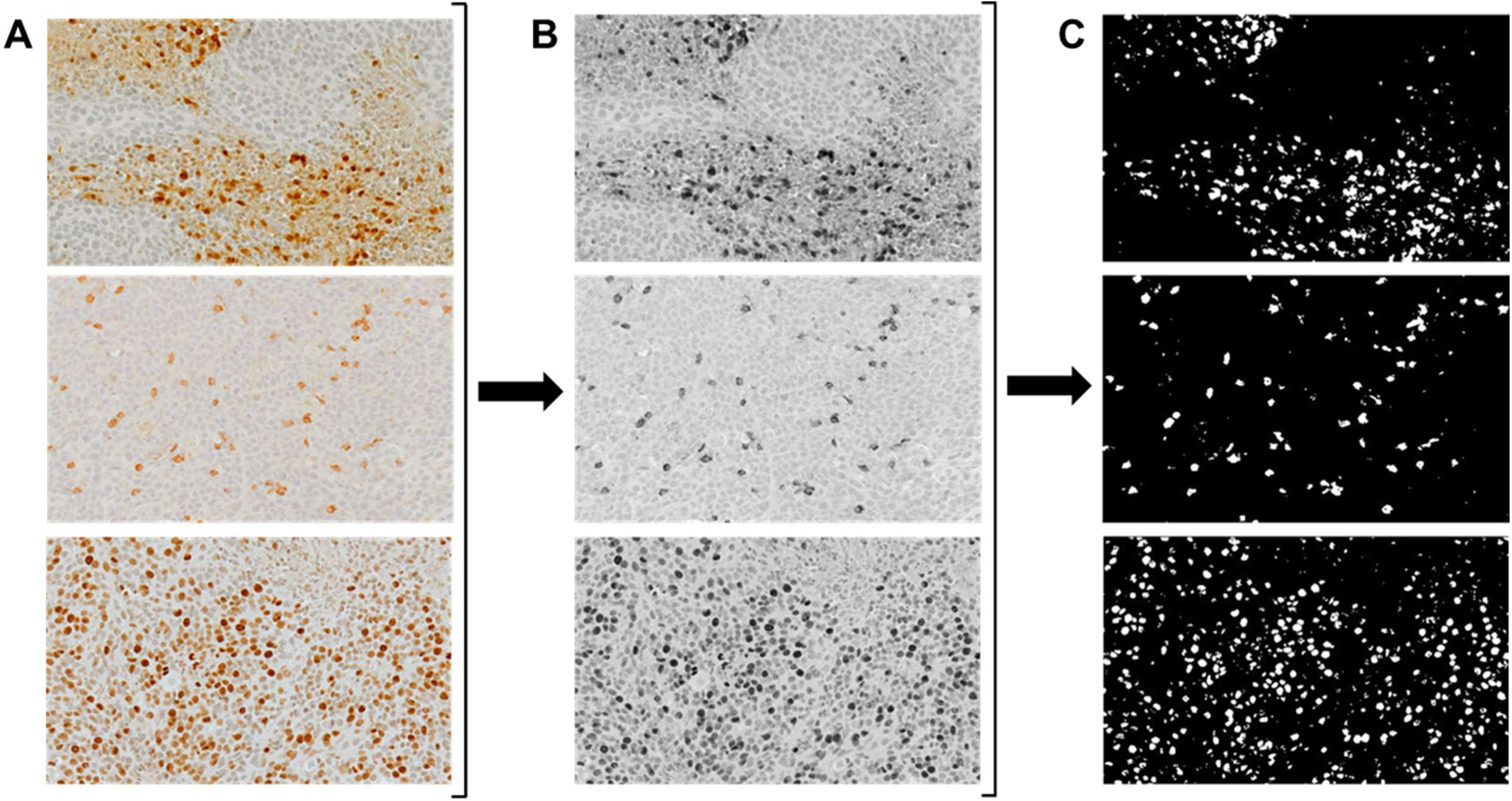
IHC thresholding analysis workflow. **(A)** 20X images were converted to **(B)** 16-bit images and were then analyzed using **(C)** MaxEntropy thresholding. Particles were analyzed and the reported percentage area covered was used to quantify the signal.

**Supplementary Figure 8.**
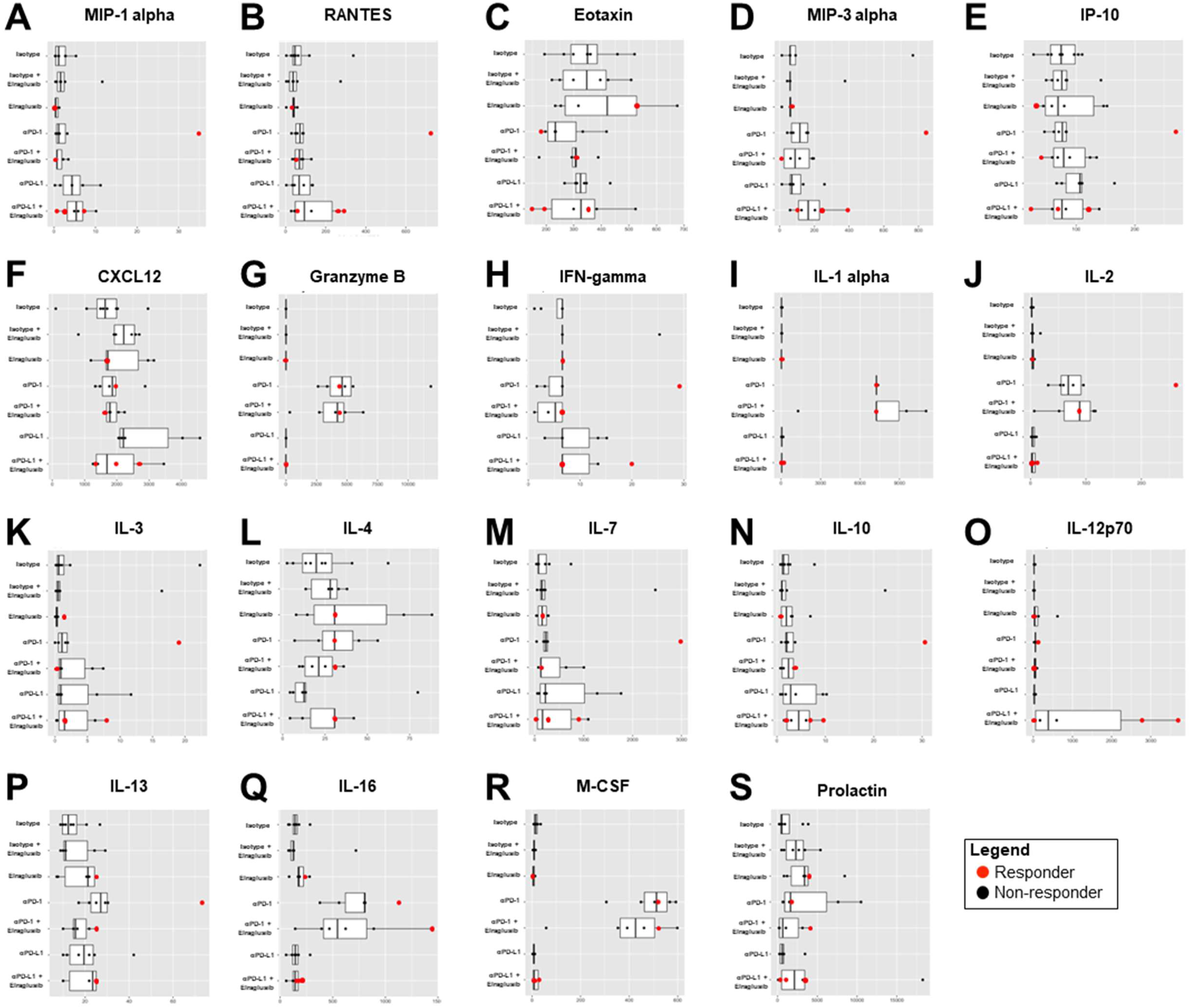
Murine serum cytokine profiling assay results. Serum from end-of-study mice was analyzed via cytokine profiling for **(A)** MIP-1 α, **(B)** RANTES, **(C)** Eotaxin, **(D)** MIP-3 α, **(E)** IP-10, **(F)** CXCL12, **(G)** Granzyme B, **(H)** IFN-γ, **(I)** IL-1 α, **(J)** IL-2, **(K)** IL-3, **(L)** IL-4, **(M)** IL-7, **(N)** IL-10, **(O)** IL-12 p70, **(P)** IL-13, **(Q)** IL-16, **(R)** M-CSF, and **(S)** Prolactin. Responders (red) and non-responders (black) were compared. A Kruskal-Wallis test was used to calculate statistical significance followed by a Benjamini-Hochberg correction for multiple comparisons.

**Supplementary Figure 9.**
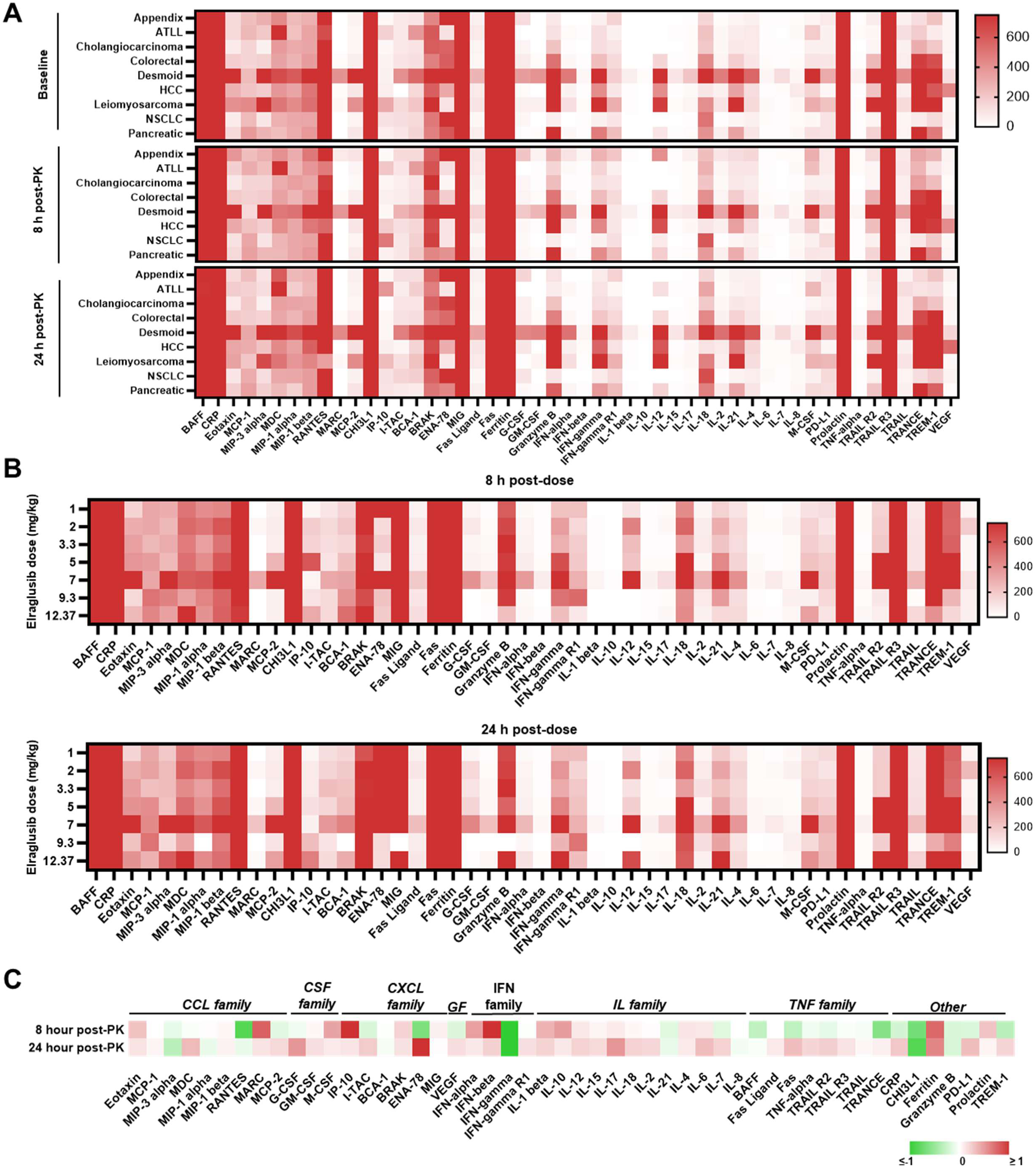
Multiplex immunoassay reveals dynamic cytokine changes 8-and 24-hours post-elraglusib dose. Plasma samples from human patients with refractory solid tumors of multiple tissue origins enrolled in a Phase 1 clinical trial investigating GSK-3 inhibitor elraglusib (NCT03678883) were analyzed using a Luminex 200. **(A)** Raw analyte values are grouped by timepoint: baseline (pre-dose), 8-hour post-PK, and 24-hour post-PK and categorized by primary tumor location: appendix, adult T cell leukemia/lymphoma (ATLL), cholangiocarcinoma, colorectal, desmoid, hepatocellular carcinoma (HCC), leiomyosarcoma, non-small cell lung cancer (NSCLC), and pancreatic. The scale bar is in pg/mL. **(B)** Raw analyte values are grouped by timepoint: 8-hour post-PK and 24-hour post-PK and categorized by elraglusib dose: 1, 2, 3.3, 5, 7, 9.3, or 12.37 mg/kg. **(C)** FC heatmaps grouped by cytokine family. 8-and 24-hours post-PK were compared to baseline (pre-PK) analyte values. Green indicates downregulation and red indicates upregulation.

**Supplementary Figure 10.**
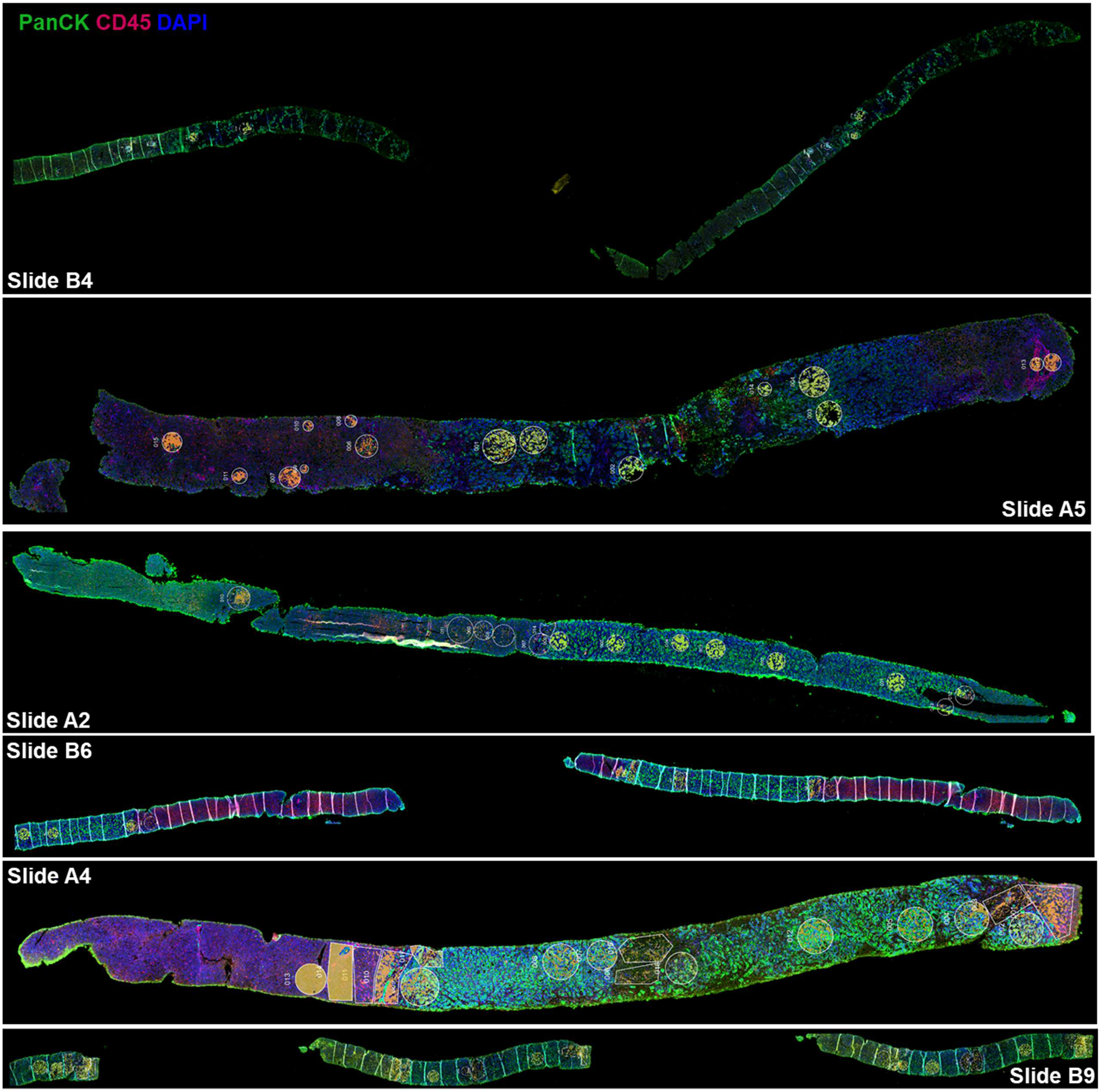
Needle biopsies scanned by the GeoMx Digital Spatial Profiler (DSP). Whole slide scans are provided for all needle biopsies imaged by the GeoMx DSP (n=6). PanCK is indicated by green, CD45 is indicated by red, and DAPI is indicated by blue staining. Slide IDs are provided on each image in white text.

**Supplementary Figure 11.**
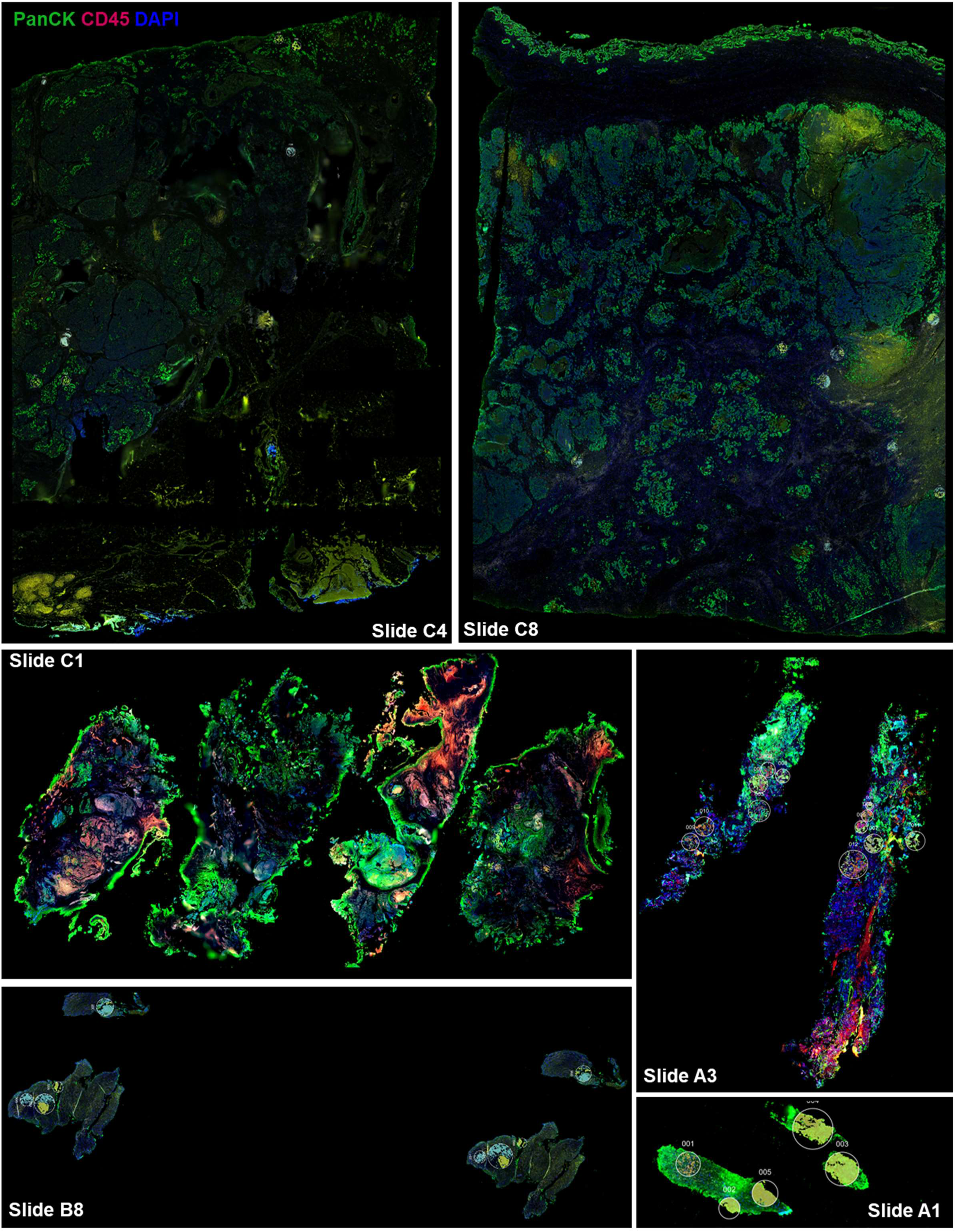
Tissue biopsies scanned by the GeoMx Digital Spatial Profiler (DSP). Whole slide scans are provided for all tissue biopsies imaged by the GeoMx DSP (n=6). PanCK is indicated by green, CD45 is indicated by red, and DAPI is indicated by blue staining. Slide IDs are provided on each image in white text.

**Supplementary Figure 12.**
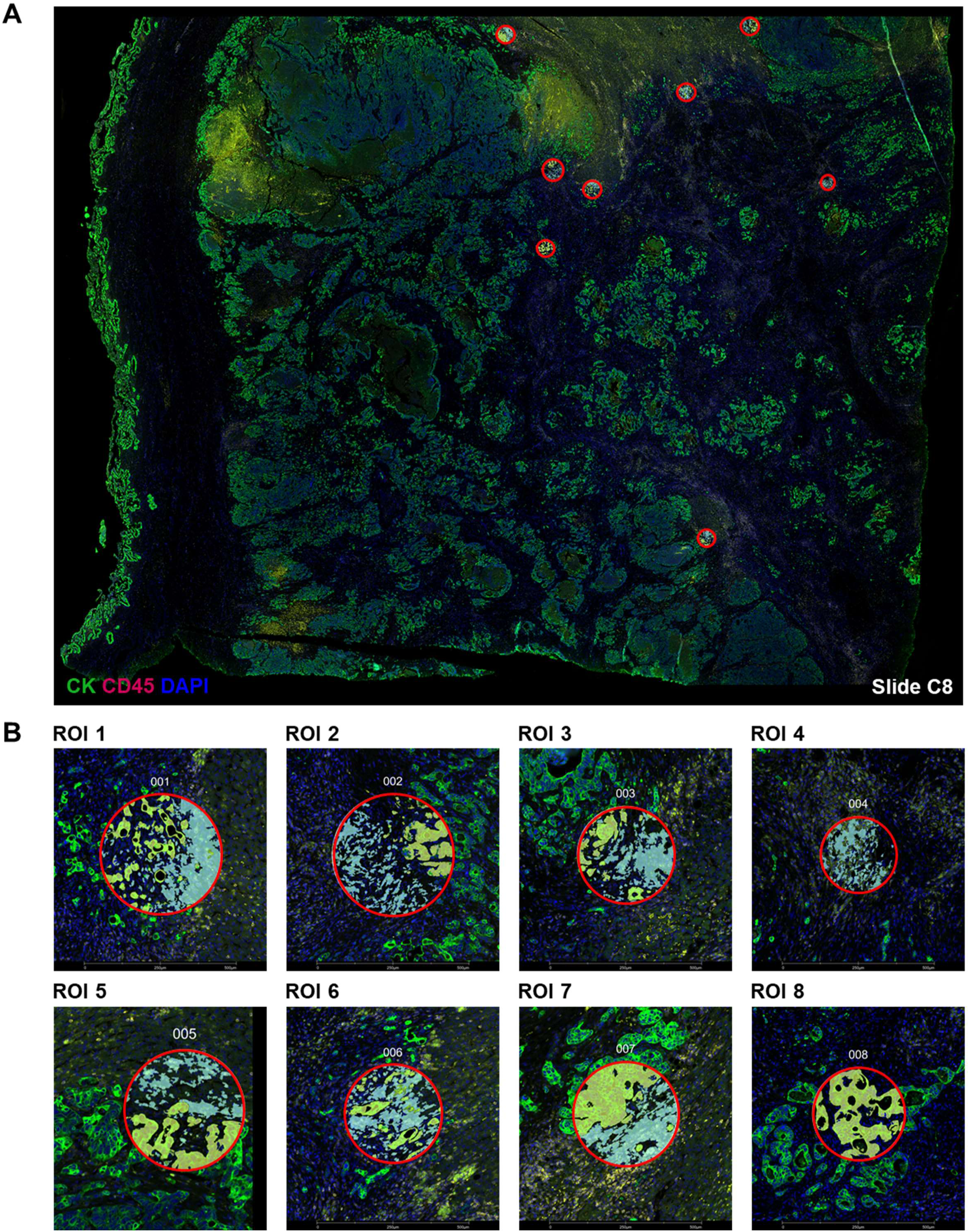
ROI selection and segmentation strategy using PanCK and CD45 markers. **(A)** Representative image of whole tissue scan used for ROI identification. ROIs are outlined in red circles. **(B)** Individual ROIs are ordered by number. Fluorescent channel settings: FITC / 525nm / SYTO 13 / DNA (Blue), Cy3 / 568nm / Alexa 532 / PanCK (Green), and Texas Red / 615nm / Alexa 594 / CD45 (Red). A yellow mask was used for PanCK+ tumor cell identification and a teal mask was used for CD45+ hematopoietic cell identification.

**Supplementary Figure 13.**
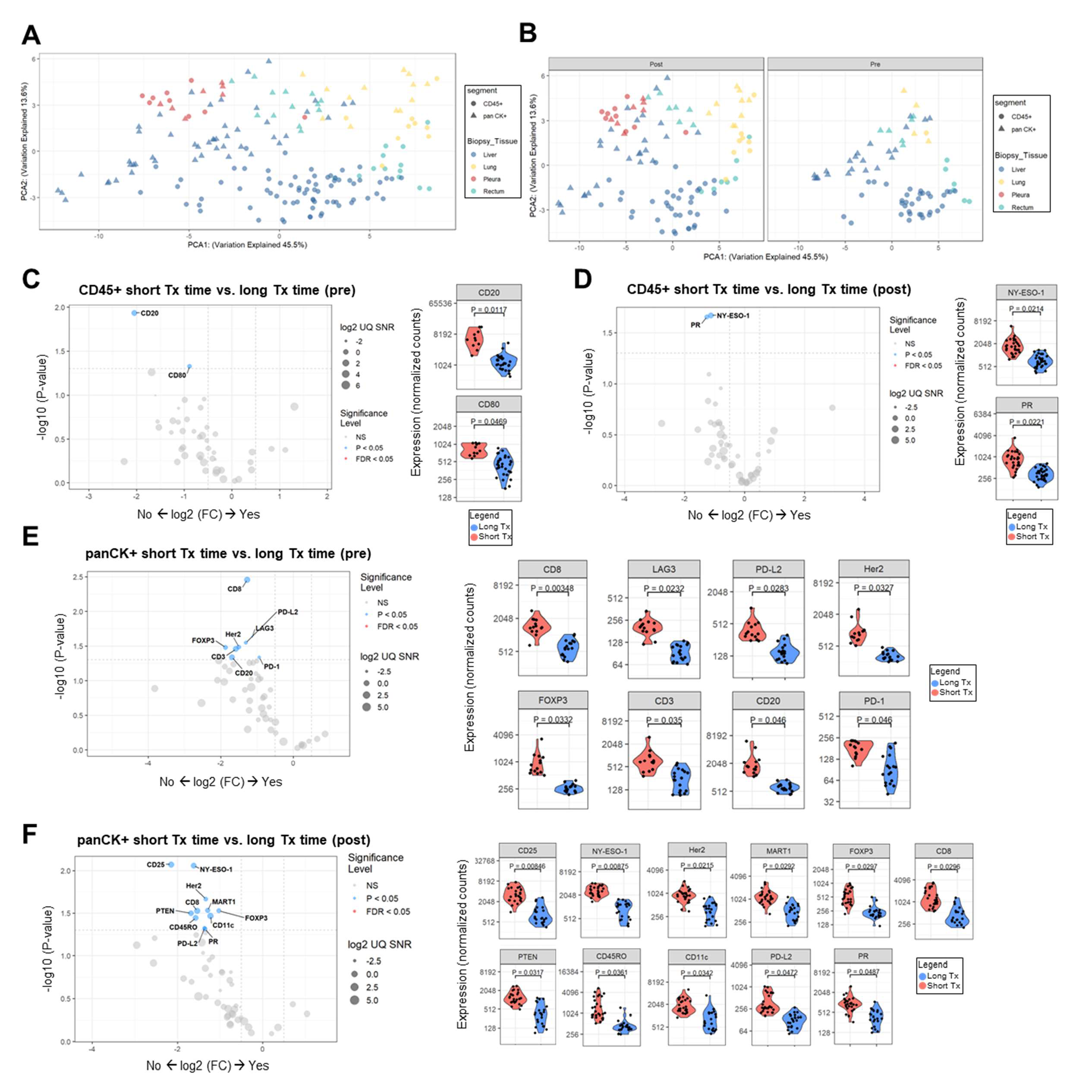
Several proteins are differentially expressed when patients are segmented into two groups based on time-on-treatment. PCA plots showing how similar the different Group levels are from one another in the **(A)** total sample set and **(B)** at both biopsies timepoints (pre-, post-treatment). Samples tend to cluster by tissue type and further separate by segment on PC2. Circles represent CD45+ segments and triangles represent pan CK+ segments. Biopsy tissue locations are color-coded where blue indicates liver, yellow indicates lung, red indicates pleura, and green indicates rectum. **(C)** Volcano plot showing the comparison of pre-treatment biopsy protein expression in CD45+ segments between long time-on-treatment (Long Tx) patients and short time-on-treatment (Short Tx) patients. Violin plots show statistically significant differentially expressed proteins including B cell marker CD20 (p = 0.012) and myeloid activation marker CD80 (p = 0.047). **(D)** Volcano plot showing the comparison of CD45+ segments in post-treatment biopsies between Long Tx patients and Short Tx patients. Violin plots show statistically significant differentially expressed proteins including antigen NY-ESO-1 (p = 0.021) and progesterone receptor (PR) (p = 0.022). **(E)** Volcano plot showing the comparison of pre-treatment protein expression in PanCK+ segments between Long Tx patients and Short Tx patients. Violin plots show statistically significant differentially expressed proteins including cytotoxic T cell marker CD8 (p = 3.5E-3), antigen Her2 (p = 0.033), Treg marker Foxp3 (p = 0.033), T cell marker CD3 (p = 0.035), B cell marker CD20 (p = 0.046), LAG3 (p = 0.023), PD-L2 (p = 0.028), and PD-1 (p = 0.046). **(F)** Volcano plot showing the comparison of post-treatment protein expression in PanCK+ segments between Long Tx patients and Short Tx patients. Violin plots show statistically significant differentially expressed proteins including mature B cell/DC marker CD35 (p = 8.5E-3), antigen NY-ESO-1 (p = 8.7E-3), antigen Her2 (p = 0.022), antigen MART1 (p = 0.029), cytotoxic T cell marker CD8 (p = 0.030), Treg marker Foxp3 (p = 0.030), antigen PTEN (p = 0.032), DC/myeloid marker CD11c (p = 0.034), memory T cell marker CD45RO (p = 0.036), checkpoint PD-L1 (p = 0.047), and PR (p = 0.049).

**Supplementary Figure 14.**
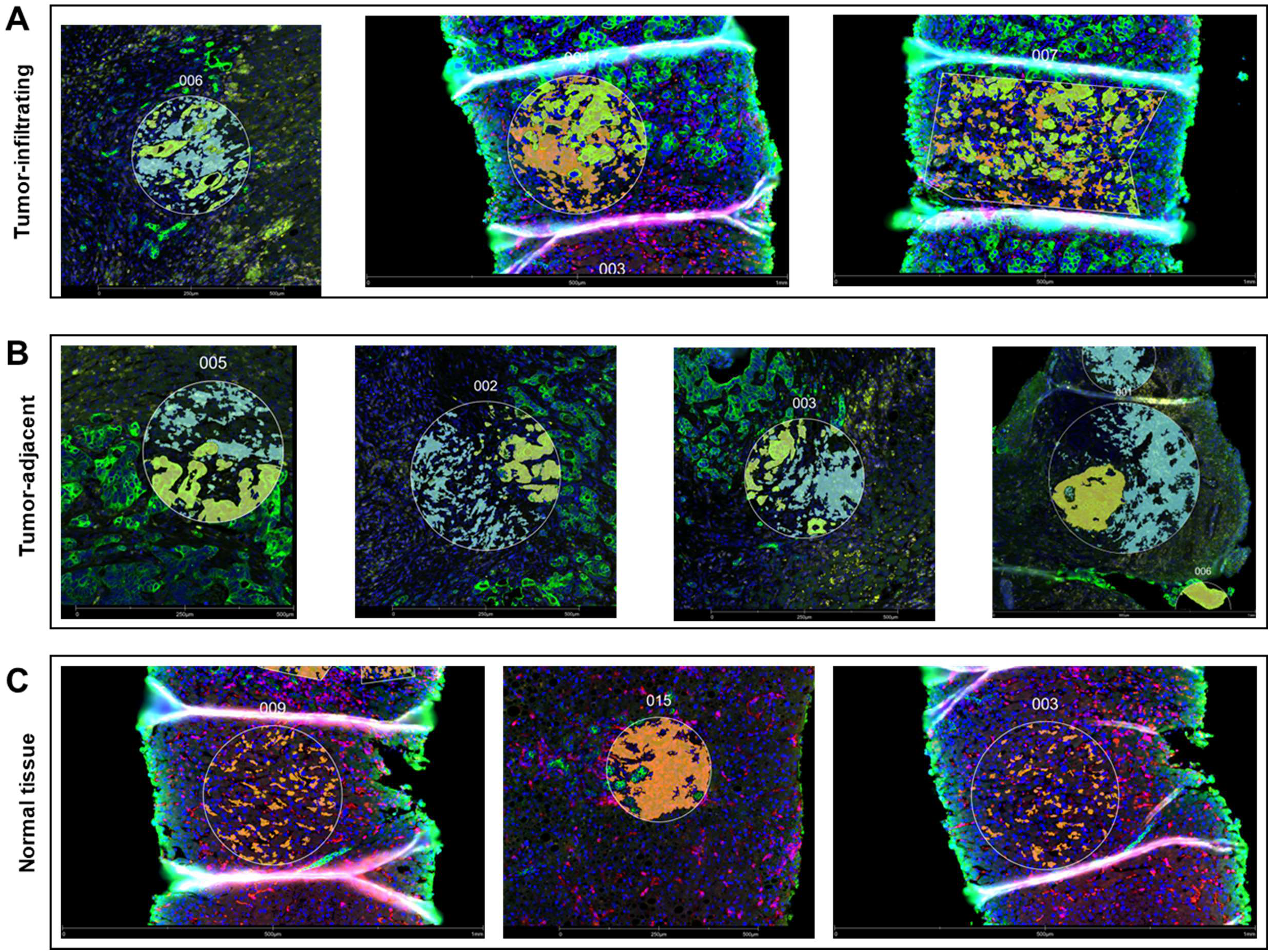
Immune cell localization categorization strategy. Immune cell locations were categorized as **(A)** tumor-infiltrating, **(B)** tumor-adjacent, or **(C)** normal tissue. Representative ROI images are shown.

**Supplementary Figure 15.**
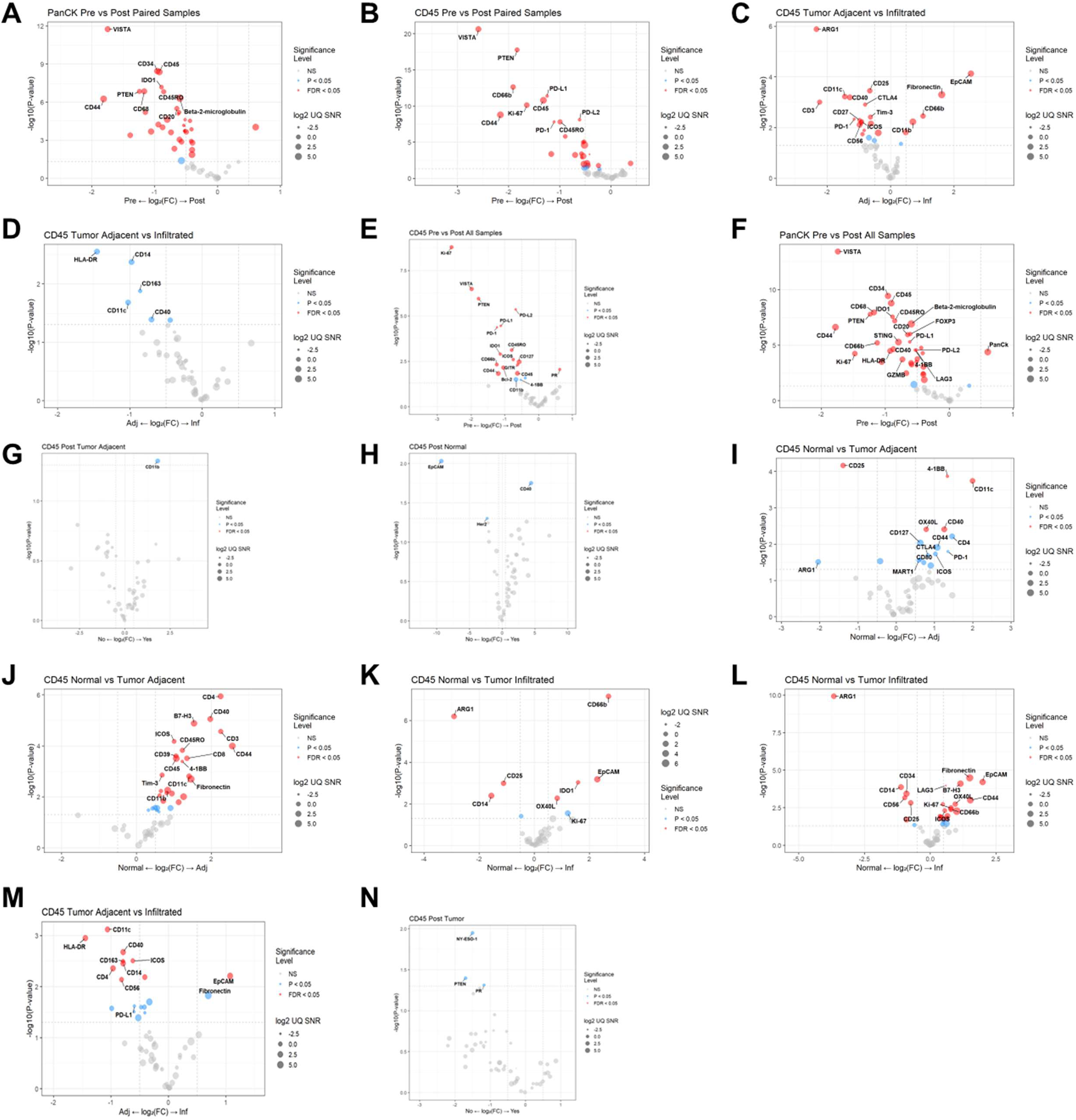
Differential expression analyses of protein expression in human tumor biopsies pre-and post-elraglusib treatment. **(A)** Volcano plot showing differential protein expression in PanCK+ regions between pre-and post-treatment biopsies in paired samples. **(B)** Volcano plot showing differential protein expression in CD45+ regions between pre-and post-treatment biopsies in paired samples. **(C)** Volcano plot showing differential protein expression between tumor-adjacent CD45+ segments and tumor-infiltrating CD45+ segments in pre-treatment biopsies. **(D)** Volcano plot showing differential protein expression between tumor-adjacent CD45+ segments and tumor-infiltrating CD45+ segments in post-treatment biopsies. **(E)** Volcano plot showing differential protein expression between CD45+ regions of pre-treatment and post-treatment biopsies. **(F)** Volcano plot showing differential protein expression panCK+ regions of pre-treatment and post-treatment biopsies. **(G)** Volcano plot showing differential post-treatment protein expression in tumor-adjacent CD45+ immune cell segments in Long Tx patients as compared to Short Tx patients. **(H)** Volcano plot showing differential post-treatment protein expression in CD45+ immune cell segments located in normal (non-tumor) tissue in Long Tx patients as compared to Short Tx patients. **(I)** Volcano plot showing differential pre-treatment protein expression in CD45+ immune cell segments located in normal (non-tumor) tissue as compared to tumor-adjacent tissue. **(J)** Volcano plot showing differential post-treatment protein expression in CD45+ immune cell segments located in normal (non-tumor) tissue as compared to tumor-adjacent tissue. **(K)** Volcano plot showing differential pre-treatment protein expression in CD45+ immune cell segments located in normal (non-tumor) tissue as compared to those in tumor tissue. **(L)** Volcano plot showing differential post-treatment protein expression in CD45+ immune cell segments located in normal (non-tumor) tissue as compared to those in tumor tissue. **(M)** Volcano plot showing a comparison of tumor-adjacent CD45+ segments with tumor-infiltrating CD45+ segments in all biopsies regardless of timepoint. **(N)** Volcano plot showing a comparison of post-treatment protein expression of tumor-infiltrating CD45+ immune cell segments in Long Tx patients as compared to Short Tx patients. Grey points are non-significant (NS), blue points have p values < 0.05, and red points have false discovery rate (FDR) values less than 0.05. The size of the point represents the log2 UQ Signal-to-noise ratio (SNR).

**Supplementary Table 1.**
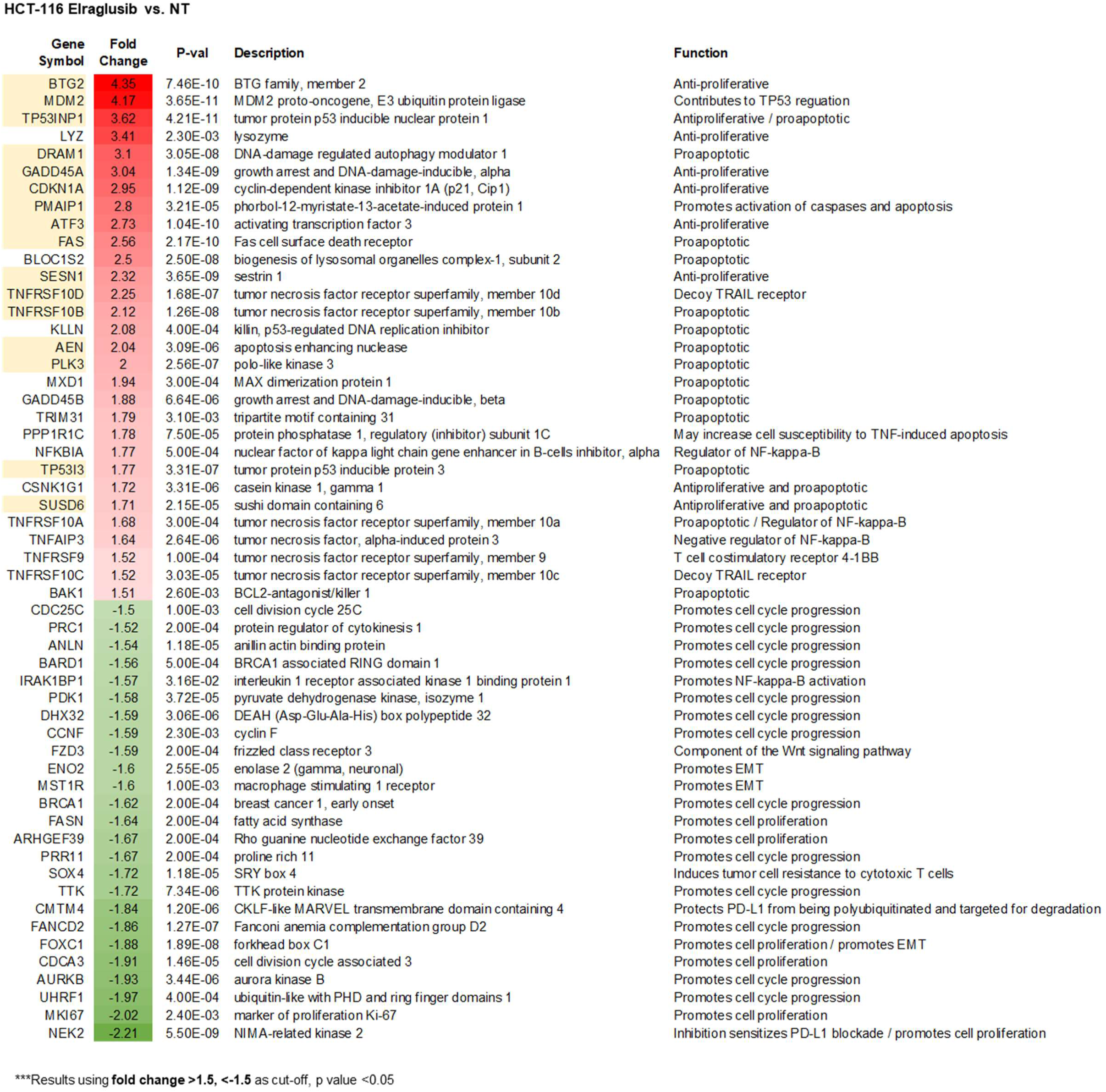
HCT-116 CRC cell microarray analysis. HCT-116 CRC cells were treated with 1 μM elraglusib for 24 hours and treated versus untreated control samples were compared in triplicate via microarray analysis. Table showing differentially expressed genes of interest with their corresponding FCs, p values, descriptions, and functions. Genes highlighted in yellow are known p53 targets (13). Genes are ordered by FC and were calculated using a FC cutoff of >1.5, <-1.5, and a minimum p value of <0.05.

**Supplementary Table 2.**
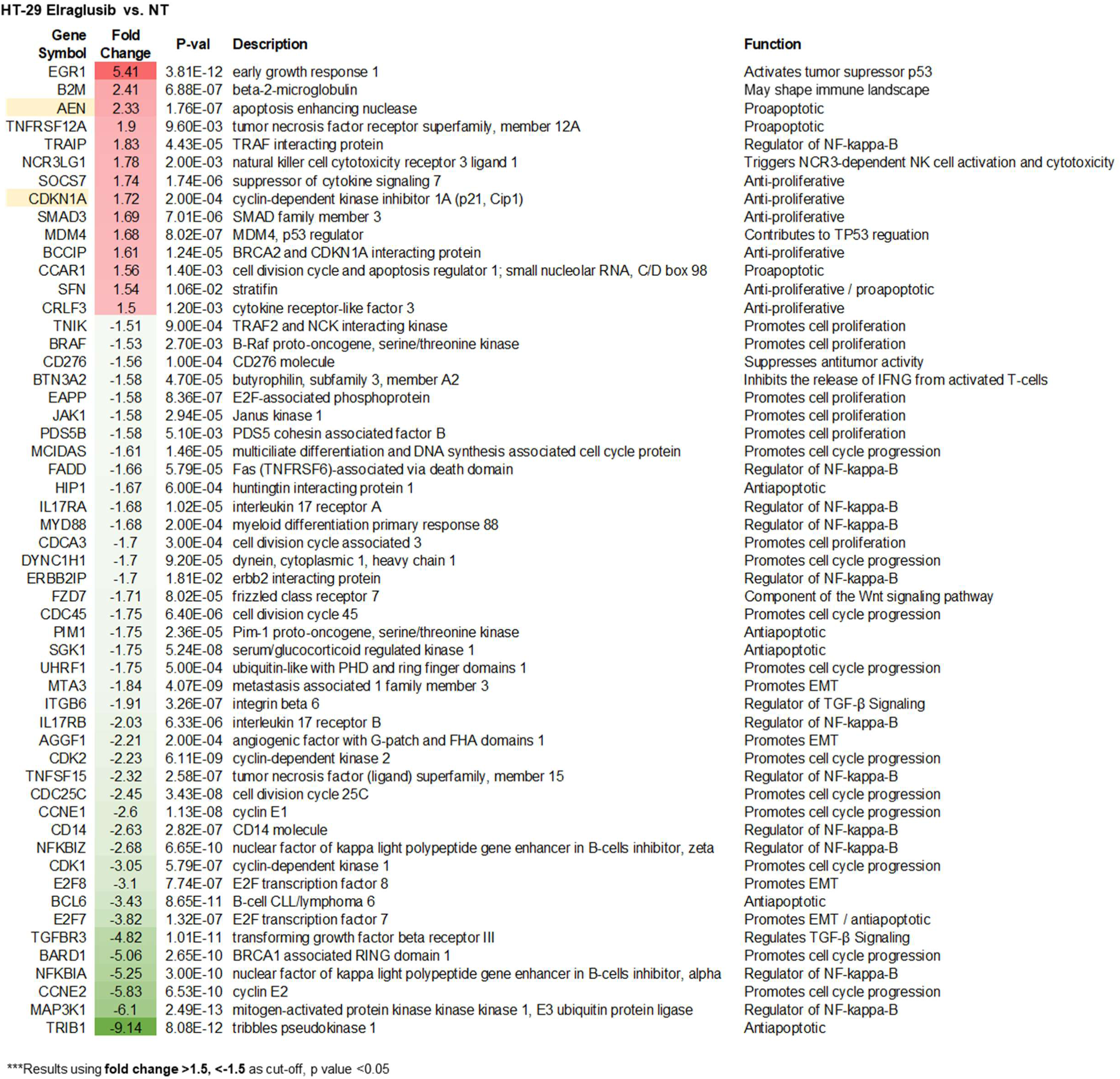
HT-29 CRC cell microarray analysis. HT-29 CRC cells were treated with 1 μM elraglusib for 24 hours and treated versus untreated control samples were compared in triplicate via microarray analysis. Table showing differentially expressed genes of interest with their corresponding FCs, p values, descriptions, and functions. Genes highlighted in yellow are known p53 targets. Genes are ordered by FC and were calculated using a FC cutoff of >1.5, <- 1.5, and a minimum p value of <0.05.

**Supplementary Table 3.**
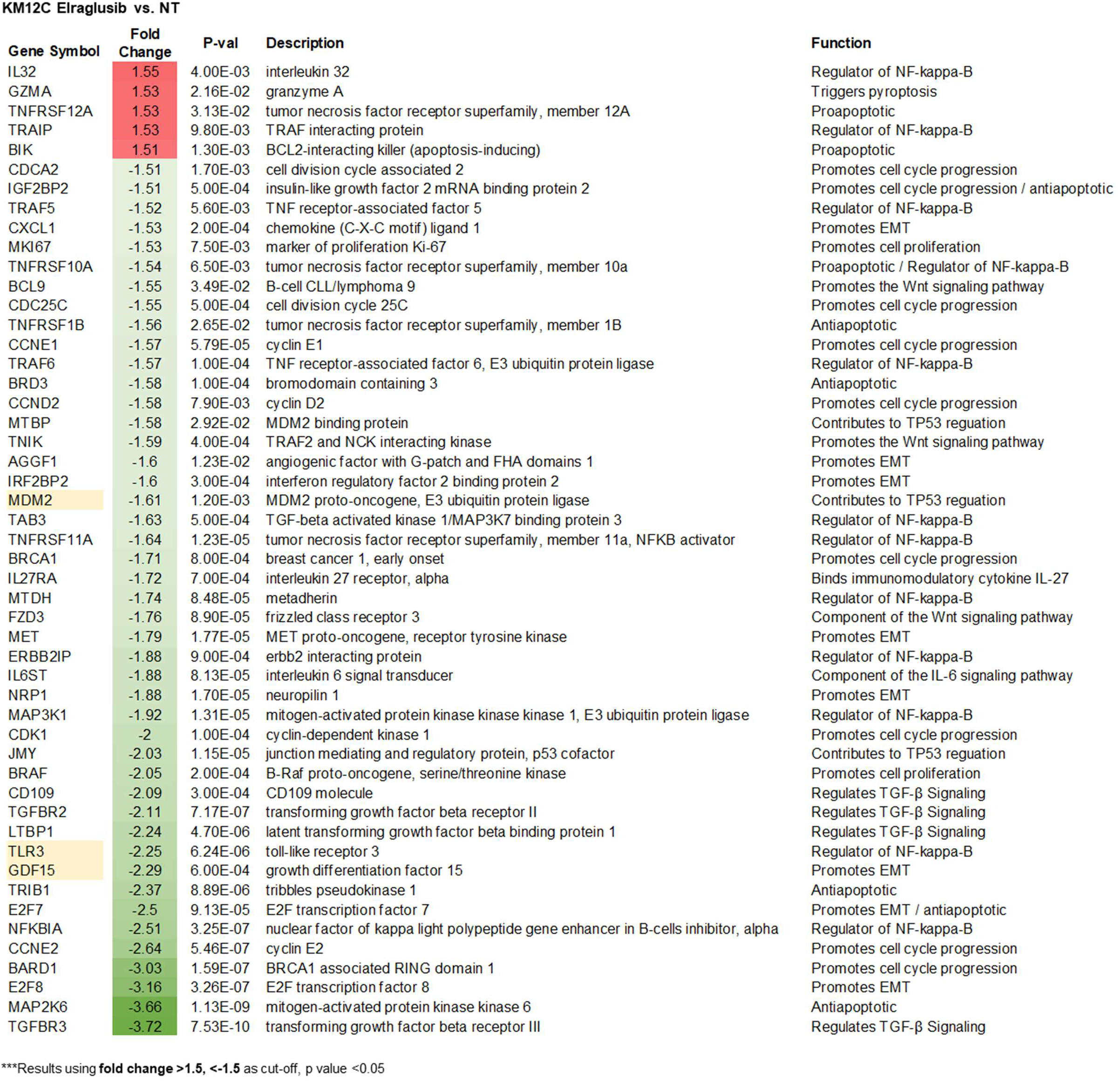
KM12C CRC cell microarray analysis. KM12C CRC cells were treated with 1 μM elraglusib for 24 hours and treated versus untreated control samples were compared in triplicate via microarray analysis. Table showing differentially expressed genes of interest with their corresponding FCs, p values, descriptions, and functions. Genes highlighted in yellow are known p53 targets. Genes are ordered by FC and were calculated using a FC cutoff of >1.5, <-1.5, and a minimum p value of <0.05.

**Supplementary Table 4.**
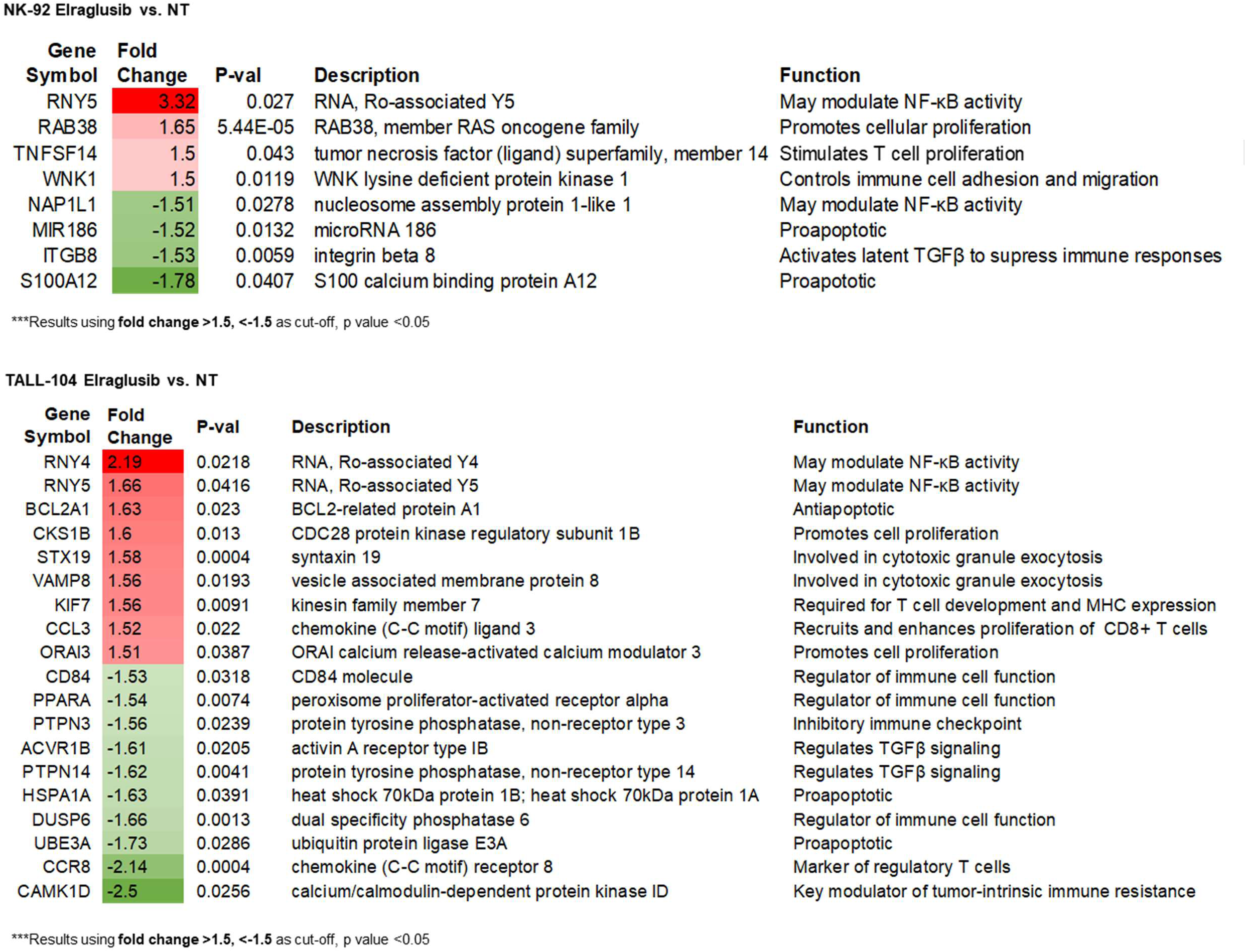
Immune cell microarray analysis. NK-92 and TALL-104 immune cells were treated with 1 μM elraglusib for 24 hours and treated versus untreated control samples were compared in triplicate via microarray analysis. Tables showing differentially expressed genes of interest with their corresponding FCs, p values, descriptions, and functions. Genes are ordered by FC and were calculated using a FC cutoff of >1.5, <-1.5, and a minimum p value of <0.05.

NK-92 Elraglusib vs. NT
| Gene Symbol | Fold Change | P-val | Description | Function |
| --- | --- | --- | --- | --- |
| RNY5 | 3.32 | 0.027 | RNA, Ro-associated Y5 | May modulate NF-κB activity |
| RAB38 | 1.65 | 5.44E-05 | RAB38, member RAS oncogene family | Promotes cellular proliferation |
| TNFSF14 | 1.5 | 0.043 | tumor necrosis factor (ligand) superfamily, member 14 | Stimulates T cell proliferation |
| WNK1 | 1.5 | 0.0119 | WNK lysine deficient protein kinase 1 | Controls immune cell adhesion and migration |
| NAP1L1 | -1.51 | 0.0278 | nucleosome assembly protein 1-like 1 | May modulate NF-κB activity |
| MIR186 | -1.52 | 0.0132 | microRNA 186 | Proapoptotic |
| ITGB8 | -1.53 | 0.0059 | integrin beta 8 | Activates latent TGFβ to suppress immune responses |
| S100A12 | -1.78 | 0.0407 | S100 calcium binding protein A12 | Proapoptotic |

\*\*\*Results using fold change >1.5, <-1.5 as cut-off, p value <0.05

TALL-104 Elraglusib vs. NT
| Gene Symbol | Fold Change | P-val | Description | Function |
| --- | --- | --- | --- | --- |
| RNY4 | 2.19 | 0.0218 | RNA, Ro-associated Y4 | May modulate NF-κB activity |
| RNY5 | 1.66 | 0.0416 | RNA, Ro-associated Y5 | May modulate NF-κB activity |
| BCL2A1 | 1.63 | 0.023 | BCL2-related protein A1 | Antiapoptotic |
| CKS1B | 1.6 | 0.013 | CDC28 protein kinase regulatory subunit 1B | Promotes cell proliferation |
| STX19 | 1.58 | 0.0004 | syntaxin 19 | Involved in cytotoxic granule exocytosis |
| VAMP8 | 1.56 | 0.0193 | vesicle associated membrane protein 8 | Involved in cytotoxic granule exocytosis |
| KIF7 | 1.56 | 0.0091 | kinesin family member 7 | Required for T cell development and MHC expression |
| CCL3 | 1.52 | 0.022 | chemokine (C-C motif) ligand 3 | Recruits and enhances proliferation of CD8+ T cells |
| ORAI3 | 1.51 | 0.0387 | ORAI calcium release-activated calcium modulator 3 | Promotes cell proliferation |
| CD84 | -1.53 | 0.0318 | CD84 molecule | Regulator of immune cell function |
| PPARA | -1.54 | 0.0074 | peroxisome proliferator-activated receptor alpha | Regulator of immune cell function |
| PTPN3 | -1.56 | 0.0239 | protein tyrosine phosphatase, non-receptor type 3 | Inhibitory immune checkpoint |
| ACVR1B | -1.61 | 0.0205 | activin A receptor type 1B | Regulates TGFβ signaling |
| PTPN14 | -1.62 | 0.0041 | protein tyrosine phosphatase, non-receptor type 14 | Regulates TGFβ signaling |
| HSPA1A | -1.63 | 0.0391 | heat shock 70kDa protein 1B; heat shock 70kDa protein 1A | Proapoptotic |
| DUSP6 | -1.66 | 0.0013 | dual specificity phosphatase 6 | Regulator of immune cell function |
| UBE3A | -1.73 | 0.0286 | ubiquitin protein ligase E3A | Proapoptotic |
| CCR8 | -2.14 | 0.0004 | chemokine (C-C motif) receptor 8 | Marker of regulatory T cells |
| CAMK1D | -2.5 | 0.0256 | calcium/calmodulin-dependent protein kinase ID | Key modulator of tumor-intrinsic immune resistance |

\*\*\*Results using fold change >1.5, <-1.5 as cut-off, p value <0.05

**Supplementary Table 5.**
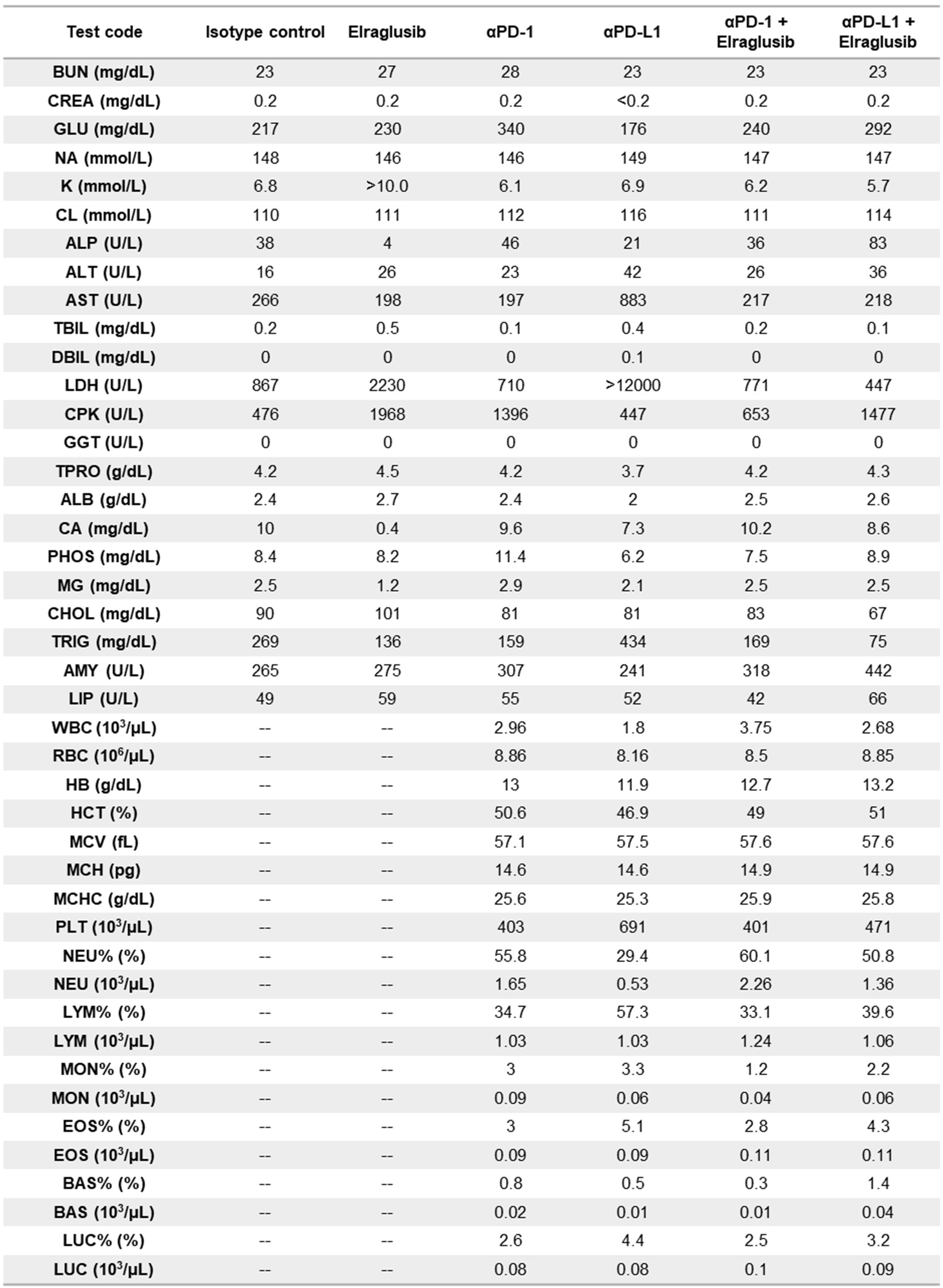
Murine serum chemistry analysis does not show treatment-related toxicity. Whole blood from long-term mice sacrificed was submitted for serum chemistry analysis. Results are shown from the complete metabolic panel and complete blood count with differential. ALB: Albumin, ALP: Alkaline Phosphatase, ALT: Alanine aminotransferase, AMY: Amylase, ANIS: Anisocytosis, AST: Aspartate aminotransferase, ATYP: Atypical Lymphs, BAS: Absolute Basophils, BAS%: % Basophils, BUN: Urea Nitrogen, CA: Calcium, CHOL: Cholesterol, CL: Chloride, CPK: Creatine kinase, CPLT: Clumped Platelets, CREA: Creatinine, DBIL: Direct Bilirubin, EOS: Absolute Eosinophils, EOS%: % Eosinophils, GGT: Gamma-glutamyl Transferase, GLU: Glucose, HB: Hemoglobin, HCT: Hematocrit, HJB: Howell-Jolly Bodies, HYPO: Hypochromasia, HYPR: Hyperchromasia, K: Potassium, LDH: Lactate Dehydrogenase, LIP: Lipase, LPLT: Large Platelets, LUC: Absolute Large Unstained Cells, LUC%: % Large Unstained Cells, LYM: Absolute Lymphocytes, LYM%: % Lymphocytes, MAC: Macrocytosis, MCH: Mean Corpuscular Hemoglobin, MCHC: Mean Corpuscular Hemoglobin Count, MCV: Mean Corpuscular Volume, MG: Magnesium, MIC: Microcytosis, MON: Absolute Monocytes, MON%: % Monocytes, NA: Sodium, NEU: Absolute Neutrophils, NEU%: % Neutrophils, PHOS: Inorganic Phosphorus, PLT: Platelet Count, POLK: Poikilocytosis, RBC: Red Blood Cell Count, TBIL: Total Bilirubin, TPRO: Total Protein, TRIG: Triglyceride, WBC: White Blood Cell Count.

| Test code | Isotype control | Elraglusib | αPD-1 | αPD-L1 | αPD-1 + Elraglusib | αPD-L1 + Elraglusib |
| --- | --- | --- | --- | --- | --- | --- |
| BUN (mg/dL) | 23 | 27 | 28 | 23 | 23 | 23 |
| CREA (mg/dL) | 0.2 | 0.2 | 0.2 | <0.2 | 0.2 | 0.2 |
| GLU (mg/dL) | 217 | 230 | 340 | 176 | 240 | 292 |
| NA (mmol/L) | 148 | 146 | 146 | 149 | 147 | 147 |
| K (mmol/L) | 6.8 | >10.0 | 6.1 | 6.9 | 6.2 | 5.7 |
| CL (mmol/L) | 110 | 111 | 112 | 116 | 111 | 114 |
| ALP (U/L) | 38 | 4 | 46 | 21 | 36 | 83 |
| ALT (U/L) | 16 | 26 | 23 | 42 | 26 | 36 |
| AST (U/L) | 266 | 198 | 197 | 883 | 217 | 218 |
| TBIL (mg/dL) | 0.2 | 0.5 | 0.1 | 0.4 | 0.2 | 0.1 |
| DBIL (mg/dL) | 0 | 0 | 0 | 0.1 | 0 | 0 |
| LDH (U/L) | 867 | 2230 | 710 | >12000 | 771 | 447 |
| CPK (U/L) | 476 | 1968 | 1396 | 447 | 653 | 1477 |
| GGT (U/L) | 0 | 0 | 0 | 0 | 0 | 0 |
| TPRO (g/dL) | 4.2 | 4.5 | 4.2 | 3.7 | 4.2 | 4.3 |
| ALB (g/dL) | 2.4 | 2.7 | 2.4 | 2 | 2.5 | 2.6 |
| CA (mg/dL) | 10 | 0.4 | 9.6 | 7.3 | 10.2 | 8.6 |
| PHOS (mg/dL) | 8.4 | 8.2 | 11.4 | 6.2 | 7.5 | 8.9 |
| MG (mg/dL) | 2.5 | 1.2 | 2.9 | 2.1 | 2.5 | 2.5 |
| CHOL (mg/dL) | 90 | 101 | 81 | 81 | 83 | 67 |
| TRIG (mg/dL) | 269 | 136 | 159 | 434 | 169 | 75 |
| AMY (U/L) | 265 | 275 | 307 | 241 | 318 | 442 |
| LIP (U/L) | 49 | 59 | 55 | 52 | 42 | 66 |
| WBC (10<sup>3</sup>/μL) | -- | -- | 2.96 | 1.8 | 3.75 | 2.68 |
| RBC (10<sup>6</sup>/μL) | -- | -- | 8.86 | 8.16 | 8.5 | 8.85 |
| HB (g/dL) | -- | -- | 13 | 11.9 | 12.7 | 13.2 |
| HCT (%) | -- | -- | 50.6 | 46.9 | 49 | 51 |
| MCV (fL) | -- | -- | 57.1 | 57.5 | 57.6 | 57.6 |
| MCH (pg) | -- | -- | 14.6 | 14.6 | 14.9 | 14.9 |
| MCHC (g/dL) | -- | -- | 25.6 | 25.3 | 25.9 | 25.8 |
| PLT (10<sup>3</sup>/μL) | -- | -- | 403 | 691 | 401 | 471 |
| NEU% (%) | -- | -- | 55.8 | 29.4 | 60.1 | 50.8 |
| NEU (10<sup>3</sup>/μL) | -- | -- | 1.65 | 0.53 | 2.26 | 1.36 |
| LYM% (%) | -- | -- | 34.7 | 57.3 | 33.1 | 39.6 |
| LYM (10<sup>3</sup>/μL) | -- | -- | 1.03 | 1.03 | 1.24 | 1.06 |
| MON% (%) | -- | -- | 3 | 3.3 | 1.2 | 2.2 |
| MON (10<sup>3</sup>/μL) | -- | -- | 0.09 | 0.06 | 0.04 | 0.06 |
| EOS% (%) | -- | -- | 3 | 5.1 | 2.8 | 4.3 |
| EOS (10<sup>3</sup>/μL) | -- | -- | 0.09 | 0.09 | 0.11 | 0.11 |
| BAS% (%) | -- | -- | 0.8 | 0.5 | 0.3 | 1.4 |
| BAS (10<sup>3</sup>/μL) | -- | -- | 0.02 | 0.01 | 0.01 | 0.04 |
| LUC% (%) | -- | -- | 2.6 | 4.4 | 2.5 | 3.2 |
| LUC (10<sup>3</sup>/μL) | -- | -- | 0.08 | 0.08 | 0.1 | 0.09 |

**Supplementary Table 6.**
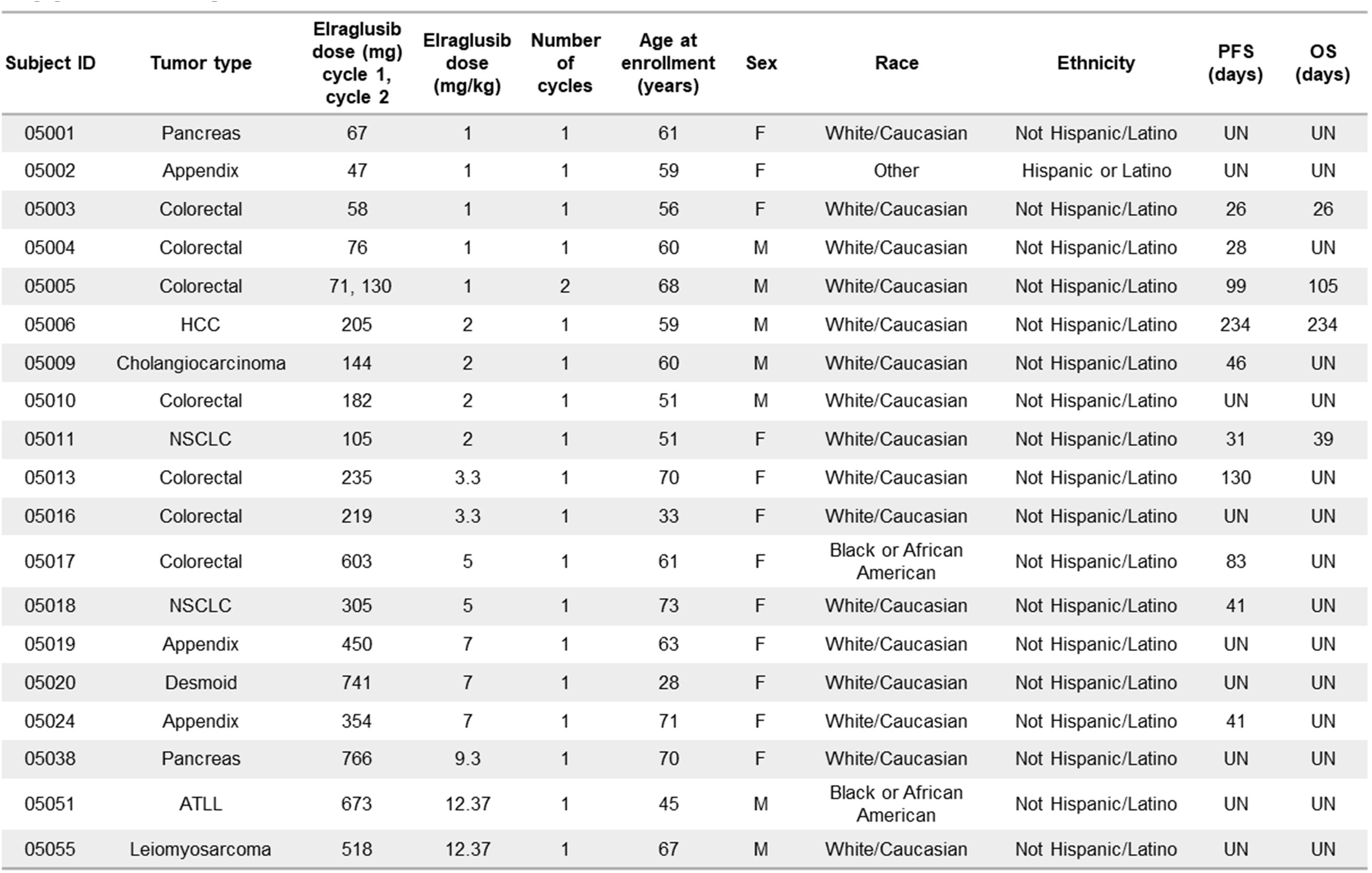
Individual patient information for human patient cytokine analysis. Tumor type, elraglusib dose (mg, mg/kg), number of cycles, age at enrollment, sex, race, ethnicity, median progression-free survival and median overall survival data is shown.

## References

1. Ding L, Madamsetty VS, Kiers S, Alekhina O, Ugolkov A, Dube J, et al. Glycogen Synthase Kinase-3 Inhibition Sensitizes Pancreatic Cancer Cells to Chemotherapy by Abrogating the TopBP1/ATR-Mediated DNA Damage Response. Clin Cancer Res 2019;25(21):6452–62 doi 10.1158/1078-0432.Ccr-19-0799.

2. Augello G, Emma MR, Cusimano A, Azzolina A, Montalto G, McCubrey JA, et al. The Role of GSK-3 in Cancer Immunotherapy: GSK-3 Inhibitors as a New Frontier in Cancer Treatment. Cells 2020;9(6) doi 10.3390/cells9061427.

3. Tsai CC, Tsai CK, Tseng PC, Lin CF, Chen CL. Glycogen Synthase Kinase-3β Facilitates Cytokine Production in 12-O-Tetradecanoylphorbol-13-Acetate/Ionomycin-Activated Human CD4(+) T Lymphocytes. Cells 2020;9(6) doi 10.3390/cells9061424.

4. Taylor A, Harker JA, Chanthong K, Stevenson PG, Zuniga EI, Rudd CE. Glycogen Synthase Kinase 3 Inactivation Drives T-bet-Mediated Downregulation of Co-receptor PD-1 to Enhance CD8(+) Cytolytic T Cell Responses. Immunity 2016;44(2):274–86 doi 10.1016/j.immuni.2016.01.018.

5. Mancinelli R, Carpino G, Petrungaro S, Mammola CL, Tomaipitinca L, Filippini A, et al. Multifaceted Roles of GSK-3 in Cancer and Autophagy-Related Diseases. Oxid Med Cell Longev 2017;2017:4629495 doi 10.1155/2017/4629495.

6. Borelli B, Antoniotti C, Carullo M, Germani MM, Conca V, Masi G. Immune-Checkpoint Inhibitors (ICIs) in Metastatic Colorectal Cancer (mCRC) Patients beyond Microsatellite Instability. Cancers (Basel) 2022;14(20) doi 10.3390/cancers14204974.

7. Gupta R, Sinha S, Paul RN. The impact of microsatellite stability status in colorectal cancer. Current Problems in Cancer 2018;42(6):548–59 doi https://doi.org/10.1016/j.currproblcancer.2018.06.010.

8. Middha S, Yaeger R, Shia J, Stadler ZK, King S, Guercio S, et al. Majority of B2M-Mutant and - Deficient Colorectal Carcinomas Achieve Clinical Benefit From Immune Checkpoint Inhibitor Therapy and Are Microsatellite Instability-High. JCO Precis Oncol 2019;3 doi 10.1200/po.18.00321.

9. Carneiro BA, Cavalcante L, Bastos BR, Powell SF, Ma WW, Sahebjam S, et al. Phase I study of 9-ing-41, a small molecule selective glycogen synthase kinase-3 beta (GSK-3β) inhibitor, as a single agent and combined with chemotherapy, in patients with refractory tumors. Journal of Clinical Oncology 2020;38(15_suppl):3507- doi 10.1200/JCO.2020.38.15_suppl.3507.

10. Rudd CE, Chanthong K, Taylor A. Small Molecule Inhibition of GSK-3 Specifically Inhibits the Transcription of Inhibitory Co-receptor LAG-3 for Enhanced Anti-tumor Immunity. Cell Rep 2020;30(7):2075–82.e4 doi 10.1016/j.celrep.2020.01.076.

11. Tang YY, Sheng SY, Lu CG, Zhang YQ, Zou JY, Lei YY, et al. Effects of Glycogen Synthase Kinase-3β Inhibitor TWS119 on Proliferation and Cytokine Production of TILs From Human Lung Cancer. J Immunother 2018;41(7):319–28 doi 10.1097/cji.0000000000000234.

12. Taylor A, Rudd CE. Glycogen synthase kinase 3 (GSK-3) controls T-cell motility and interactions with antigen-presenting cells. BMC Res Notes 2020;13(1):163- doi 10.1186/s13104-020-04971-0.

13. Fischer M. Census and evaluation of p53 target genes. Oncogene 2017;36(28):3943–56 doi 10.1038/onc.2016.502.

14. Zeng Q, Huang Y, Zeng L, Lan X, Huang Y, He S, et al. Effect of IPP5, a novel inhibitor of PP1, on apoptosis and the underlying mechanisms involved. Biotechnol Appl Biochem 2009;54(4):231–8 doi 10.1042/ba20090168.

15. Mezzadra R, Sun C, Jae LT, Gomez-Eerland R, de Vries E, Wu W, et al. Identification of CMTM6 and CMTM4 as PD-L1 protein regulators. Nature 2017;549(7670):106–10 doi 10.1038/nature23669.

16. Zhang X, Huang X, Xu J, Li E, Lao M, Tang T, et al. NEK2 inhibition triggers anti-pancreatic cancer immunity by targeting PD-L1. Nature Communications 2021;12(1):4536 doi 10.1038/s41467-021-24769-3.

17. Brandt CS, Baratin M, Yi EC, Kennedy J, Gao Z, Fox B, et al. The B7 family member B7-H6 is a tumor cell ligand for the activating natural killer cell receptor NKp30 in humans. Journal of Experimental Medicine 2009;206(7):1495–503 doi 10.1084/jem.20090681.

18. Zhou Z, He H, Wang K, Shi X, Wang Y, Su Y, et al. Granzyme A from cytotoxic lymphocytes cleaves GSDMB to trigger pyroptosis in target cells. Science 2020;368(6494) doi 10.1126/science.aaz7548.

19. Huntington KE, Louie A, Zhou L, El-Deiry WS. A high-throughput customized cytokinome screen of colon cancer cell responses to small-molecule oncology drugs. Oncotarget; Vol 12, No 20 2021.

20. Kaler P, Augenlicht L, Klampfer L. Activating Mutations in β-Catenin in Colon Cancer Cells Alter Their Interaction with Macrophages; the Role of Snail. PLOS ONE 2012;7(9):e45462 doi 10.1371/journal.pone.0045462.

21. Rowan AJ, Lamlum H, Ilyas M, Wheeler J, Straub J, Papadopoulou A, et al. APC mutations in sporadic colorectal tumors: A mutational “hotspot” and interdependence of the “two hits”. Proceedings of the National Academy of Sciences 2000;97(7):3352 doi 10.1073/pnas.97.7.3352.

22. Honey K. CCL3 and CCL4 actively recruit CD8+ T cells. Nature Reviews Immunology 2006;6(6):427- doi 10.1038/nri1862.

23. Chou W-C, Rampanelli E, Li X, Ting JPY. Impact of intracellular innate immune receptors on immunometabolism. Cellular & Molecular Immunology 2022;19(3):337–51 doi 10.1038/s41423-021-00780-y.

24. Sideras K, Galjart B, Vasaturo A, Pedroza-Gonzalez A, Biermann K, Mancham S, et al. Prognostic value of intra-tumoral CD8+/FoxP3+ lymphocyte ratio in patients with resected colorectal cancer liver metastasis. Journal of Surgical Oncology 2018;118(1):68–76 doi https://doi.org/10.1002/jso.25091.

25. Borsellino G, Kleinewietfeld M, Di Mitri D, Sternjak A, Diamantini A, Giometto R, et al. Expression of ectonucleotidase CD39 by Foxp3+ Treg cells: hydrolysis of extracellular ATP and immune suppression. Blood 2007;110(4):1225–32 doi 10.1182/blood-2006-12-064527.

26. Gavalas NG, Tsiatas M, Tsitsilonis O, Politi E, Ioannou K, Ziogas AC, et al. VEGF directly suppresses activation of T cells from ascites secondary to ovarian cancer via VEGF receptor type 2. British Journal of Cancer 2012;107(11):1869–75 doi 10.1038/bjc.2012.468.

27. Starnes T, Rasila KK, Robertson MJ, Brahmi Z, Dahl R, Christopherson K, et al. The chemokine CXCL14 (BRAK) stimulates activated NK cell migration: Implications for the downregulation of CXCL14 in malignancy. Experimental Hematology 2006;34(8):1101–5 doi https://doi.org/10.1016/j.exphem.2006.05.015.

28. Hassounah NB, Malladi VS, Huang Y, Freeman SS, Beauchamp EM, Koyama S, et al. Identification and characterization of an alternative cancer-derived PD-L1 splice variant. Cancer Immunol Immunother 2019;68(3):407–20 doi 10.1007/s00262-018-2284-z.

29. Lu X, Guo T, Zhang X. Pyroptosis in Cancer: Friend or Foe? Cancers. Volume 132021.

30. Medunjanin S, Schleithoff L, Fiegehenn C, Weinert S, Zuschratter W, Braun-Dullaeus RC. GSK-3β controls NF-kappaB activity via IKKγ/NEMO. Sci Rep 2016;6:38553 doi 10.1038/srep38553.

31. Thu YM, Richmond A. NF-κB inducing kinase: a key regulator in the immune system and in cancer. Cytokine Growth Factor Rev 2010;21(4):213–26 doi 10.1016/j.cytogfr.2010.06.002.

32. Park R, Coveler AL, Cavalcante L, Saeed A. GSK-3β in Pancreatic Cancer: Spotlight on 9-ING-41, Its Therapeutic Potential and Immune Modulatory Properties. Biology (Basel) 2021;10(7) doi 10.3390/biology10070610.

33. Tay RE, Richardson EK, Toh HC. Revisiting the role of CD4+ T cells in cancer immunotherapy—new insights into old paradigms. Cancer Gene Therapy 2021;28(1):5–17 doi 10.1038/s41417-020-0183-x.

34. Rihacek M, Bienertova-Vasku J, Valik D, Sterba J, Pilatova K, Zdrazilova-Dubska L. B-Cell Activating Factor as a Cancer Biomarker and Its Implications in Cancer-Related Cachexia. Biomed Res Int 2015;2015:792187- doi 10.1155/2015/792187.

35. Lee YS, Kim SY, Song SJ, Hong HK, Lee Y, Oh BY, et al. Crosstalk between CCL7 and CCR3 promotes metastasis of colon cancer cells via ERK-JNK signaling pathways. Oncotarget 2016;7(24):36842–53 doi 10.18632/oncotarget.9209.

36. Yu X, Wang D, Wang X, Sun S, Zhang Y, Wang S, et al. CXCL12/CXCR4 promotes inflammation-driven colorectal cancer progression through activation of RhoA signaling by sponging miR-133a-3p. Journal of Experimental & Clinical Cancer Research 2019;38(1):32 doi 10.1186/s13046-018-1014-x.

37. Bendardaf R, Buhmeida A, Hilska M, Laato M, Syrjänen S, Syrjänen K, et al. VEGF-1 expression in colorectal cancer is associated with disease localization, stage, and long-term disease-specific survival. Anticancer Res 2008;28(6b):3865–70.

38. Zhong M, Li N, Qiu X, Ye Y, Chen H, Hua J, et al. TIPE regulates VEGFR2 expression and promotes angiogenesis in colorectal cancer. Int J Biol Sci 2020;16(2):272–83 doi 10.7150/ijbs.37906.

39. Li J, Sun R, Tao K, Wang G. The CCL21/CCR7 pathway plays a key role in human colon cancer metastasis through regulation of matrix metalloproteinase-9. Dig Liver Dis 2011;43(1):40–7 doi 10.1016/j.dld.2010.05.013.

40. Williford J-M, Ishihara J, Ishihara A, Mansurov A, Hosseinchi P, Marchell TM, et al. Recruitment of CD103(+) dendritic cells via tumor-targeted chemokine delivery enhances the efficacy of checkpoint inhibitor immunotherapy. Sci Adv 2019;5(12):eaay1357-eaay doi 10.1126/sciadv.aay1357.

41. Nakayama M, Kayagaki N, Yamaguchi N, Okumura K, Yagita H. Involvement of TWEAK in interferon gamma-stimulated monocyte cytotoxicity. J Exp Med 2000;192(9):1373–80 doi 10.1084/jem.192.9.1373.

42. Saitoh T, Nakayama M, Nakano H, Yagita H, Yamamoto N, Yamaoka S. TWEAK induces NF-kappaB2 p100 processing and long lasting NF-kappaB activation. J Biol Chem 2003;278(38):36005–12 doi 10.1074/jbc.M304266200.

43. Taghipour Fard Ardekani M, Malekzadeh M, Hosseini SV, Bordbar E, Doroudchi M, Ghaderi A. Evaluation of Pre-Treatment Serum Levels of IL-7 and GM-CSF in Colorectal Cancer Patients. Int J Mol Cell Med 2014;3(1):27–34.

44. Watkins SK, Egilmez NK, Suttles J, Stout RD. IL-12 rapidly alters the functional profile of tumor-associated and tumor-infiltrating macrophages in vitro and in vivo. J Immunol 2007;178(3):1357–62 doi 10.4049/jimmunol.178.3.1357.

45. Steding CE, Wu ST, Zhang Y, Jeng MH, Elzey BD, Kao C. The role of interleukin-12 on modulating myeloid-derived suppressor cells, increasing overall survival and reducing metastasis. Immunology 2011;133(2):221–38 doi 10.1111/j.1365-2567.2011.03429.x.

46. Allard D, Allard B, Stagg J. On the mechanism of anti-CD39 immune checkpoint therapy. Journal for ImmunoTherapy of Cancer 2020;8(1):e000186 doi 10.1136/jitc-2019-000186.

47. Shaw G, Cavalcante L, Giles FJ, Taylor A. Elraglusib (9-ING-41), a selective small-molecule inhibitor of glycogen synthase kinase-3 beta, reduces expression of immune checkpoint molecules PD-1, TIGIT and LAG-3 and enhances CD8(+) T cell cytolytic killing of melanoma cells. J Hematol Oncol 2022;15(1):134 doi 10.1186/s13045-022-01352-x.

48. Kim SH, Kim K, Kwagh JG, Dicker DT, Herlyn M, Rustgi AK, et al. Death induction by recombinant native TRAIL and its prevention by a caspase 9 inhibitor in primary human esophageal epithelial cells. J Biol Chem 2004;279(38):40044–52 doi 10.1074/jbc.M404541200.

49. Campeau E, Ruhl VE, Rodier F, Smith CL, Rahmberg BL, Fuss JO, et al. A versatile viral system for expression and depletion of proteins in mammalian cells. PLoS One 2009;4(8):e6529 doi 10.1371/journal.pone.0006529.

